# Biosynthesis of cittilins, unusual ribosomally synthesized and post-translationally modified peptides from *Myxococcus xanthus*

**DOI:** 10.1101/2020.05.25.114512

**Authors:** Joachim J. Hug, Jan Dastbaz, Sebastian Adam, Ole Revermann, Jesko Koehnke, Daniel Krug, Rolf Müller

## Abstract

Cittilins are secondary metabolites from myxobacteria comprised of three L-tyrosines and one L-isoleucine forming a bicyclic tetrapeptide scaffold with biaryl and aryl-oxygen-aryl ether bonds. Here we reveal that cittilins belong to the ribosomally synthesized and post-translationally modified peptide (RiPP) family of natural products, for which only the crocagins have been reported from myxobacteria. A 27 amino acid precursor peptide harbors a *C*-terminal four amino acid core peptide, which is enzymatically modified and finally exported to yield cittilins. The small biosynthetic gene cluster responsible for cittilin biosynthesis also encodes a cytochrome P450 enzyme and a methyltransferase, whereas a gene encoding a prolyl endopeptidase for the cleavage of the precursor peptide is located outside of the cittilin biosynthetic gene cluster. We confirm the roles of the biosynthetic genes responsible for the formation of cittilins using targeted gene inactivation and heterologous expression in *Streptomyces*. We also report first steps towards the biochemical characterization of the proposed biosynthetic pathway *in vitro*. An investigation of the cellular uptake properties of cittilin A connected it to a potential biological function as an inhibitor of the prokaryotic carbon storage regulator A (CsrA).

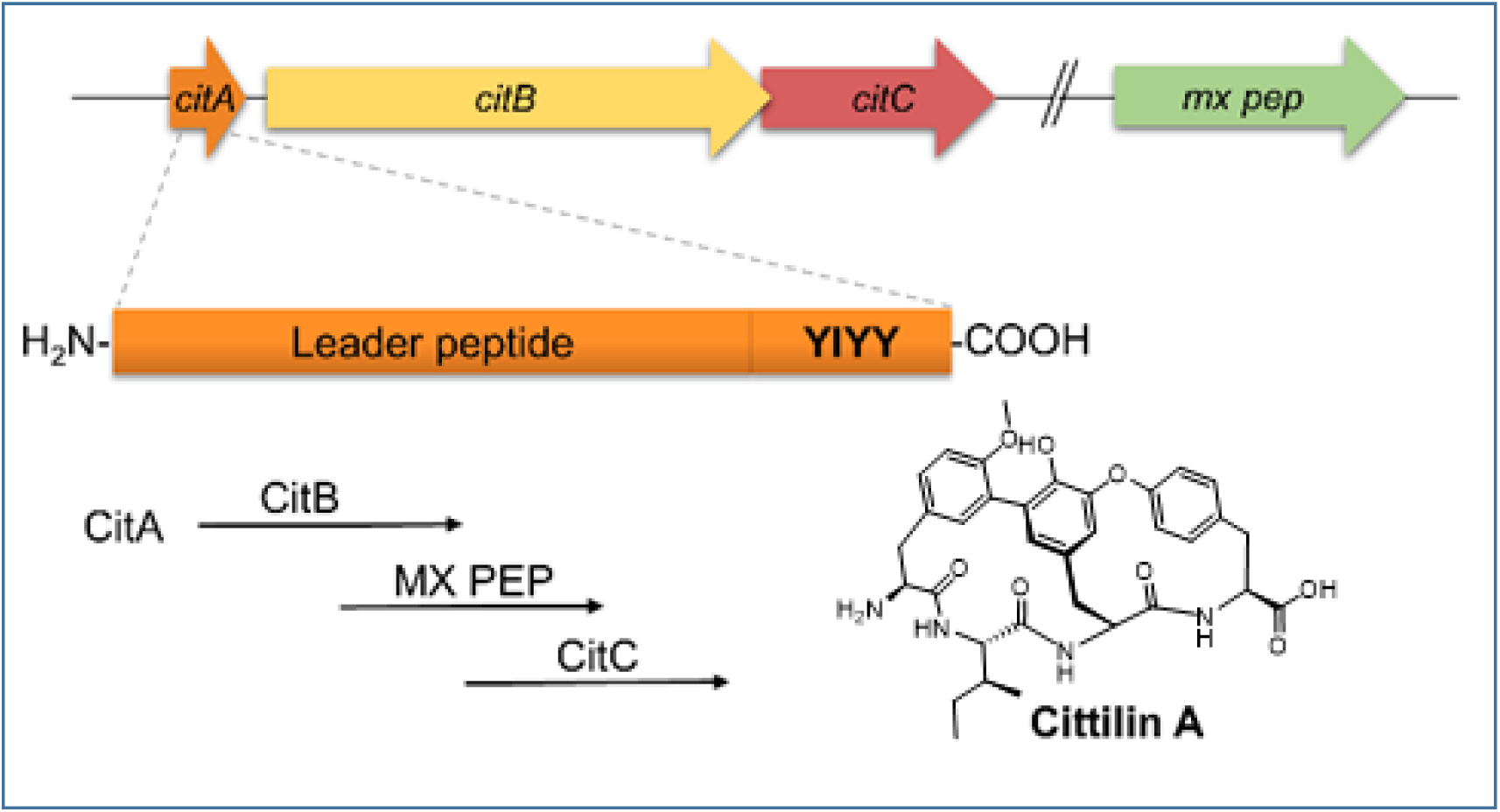

## 1 Introduction

Myxobacteria provide a multitude of natural products that exhibit diverse biological activities, which frequently feature novel modes-of-action *(1)*. Many natural products produced by myxobacteria are derived from large biosynthetic enzyme complexes of the modular non-ribosomal peptide synthetase (NRPS) and polyketide synthase (PKS) types *(2)*. The combinatorial nature of these megasynthetases contributes to the structural diversity of myxobacterial secondary metabolites, and the high mutual similarity of catalytic domains involved in their biosynthesis greatly facilitates the *in silico* identification and characterization of genes encoding these microbial biosynthetic pathways *(3)*. Tools like the “antibiotics and secondary metabolite analysis shell” (antiSMASH) allow the rapid genome-wide identification, annotation and analysis of biosynthetic gene clusters (BGCs) *(4)* and are therefore instrumental for the discovery of new microbial natural products by genome-mining. However, other types of biosynthetic pathways are smaller, or the involved genes are less obviously clustered than the large multi-domain NRPS and PKS genes, making them more difficult to identify in bacterial genomes. An example for such BGCs is the relatively young but steadily expanding family of ribosomally produced and post-translationally modified peptides (RIPPs), which includes the lantibiotics pinensins *(5)* and epidermin *(6)*, lassopeptides such as citrocin *(7)*, the antibiotic bottromycins, which feature a unique macoramidine linkage *(8,9)*, and the marine-derived divamides *(10)*, just to name a few. Recently developed bioinformatics tools allow the automated detection of certain classes of RIPPs in bacterial genomes based on signature genes and canonical recognition elements *(11–16)*. However, the pace of novel RIPPs discovery in recent years suggests that an abundance of yet uncovered RIPP families likely exist. Most of these might until now have evaded identification because their structures and the underlying biosynthetic pathways deviate from previously described patterns. The only myxobacterial RiPP for example that has been characterized biosynthetically are crocagins, which contain a tetrahydropyrrolo [2,3-*b*] indoline core *(17)*.

In this study we reveal cittilins (Figure 1) as a new class of RIPPs and connect them to a small BGC in the myxobacterium *Myxococcus xanthus* DK1622. This strain is a model organism for the study of bacterial motility and multicellular differentiation *(18)* and exemplifies the concept of a myxobacterial multiproducer of secondary metabolites *(19)*. Its genome contains 18 BGCs encoding NRPS, PKS, and NRPS/PKS hybrid systems *(20)*. Cittilins are also produced by *M. xanthus* DK1622, and in fact by more than half of all *M. xanthus* strains analyzed to date *(21)*. Cittilin A and B (Figure 1) were first isolated in the course of the study on the myxobacterial metabolite saframycin in *Myxococcus xanthus* Mx x48 *(22–24)*. In a later screening, cittilin B was isolated independently from *Streptomyces* strain 9738 and therefore named RP-66453 *(25)*. Trowitzsch-Kienast *et al.* elucidated the planar structure of cittilins, whereas the stereochemistry of all five carbonic stereocenters and the configuration of the biaryl-atropisomer was determined via total synthesis *(26–29)*. Stereochemical determination revealed the absolute configuration of all carbonic stereocenters as *S* and the molecule axis has been assigned as *R*. Hence, all four amino acid building blocks are L-configured. The bicyclic structure of cittilins resembles the cross-linked structures of the glycopeptide antibiotics vancomycin, teicoplanin and kistamicin (Figure 1), which are generated via known modular NRPS machineries and a subsequent oxidative cyclization cascade of cytochrome P450 enzymes. These cytochrome P450 enzymes perform stepwise cyclization of the NRPS-bound heptapeptide to generate rigid, active glycopeptide antibiotics *(30)*.

**Figure 1.**
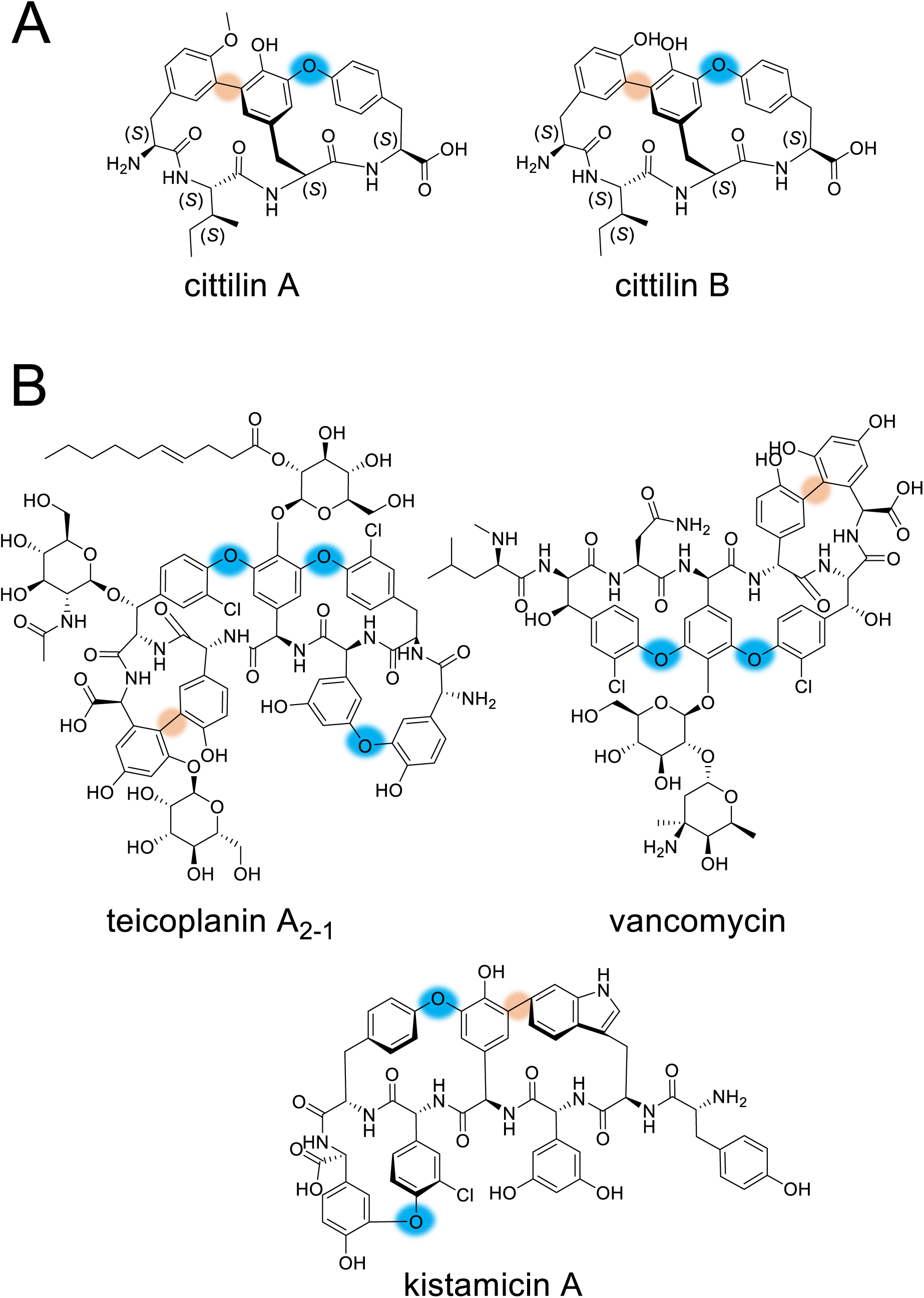
Chemical structures of the biaryl tetrapeptides cittilin A and B (**A**), and glycopeptide antibiotics vancomycin, teicoplanin A_2-1_ and kistamicin (**B**) showing related substructures. Blue circles indicate aryl-oxygen-aryl ether bonds and beige-colored circles indicate biaryl bonds.

In addition to antibiotics, there are also natural products that target bacterial virulence by abolishing pathogenic features without killing the bacterial pathogens *(31)*. The identification of these natural products is more challenging and requires sophisticated screening platforms, but at the same time these screening efforts pave the way for new antimicrobial treatments with less compound-induced selection pressure, leading to reduced rate of resistance development *(32)*. The carbon storage regulator protein A (CsrA) was identified in many Gram-negative bacteria (such as *Helicobacter pylori (32)* and *Salmonella thyphimurium (33)*) to be involved in the regulation and translation of numerous virulence factors by binding the 5’ untranslated region of mRNA. These CsrA-RNA interaction represses the translation of target transcripts through competition with the ribosome for RNA binding and modulates expression of multiple virulence-relevant processes, necessary for successful host infection *(34)*. A test system based on surface plasmon resonance and fluorescence polarization technologies was established and validated to test approx. 1000 small-molecules as potential inhibitors of the CsrA-RNA interaction. This study revealed cittilin A (named in the publication as the undisclosed structure “MM14”) as the most potent inhibitor of the CsrA-RNA interaction of all compounds tested (IC_50_: 4 μM) *(35)*.

We present the identification and heterologous expression of the genes encoding the biosynthetic machinery required to produce cittilin A and B. We delineate the biosynthetic origin of cittilins as RiPPs through targeted gene inactivation experiments and *in vitro* biochemical characterization of involved enzymes. In addition, we provide further biological characterization of cittilin A by investigating its cell entry properties and link it to a potential biological function as an inhibitor of the CsrA *(35)*.

## 2 RESULTS AND DISCUSSION

### Identification of cittilin biosynthesis genes

Cittilin A and B are commonly found in extracts derived from the species *M. xanthus* including the model strain DK1622, as judged on the basis of high-resolution LC-MS data from myxobacterial secondary metabolome studies *(21,36)*. We first verified the structures of cittilin A and B using NMR spectroscopy, since no analytical data for these compounds isolated from myxobacteria was available in the literature *(37)* (**Supporting Information**). Our analysis confirmed that the structure of myxobacterial cittilin B is identical to RP-66453 from *Streptomyces* strain 9738 *(37)* (**Supporting Information**). We then set out to investigate the genome of *M. xanthus* DK1622 for a potential cittilin BGC. However, retrobiosynthetic scrutiny of the *M. xanthus* DK1622 genome sequence based on NRPS biosynthesis did not highlight biosynthetic genes that could plausibly be responsible for the production of cittilins. This *in silico* analysis result agrees with a previous study, where the construction of a targeted mutant library including genetic disruption of nine NRPS-related BGCs in the genome of *M. xanthus* DK1622 did not yield any mutant with abolished production of cittilins *(20)*.

Given the peptide-based chemical structure of cittilins, we decided to explore the hypothesis that cittilin biogenesis could be a hitherto undescribed type of RiPP pathway. Formation of these peptide-derived compounds starts with ribosomal translation to give a precursor peptide, which typically undergoes diverse enzymatic modifications while it matures into the fully decorated natural product. Most often RiPP precursor peptides consist of a leader peptide important for substrate recognition by the modifying enzymes and a core peptide, which undergoes enzymatic modifications. We thus searched the genome of *M. xanthus* DK1622 for a locus encoding a core peptide with the amino acid sequence YIYY. One such genetic locus was found encoding the tetrapeptide YIYY, located at the end of a small 84-base pairs open reading frame (ORF) we named *citA*. Upstream of *citA*, which might encode the precursor peptide CitA, we identified four ABC transporter genes (*citT*_*a*_*–T*_*d*_), while downstream one gene encoding a cytochrome P450-type enzyme (*citB*) and a gene encoding a methyltransferase (*citC*) are located (Figure 2). Since several *M. xanthus* strains can produce cittilins, we investigated several confirmed myxobacterial producers of cittilins with available genome sequence data for the presence of this genetic locus. In fact, the genetic region putatively responsible for the biosynthesis of cittilins was identified in eight strains with available genome sequences (**Supporting Information**). Comparison of these additional BGCs assisted the annotation of the region thought to be responsible for the production of cittilin A and B. Gene cluster comparison revealed that the core region of cittilin biosynthesis is very small and the surrounding genes are highly variable. In fact, only the genes encoding CitA, CitB and CitC are clustered and thus likely constitute the essential cittilin biosynthesis operon.

**Figure 2:**
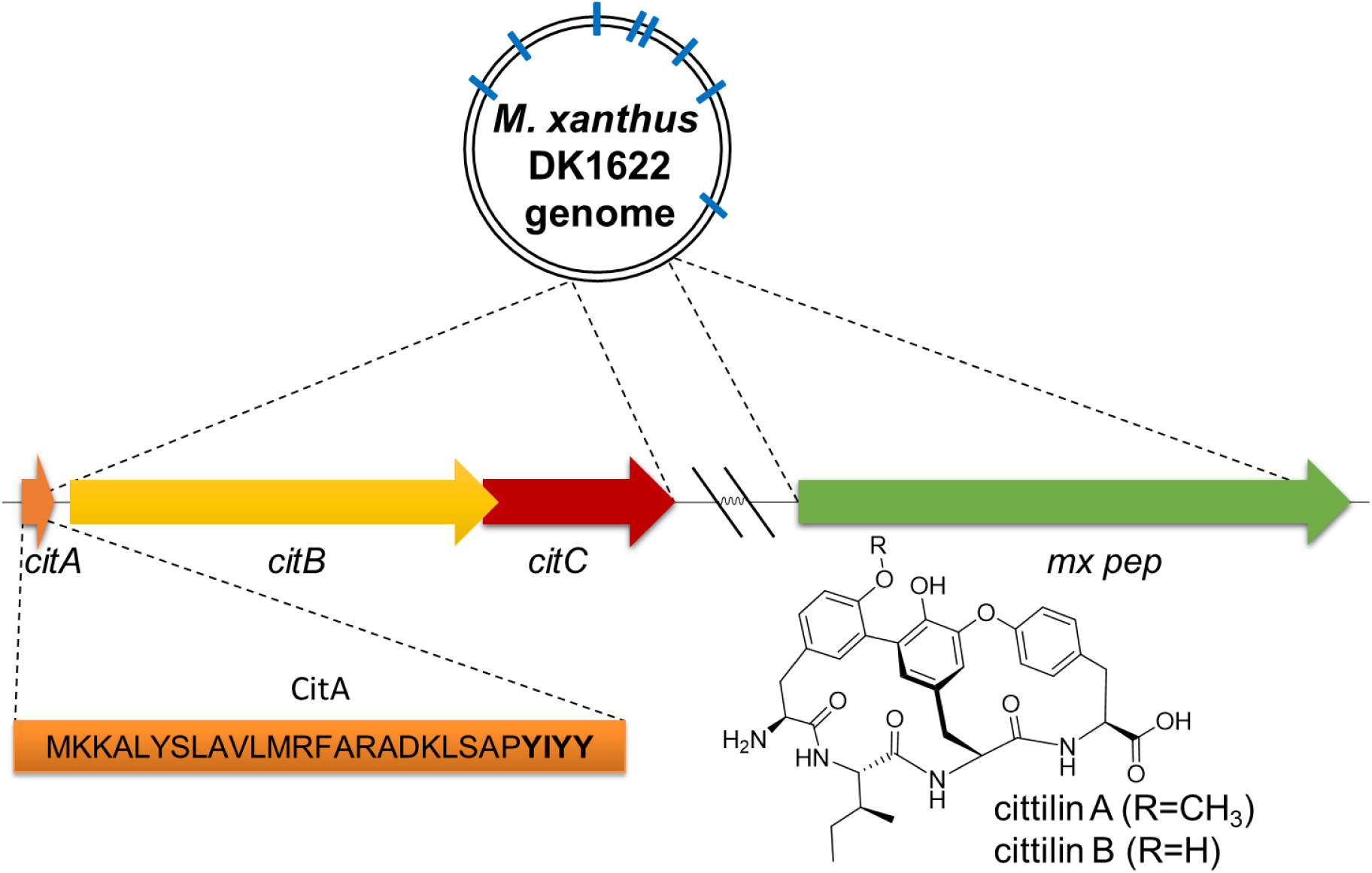
Retrobiosynthetic investigation of cittilin biosynthesis. The chemical structures of cittilin A and B consist of the proteinogenic amino acids L-tyrosine, L-isoleucine, L-tyrosine and L-tyrosine (YIYY). One genetic locus in the genome of *Myxococcus xanthus* DK1622 encodes the tetrapeptide YIYY, located at the end of a small open reading frame. Gene cluster comparison with other cittilin producers confirmed that genes encoding the precursor peptide (*citA*), the cytochrome P450 enzyme (*citB*) and the methyltransferase (*citC*) are strictly conserved and are thus likely to constitute the essential biosynthetic core region.

Next, publicly available genome sequences were investigated to find genes resembling the putative cittilin gene cluster. From the genomes accessible in the GenBank database, six *M. xanthus* strains and *Myxococcus fulvus* HW-1 harbor gene clusters with high similarity to the cittilin gene cluster from *M. xanthus* DK1622; *Myxococcus hansupus mixupus* features the aforementioned biosynthetic organization to afford cittilins but has a nucleotide sequence encoding the deviant core peptide YHYY *(38)* (**Supporting Information**). Furthermore, the genome sequence of *Streptomyces sp.* Ncost-T10-10d contains a gene encoding a putative precursor peptide consisting of 27 amino acids (albeit the core peptide ends with the amino acid sequence YSYY). Similar to the myxobacterial biosynthetic gene cluster organization, this hypothetical cittilin-related operon also contains a gene encoding a CitB homolog. However, different from the myxobacterial cittilin gene cluster architecture, no *citC* homolog was found downstream of the *citB* homolog in *Streptomyces sp.*. This finding is consistent with an apparent difference in decoration of the core scaffold between *Streptomyces* and *Myxococcales*: it appears that *Streptomyces* lacks a methyltransferase, whereas all genome sequence data available for myxobacteria strictly shows the presence of a gene encoding a CitC homolog.

### Cittilin biosynthesis in *M. xanthus* DK1622

To confirm the assignment of cittilins to the identified candidate BGC in the native producer *M. xanthus* DK1622, we conducted gene disruption experiments by single crossover recombination of a plasmid into the putative biosynthetic genes (Figure 3A_1_–A_3_). In addition, a vanillate-inducible promoter system was inserted via single crossover recombination in front of the ABC transporter genes *citT*_*a*_*–citT*_*d*_ or the identified ORF encoding CitA, respectively, to control the expression of the cittilin operon in *M. xanthus* DK1622 through supplementation of vanillate. Genetic disruption of *citB* abolished the production of cittilin A and B. Disruption of *citC* located within the cittilin operon also abolished the production of cittilin A as predicted, whereas the production rate of cittilin B increased significantly (Figure 3B). In contrast, disruption of *citT*_*a*_ intended to abolish tetrameric ABC transporter formation, did not significantly influence the production rate of cittilin A or B in *M. xanthus* DK1622 (Figure 3B). Since the nucleotide sequence length of *citA* was deemed too short for genetic disruption via single crossover recombination, a vanillate-inducible promoter system was inserted upstream of the identified precursor peptide gene *citA*. Subsequent HPLC-MS analysis clearly showed the influence of vanillate-induced expression of *citA* but not *citT*_*a*_ on the production rate of cittilin A and B in the generated mutant (Figure 3C).

**Figure 3:**
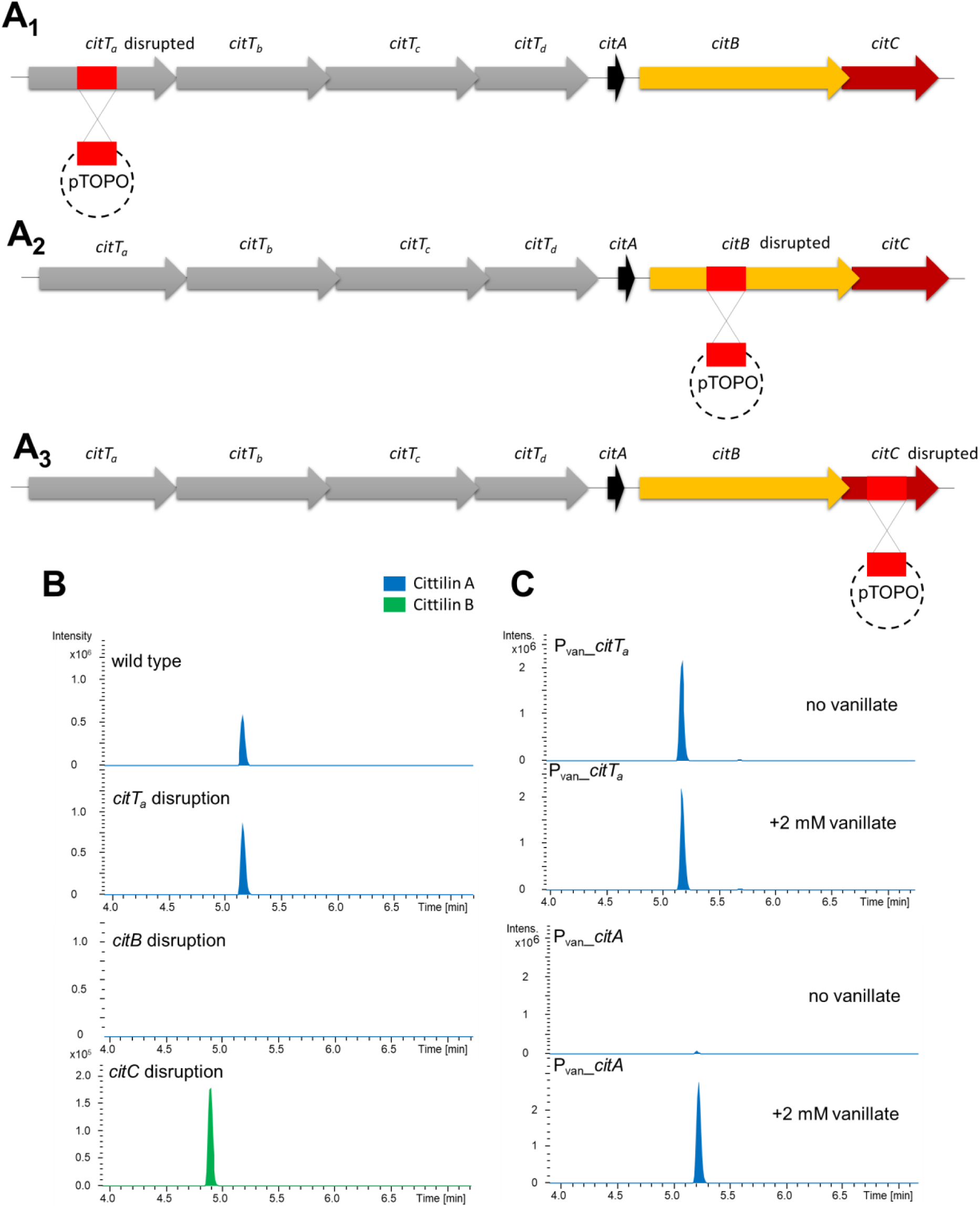
Schematic overview of the cittilin biosynthetic gene cluster identified in *M. xanthus* DK1622, including the ABC transporter genes *citT*_*a*_*–citT*_*d*_, the precursor peptide gene *citA*, the cytochrome P450 enzyme gene *citB* and methyltransferase gene *citC* (**A**_**1**_**–A**_**3**_). Genetic modifications comprise *citT*_*a*_ (**A**_**1**_), *citB* (**A**_**2**_), and *citC* (**A**_**3**_) disruption and vanillate-inducible promoter insertion in front of *citT*_*a*_ and *citA* (not shown). **B**) HPLC-MS EIC of crudes extracts from *M. xanthus* DK1622 wild type and the respective gene disruption mutant. **C**) HPLC-MS EIC of crude extracts from *M. xanthus* DK1622_P_van__*citT*_*a*_ and *M. xanthus* DK1622_P_van__*citA* with and without supplementation of 2 mM vanillate. EIC: Extracted ion chromatogram, blue: 631.2768 m/z, with a width of 7.9 ppm, cittilin A [M+H]; green: 617.2611 m/z, with a width of 7.9 ppm, cittilin B [M+H].

These results confirmed the proposed cittilin biosynthetic locus and the operon structure of the essential biosynthetic core region, consisting of the precursor peptide, the cytochrome P450 enzyme and the methyltransferase encoded by *citA, citB* and *citC*, respectively. Nevertheless, at least one additional enzyme was likely to be involved in the biosynthesis of the cittilins. Rather surprisingly, the cittilin operon does not contain a gene encoding a peptidase that is commonly found in RiPP biosynthetic gene clusters and is usually required to catalyze the cleavage of the leader peptide and thus releases the modified core peptide *(39)*. Therefore, we attempted to define which enzymes are required for the biosynthesis of cittilins via heterologous expression of the biosynthetic genes encoded in the cittilin BGC.

### Heterologous expression of cittilin BGC

To confirm the functions of the identified gene products for the production of cittilins, we aimed to perform heterologous expression of candidate genes in the distantly related order of *Streptomyces*. Using a streptomycete as the heterologous host to produce cittilin appeared promising, since cittilin B is known from *Streptomyces* strain 9738 *(25)* and we thus reasoned that the streptomycete host should be compatible with cittilin production. Nevertheless, one should emphasize that previous attempts towards heterologous expression of myxobacterial gene clusters in streptomycetes were met with little success *(40)*. The heterologous production of epothilone *(41)* and soraphen *(42)* were described, albeit at very low yields.

The operon consisting of *citA, citB* and *citC* was amplified by PCR and subcloned into the expression vector based on plasmid pSET152 (**Supporting Information**). Following transformation into *Streptomyces albus* del14 *(43)* by conjugation, this initial heterologous expression experiment did yield cittilin A at very low concentrations according to LC-MS analysis of the exconjugants (Figure 4A). As mentioned above, the identified genetic locus for production of cittilin A and B lacks an ORF for a peptidase that removes the leader peptide during a typical RiPP biosynthesis *(44)*. Since L-proline is located in front of the core peptide amino acid sequence YIYY, one would expect a prolyl endopeptidase to catalyze the cleavage of the precursor peptide. Among the numerous peptidase genes present in the genome sequence of *M. xanthus* DK 1622, one specific prolyl endopeptidase gene (*mx pep*) had been investigated previously *(45)*. Since MX PEP showed a higher preference for cleavage of Pro-Tyr/Phe bonds over Pro-Gln sites *(46,47)* and no homolog of *mx pep* was present in the genome of the heterologous host, we postulated that MX PEP could be involved in the formation of cittilins. We thus constructed a fused cittilin operon that included the prolyl endopeptidase MX PEP. This operon yielded higher production of cittilin A (Figure 4B) in the heterologous host *S. albus* del14 *(43)*. The corresponding LC-MS results readily showed the production of cittilin A in the heterologous host *S. albus* del14, and the identity of heterologously produced cittilin A was unambiguously confirmed through tandem MS (MS/MS) measurements, and by evaluation of fragment masses in the obtained MS/MS spectra compared to the purified cittilin A as reference (**Supporting Information**).

**Figure 4:**
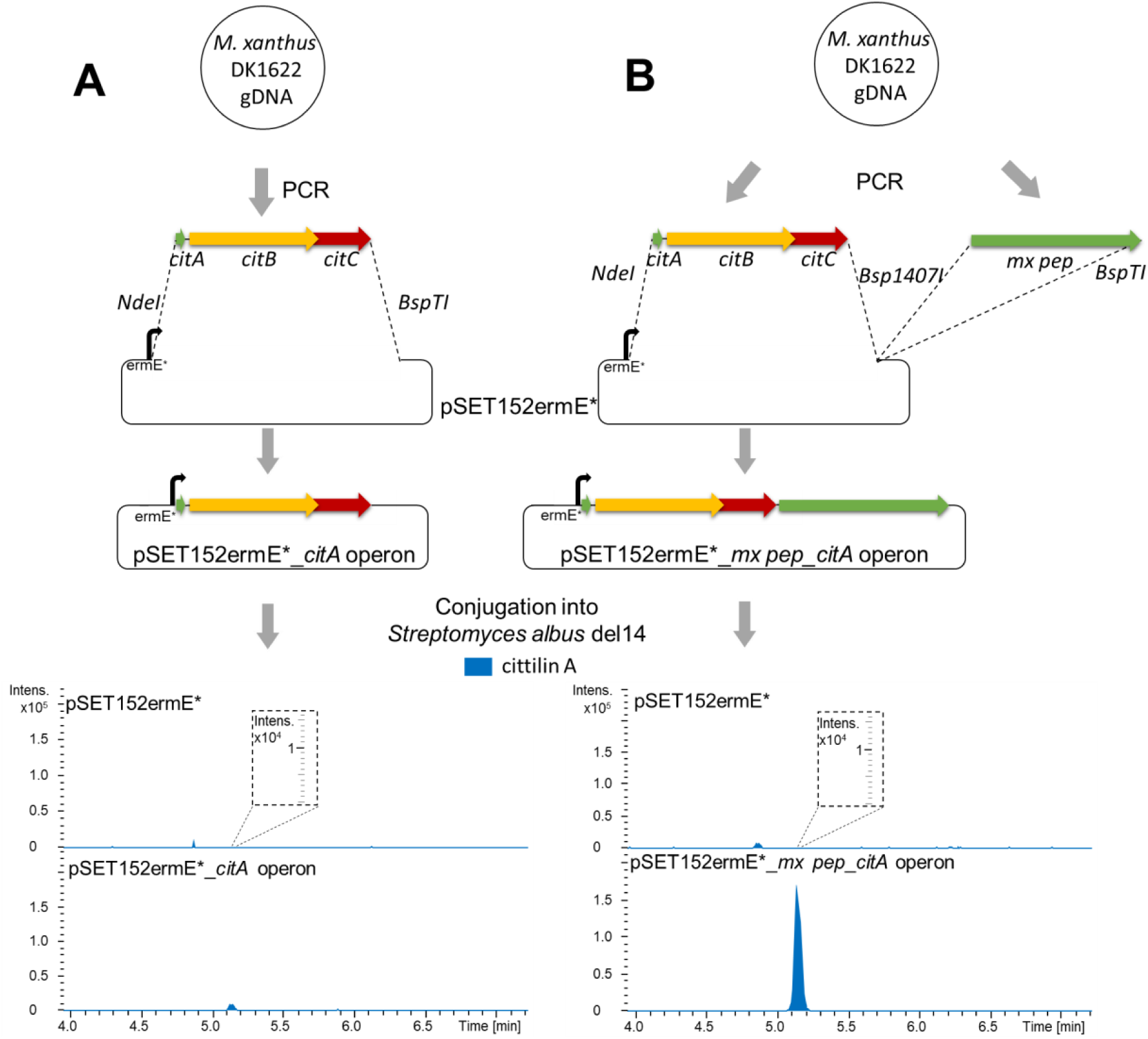
Heterologous production of cittilin A. (**A**) Heterologous expression of the cittilin operon *citA*–*citC (citA* operon) based on the constructed expression vector pSET152 resulted in low production of cittilin A. (**B**) Heterologous expression of *citA* operon and the gene encoding a previously characterized prolyl endopeptidase MX PEP, increased the amount of produced cittilin A. EIC: Extracted ion chromatogram, blue: 631.2768 m/z, with a width of 7.9 ppm, cittilin A [M+H].

Following these promising results from the heterologous expression platform, we sought to verify the influence of MX PEP on the production of cittilins in the native producer *M. xanthus* DK1622. However, to our surprise genetic disruption of the *mx pep* gene in *M. xanthus* DK1622 did not significantly influence the production rates of cittilin A or B. The fact that MX PEP helps the heterologous host to cleave off the leader peptide is not unexpected, but also not an unambiguous proof that this myxobacterial prolyl endopeptidase is in fact the only protease involved in the cleavage of CitA in the myxobacterial host.

### *In vitro* investigation of cittilin biosynthesis

The successful heterologous expression of cittilin genes provided the conceptual proof for the enzymatic steps needed for cittilin biosynthesis. Next, we aimed to shed light on the sequence of enzymatic steps leading to cittilin formation. Following the conventional biosynthetic logic of RiPP pathways *(39)*, the overall process can be subdivided into four steps: 1) production of the ribosomal precursor peptide, 2) posttranslational modification of the core peptide, usually dependent on leader peptide recognition (primary modifications), 3) proteolysis of the leader peptide to yield the modified core peptide and 4) further modifications (also termed secondary modifications) and export of the mature peptide-derived natural product. In the case of cittilin, following the biosynthesis of the 27 amino acid precursor peptide CitA, the next step would involve the catalysis of the biaryl and aryl-oxygen-aryl linkage reactions by the cittilin cytochrome P450 enzyme CitB. From the heterologous expression experiments, it was concluded that CitB alone is likely to catalyze both of these intriguing biosynthetic transformations.

Heterologous production of recombinant CitB in *E. coli* could only be achieved through co-expression of the heat shock chaperone system *groEL*/*groES* in low yields, despite serious efforts (using different strains, genetic constructs and expression conditions, see **Supporting Information**). Production of the CitB homolog (CitB_MCy9171_) from MCy9171 (a confirmed alternative producer of cittilin A and B, see **Supporting Information**)) was ultimately achieved in higher yields and without chaperone gene co-expression in the heterologous host *Streptomyces coelicolor* CH999. However, the carbon monoxide (CO)-spectral analysis of this homolog after purification shows an absorption maximum at 420 nm. The spectral peak at 420 nm for the ferrous (Fe^2+^) CitB_MCy9171_-CO complex indicates denaturation of cytochrome P450 enzyme in the process of isolation *(48)*. Accordingly, during catalytic activity testing of recombinantly produced CitB_MCy9171_ in various experiments (including different co-factors and the Fdx/FdR reductase pair system from *Spinacia oleracea*, see **Supporting Information**) no obvious conversion of the precursor peptide was observed. These results, combined with the observation that CitB_DK1622_ could only be obtained in *E. coli* in the presence of co-produced chaperone system GroEL/GroES, hint towards improperly folded protein.

As a workaround and to link CitB to the conversion of the precursor peptide *in vitro*, cell-free lysates of induced *S. coelicolor* CH999, containing different expression plasmids were incubated with chemically synthesized CitA (**Supporting Information**). These cell-free lysate reactions displayed consumption of the supplemented precursor peptide only for cell-free lysate originating from *S. coelicolor* CH999 expressing CitB_MCy9171_, but not the negative controls (**Error! Reference source not found.**A). Supplying the core peptide YIYY instead of full-length CitA did also not result in peptide conversion, which suggests that the leader peptide is essential for processing. Next, we simultaneously supplemented recombinant MX PEP and precursor peptide to cell free lysate from CitB_MCy9171_-producing *S. coelicolor* CH999 and observed the formation of cittilin B (**Error! Reference source not found.**B).

The methyltransferase CitC was suspected to catalyze the last modification step in the biosynthesis, since the native host is naturally producing both, cittilin A and B. In addition, genetic disruption of *citC* resulted in the exclusive production of cittilin B (see above). We could also observe the CitC-dependent methylation of cittilin B with the expected mass shift of +14 Da yielding cittilin A, when a concentrated crude extract (containing cittilin B, but not cittilin A) of a *M. xanthus* DK1622 mutant with *citC* disruption was supplemented with recombinantly produced CitC (Figure 6). In contrast, recombinantly produced CitC showed no methylation activity when incubated with CitA or unmodified core peptide (YIYY) in *in vitro* assays (**Supporting Information**).

**Figure 5:**
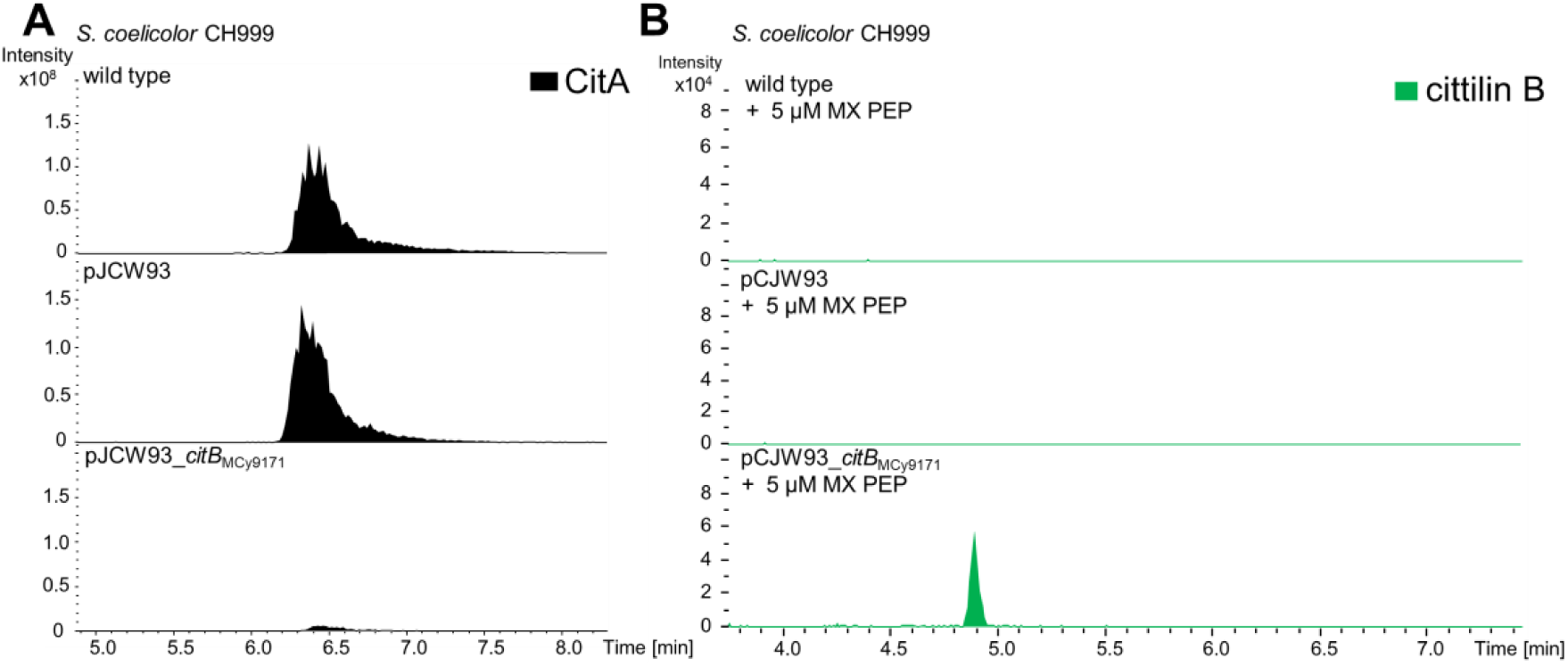
Investigation of cittilin biosynthesis through cell free lysate reactions in the heterologous host *Streptomyces coelicolor* CH999 (**A**) HPLC-MS EIC (3051.75 m/z, chemically synthesized precursor peptide [M+H]) of cell-free lysates of *S. coelicolor* CH999, *S. coelicolor* CH999 with pCJW93 and *S. coelicolor* CH999 with pCJW93_*citB*_MCy9171_ are shown to observe conversion of supplemented chemically synthesized precursor peptide (62.5 µM). Only the cell-free lysate of *S. coelicolor* CH999 with pCJW93_*citB*_MCy9171_ shows significant consumption of the supplemented precursor peptide. (**B**) HPLC-MS EIC (617.2611 m/z cittilin B [M+H]) of cell-free lysates of *S. coelicolor* CH999, *S. coelicolor* CH999 with pCJW93 and *S. coelicolor* CH999 with pCJW93_*citB*_MCy9171_ are shown to observe conversion of supplemented chemically synthesized precursor peptide (62.5 µM) in the presence of recombinantly produced prolyl endopeptidase MX PEP (5 µM). Only the cell-free lysate of *S. coelicolor* CH999 with pCJW93_*citB*_MCy9171_ with added recombinant MX PEP shows production of cittilin B.

**Figure 6:**
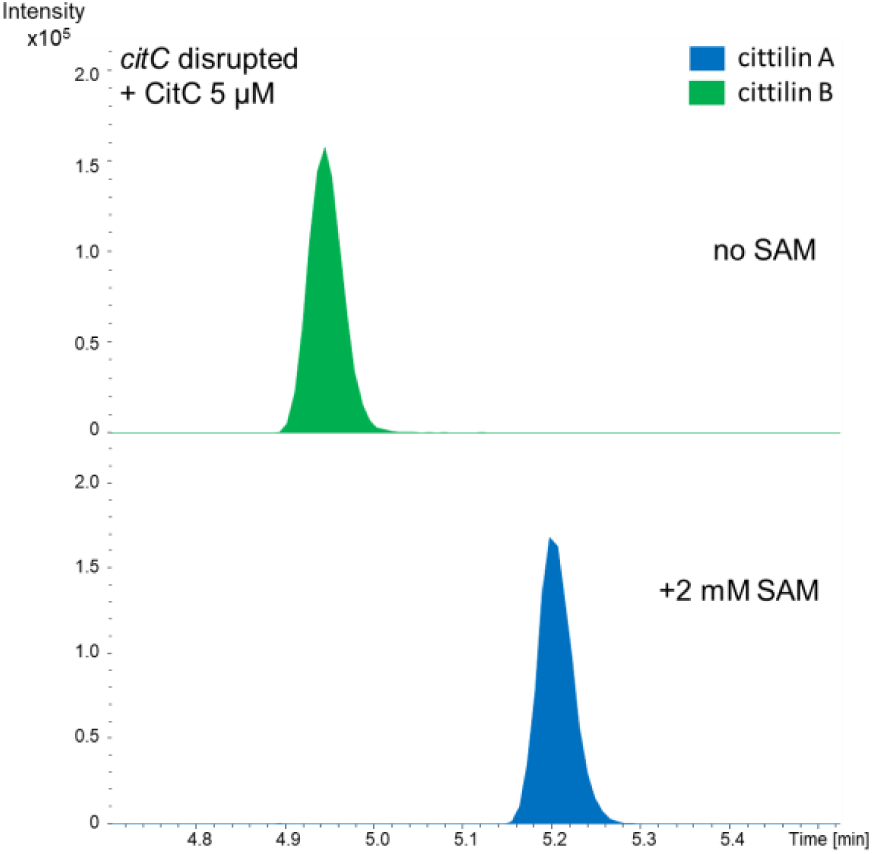
*In vitro* conversion of cittilin B to cittilin A through recombinantly produced methyltransferase CitC. HPLC-MS EIC shows the catalytic conversion of cittilin B to cittilin A via CitC in dependence of *S*-adenosyl-L-methionine (SAM) as a co-factor. EIC: Extracted ion chromatogram, blue: 631.2768 m/z, with a width of 7.9 ppm, cittilin A [M+H]; green: 617.2611 m/z, with a width of 7.9 ppm, cittilin B [M+H].

We conclude that the first enzyme to act on CitA is CitB, which catalyzes both, the biaryl and aryl-oxygen-aryl bond formation. The modified CitA is proteolytically processed by MX PEP, which removes the leader peptide from the modified core peptide (cittilin B). Finally, the cittilin methyltransferase CitC specifically methylates cittilin B, to yield the major derivative cittilin A (Figure 7). In conclusion, we confirmed the proposed biosynthetic pathway as shown in Figure 7 through cell-free lysate experiments for the catalytic investigation of CitB and *in vitro* experiments of CitC and MX PEP. These findings set the stage for further in-depth biochemical analysis; in particular, the formation of the bicyclic ring system in cittilin catalyzed by CitB should become the subject of future studies.

**Figure 7:**
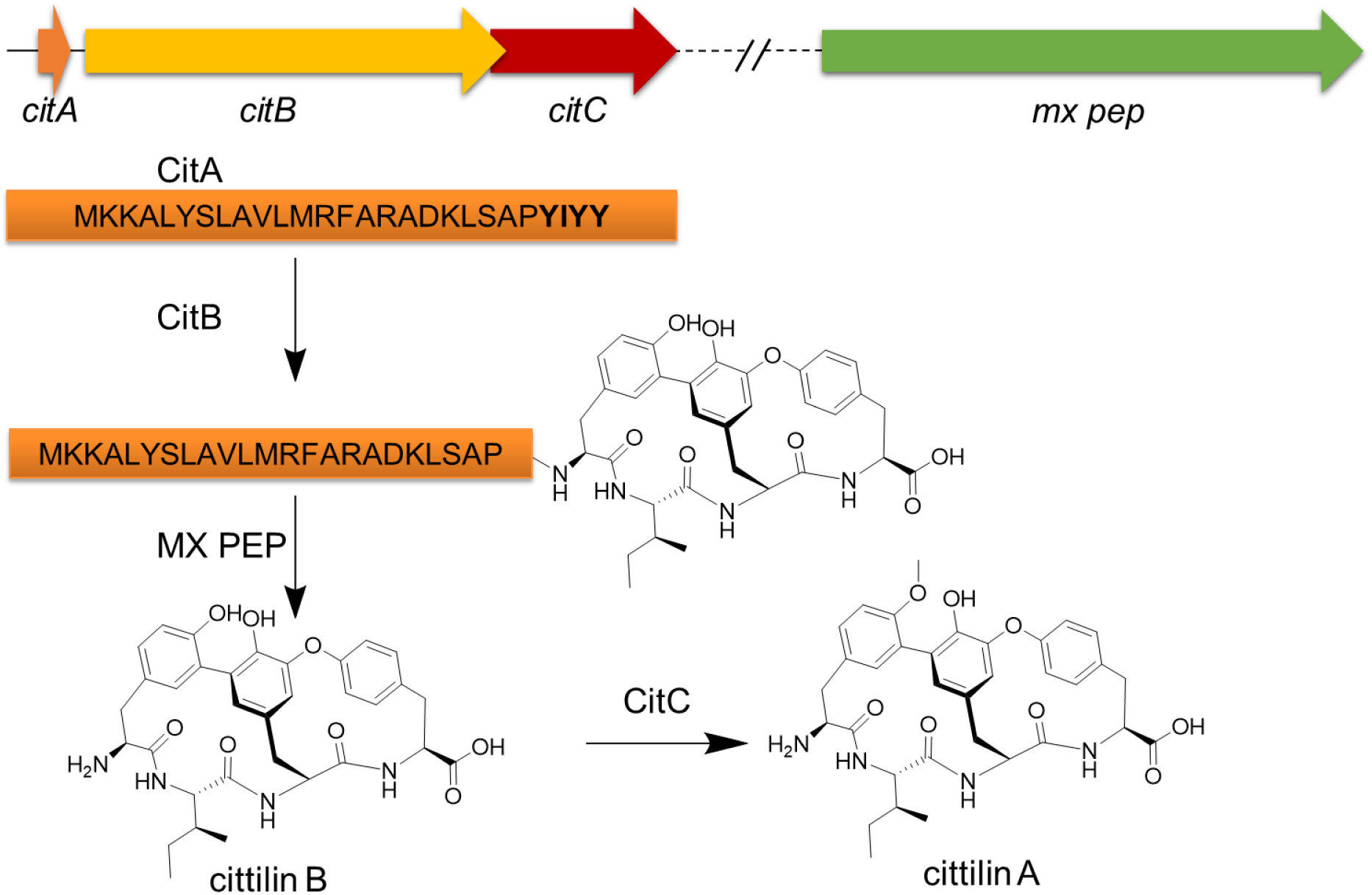
Schematic overview of the cittilin gene operon (consisting of precursor peptide gene *citA*, cytochrome P450 enzyme gene *citB* and methyltransferase gene *citC*) and the distantly located prolyl endopeptidase gene *mx pep*, which likely contributes to formation of cittilin A and B in *M. xanthus* DK1622. The proposed biosynthesis of cittilin A starts with the ribosomally synthesized precursor peptide CitA, which is then further processed by the cytochrome P450 enzyme CitB. Subsequently, the modified precursor peptide (hypothetical chemical structure of the cyclized tetrapeptide is shown) is cleaved by the action of a prolyl endopeptidase (MX PEP being a candidate according to heterologous expression results), which yields cittilin B. Finally, cittilin B is undergoing methylation via the methyltransferase CitC yielding the major myxobacterial compound cittilin A.

### Biological function of cittilin A

Since the production of cittilin occurs widespread among strains of the genus *Myxococcus (21)*, the question of the biological function of this natural product arises. Cittilin A appears to serve none of the biological functions often assigned to other myxobacterial natural products since no significant antibiotic, antifungal or cytotoxic activity was observed (**Supporting Information**) *(49)*. Relevant biological functions of cittilin A were previously listed in the literature as a moderate inhibitor of pancreatic elastase, whereas cittilin B was considered as a neurotensin inhibitor (IC_50_: 30 µg mL^-1^) *(25)*, which even led to synthesis efforts towards the drug development of peptides resembling the tetrapeptidic core structure of cittilin *(50)*.

In a recently conducted screen for inhibitors of CsrA, cittilin A was found to be the most potent inhibitor *(35)*. Cittilin A (named in the publication as the undisclosed structure “MM14”) featured the highest *in vitro* affinity for inhibition of the interaction of CsrA with RNA (IC_50_: 4 μM) *(35)*. Cittilin A was consequently considered as a potential small-molecule inhibitor of this essential virulence factor in opportunistic pathogens. The obvious discrepancy of cittilin A between its potent *in vitro* inhibition of CsrA and the lack of any biological effect against Gram-negative pathogens, implies that pharmacokinetic obstacles account for the absence of its *in cellulo* activity. Therefore, we chose to investigate the cellular uptake of cittilin A with the help of LC-MS analysis as well as fluorescence microscopy. Cell entry of synthesized rhodamine-tagged cittilin A was tested in *E. coli* by fluorescence microscopy of treated bacterial cells in comparison with free rhodamine. Microscopic analysis reveals that *E. coli* cells incubated with free rhodamine display strong fluorescence, whereas cultures treated with rhodamine-tagged cittilin A show significantly less fluorescence (Figure 8). Furthermore, the supernatant of *E. coli* TolC mutants treated with either cittilin A or free rhodamine were extracted separately and subjected to LC-MS analysis to compare cell entry properties of both compounds (or strong adhesion to the cell membrane or cell wall respectively). Only free rhodamine was taken up by the cells, according LC-MS analysis (**Supporting Information**). Given these results, we conclude that cittilin A has poor cell entry capabilities for *E. coli*, which impedes the inhibition of intracellular CsrA and may explain why cittilin A showed no inhibition of pyocyanin production when administered to *Pseudomonas aeruginosa*, an assay commonly used to evaluate anti-virulence efficiency of small-molecules (**Supporting Information**) *(51)*. To support the assumption of poor pharmacokinetics, studies using cittilin A with improved cell entry properties through coupling with iron chelators might give further insights into its biological function. At present we cannot exclude that the biological function of cittilins is distinct from its potential activity as an anti-infective pathoblocker *(52)*.

**Figure 8:**
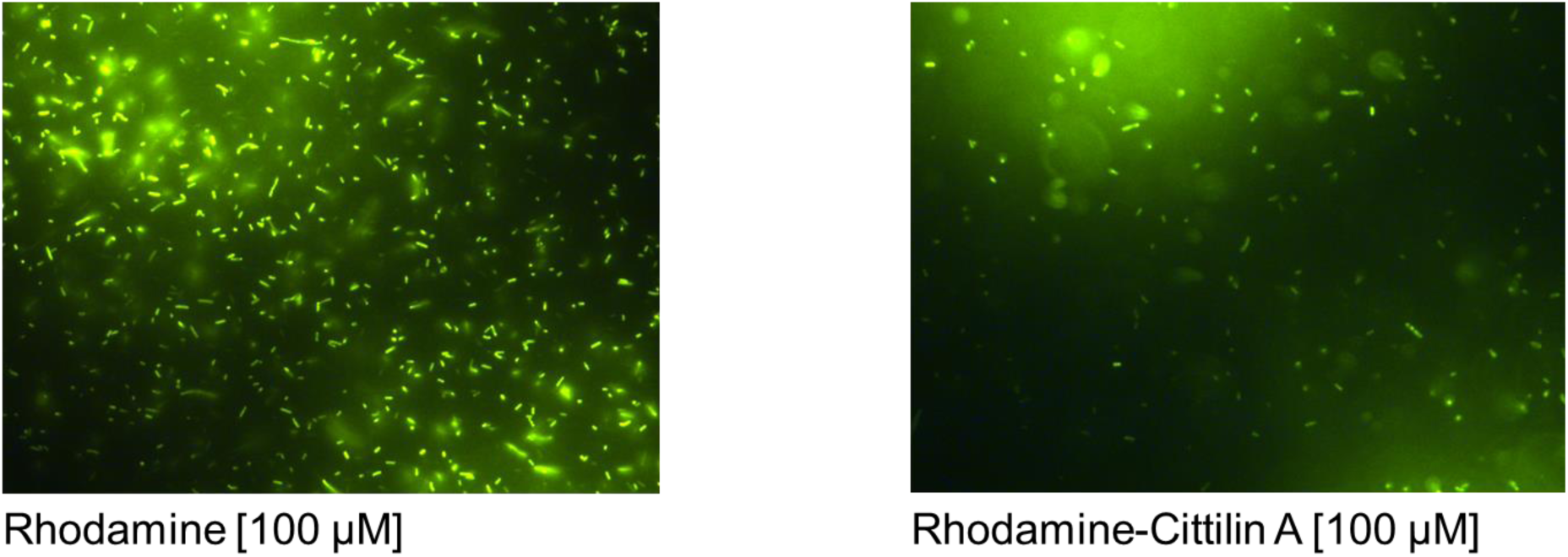
Test for bacterial cell entry of rhodamine (left) and rhodamine-tagged cittilin A (right) in the Gram-negative bacterium *Escherichia coli* (TolC efflux deficient *E. coli* mutant, internal strain collection *(53)*, **Supporting Information**).

## 3 CONCLUSION

We established the molecular basis for the biosynthesis of cittilin A and B, two common natural products of myxobacteria from the genus *Myxococcus*. Even though the rare bicyclic structure of cittilins resembles those of the cross-linked substructures known from the glycopeptide antibiotics vancomycin, teicoplanin and kistamicin, their biosynthesis and biological function deviates significantly. The small BGC involved in the biosynthesis of the cittilins consists of a gene encoding the 27 amino acid precursor peptide CitA, the cytochrome P450 enzyme CitB and the methyltransferase CitC, which is unprecedented for RiPPs. Although the recently described darobactin genetic locus remotely resembles the cittilin BGC, the precursor peptide of darobactin is twice the size of the cittilin precursor peptide and the proposed radical *S*-adenosyl-L-methionine (SAM)-catalyzed cyclization differs significantly from the biaryl and aryl-oxygen-aryl formation in cittilin biosynthesis *(54)*. Strikingly, the formation of the bicyclic ring system in cittilin apparently requires the enzymatic activity of only one cytochrome P450 enzyme. These features are not shared by any of the known RiPP compound classes *(39)* and lead us to the conclusion that cittilins constitute a new class of RiPPs. Our study provides the foundation for further investigation of this small, but intriguing biosynthetic pathway. *In silico* analysis based on publicly available genome sequences already hints towards proposed cittilin derivatives, which feature the alternative core peptide sequences YHYY and YSYY in *M. hansupus mixupus* and *Streptomyces sp.* Ncost-T10-10d, respectively.

The biotechnological generation of bicyclic tetrapeptides resembling the cittilin scaffold are now conceivable both in the native and the heterologous host, but also further investigation of the underlying mechanism of the interesting cytochrome P450 enzyme reaction yielding the bicyclic linkage will be feasible once reaction conditions to establish the respective steps purely *in vitro* are established. In addition, identification of the biosynthetic genes producing putative cittilin derivatives across phylogenetic borders might further help to determine the biological function of these intriguing natural products.

## 4 METHODS

Details of experimental procedures are provided in the **Supporting Information**.

## 5 ASSOCIATED CONTENT

Supporting Information Available: This material is available free of charge via the Internet. The sequence of the cittilin biosynthetic gene cluster originating from *M. xanthus* DK1622 has been deposited in the Minimum Information about a Biosynthetic Gene cluster (MIBiG) database under the accession number BGC0002043.

## 6 AUTHOR INFORMATION

### Notes

The authors declare no competing interests.

## 7 ACKNOWLEDGMENTS

The authors thank W. Hofer for the synthesis of the rhodamine-coupled cittilin A and recording the NMR spectra of the cittilins, J. Herrmann and A. Boese for microscopic fluorescence imaging, S. Schmidt for performing bioactivity assays, S. Amann for conducting the pyocyanin assay, A. Abdulmughni for CO-spectral analysis and N. Zaburannyi for bioinformatic support. J. Hug acknowledges funding by a PhD fellowship of the Boehringer Ingelheim Fonds. Research in R. Müller’s laboratory is funded by the Deutsche Forschungsgemeinschaft (DFG) and the Bundesministerium für Bildung und Forschung (BMBF) and the Deutsches Zentrum für Infektionsforschung Standort Hannover-Braunschweig.

**Notes**

The authors declare no competing interests.

