## Supporting Information for "Biosynthesis of cittilins, unusual ribosomally synthesized and post-translationally modified peptides from *Myxococcus xanthus*"

bioRχiv

### 1 General materials and methods

#### Applied software, sequence analysis and bioinformatics methods

Geneious prime (Biomatters Ltd., Auckland, New Zealand) was used to design primers for PCR and sequencing, to create plasmid maps, to find open reading frames (ORF) and to predict molecular weights of proteins. Furthermore all pairwise and multiple alignments of nucleotide or amino acid sequences where performed with the plugin software from Geneious by using the MUSCLE (Multiple Sequence Comparison by Log- Expectation) alignment (3.8.425 by Robert C. Edgar) since it claims to achieve higher average accuracy and better speed than ClustalW2 or T-Coffee algorithm *(1)*. In order to find homologous genes or proteins, either the nucleotide or amino acid sequence of interest was aligned with the basic local alignment search tool (BLAST) against our in-house genome database or the publically available nucleotide database. Raw data from alignments for *in silico* evaluation of cittilin biosynthetic gene clusters (BGCs) are deposited on our in-house server. The functional prediction of ORFs was performed by either using protein blast and/or blastx program *(2)*. The in-house standard extract database embedded in the software bundle Mxbase Explorer 3.2.27 *(3)* was used for the search of alternative producers of cittilins. The molecular formula and experimentally determined retention times of ions typically observed from cittilins were used as data input. The BGC sequences of confirmed producers of cittilin deriving from our in-house database have been deposited in GenBank and are accessible under the accession number as displayed in **Tab. S1**. In addition, the sequence of the cittilin BGC originating from *Myxococcus xanthus* DK1622 has been deposited in the Minimum Information about a Biosynthetic Gene cluster (MIBiG) database under the accession number BGC0002043

**Tab. S1** In-house genome sequence analysis of myxobacterial strains

| Cittilin A producing strain | Reference | Accession number BGC |
|---|---|---|
| *M. xanthus* Mxx48 (MCy8278) | *(4)* | MN731370 |
| *Archangium violaceus* Cb vi35 (MCy8337) | *(5)* | MN731363 |
| *Archangium gephyra* (MCy8375) | *(6)* | MN731362 |
| *Myxococcus fulvus* (MCy10608) | *(7)* | MN731365 |
| *Myxococcus xanthus* DK897 (MCy8986) | *(8)* | MN731369 |
| *Myxococcus xanthus* DK1622 (MCy9151) | *(9)* | MN731368 |
| *Myxococcus fulvus* strain ATCC BAA-855/HW-1 (MCy11108) | *(10)* | NC_015711.1 |
| *Myxococcus virescens* ST200611 (MCy11474) | *(11)* | MN731367 |
| *Myxococcus sp.* MCy9171 | *(12)* | MN731366 |
| *Myxococcus fulvus* MCy8286 (Mxf65) | *(13)* | MN731364 |

### Maintenance of bacterial cultures, molecular cloning and construction of plasmids *(14)*

Routine handling of nucleic acids, such as isolation of plasmid DNA, restriction endonuclease digestions, DNA ligations, and other DNA manipulations, was performed according to the standard protocols *(15)*. *Escherichia coli* HS996 (Invitrogen) was used as host for standard cloning experiments and *E. coli* SCS110 (Stratagene) for preparation of plasmid DNA free of Dam or Dcm methylation. *E. coli* strains were cultured in LB liquid medium or on LB agar (1% tryptone, 0.5% yeast extract, 0.5% NaCl, (1.5% agar) at 30–37 °C and 200 rpm) overnight (o/n). Antibiotics were used at the following final concentrations: 100 µg/mL ampicillin, 50 µg/mL kanamycin, 12 µg/mL oxytetracycline, 50 µg/mL apramycin, 25 µg/mL chloramphenicol and 25 µg/mL nalidixic acid. Transformation of *E. coli* strains was achieved via electroporation in 0.1 cm wide cuvettes at 1250 V, a resistance of 200 Ω, and a capacity of 25 µF. Plasmids were purified either by standard alkaline lysis *(15)* or by using the GeneJet Plasmid Miniprep Kit (Thermo Fisher Scientific™) or the NucleoBond PC100 kit (Macherey-Nagel). Restriction endonucleases, alkaline phosphatase (FastAP) and T4 DNA ligase were purchased from Thermo Fisher Scientific™. Oligonucleotides used for PCR and sequencing were acquired from Sigma-Aldrich and are listed in **Tab. S2 and Tab. S3**. PCRs were carried out in a Mastercycler® pro (Eppendorf) using Phusion™ High-Fidelity according to the manufacturer's protocol. Temperature and duration setting for each thermocycling step in PCR with Phusion™ High-Fidelity polymerase were performed as follows: Initial denaturation (30 s, 98 °C); 33 cycles of denaturation (15 s, 98 °C), annealing (15 s, 53–72 °C, depending on the melting temperature of primers) and elongation (based on PCR product length 30 s/1 kb, 72 °C); and final extension (10 min, 72 °C). When amplification of desired PCR product could not be achieved according to the manufacturer's instructions, additives like DMSO (3–8%), glycerol (4–8%) and betaine (0.5 mM) were supplemented. PCR products or DNA fragments from restriction digestions were purified by agarose gel electrophoresis and isolated using the PCR clean-up gel extraction kit using Nucleo Spin® (Macherey-Nagel). To recover DNA fragments larger than 8 kb, the Agarose Gel DNA Extraction Kit from Roche was used. After selection with suitable antibiotics, clones harboring correct ligation products were identified by plasmid isolation and restriction analysis with a set of different restriction endonucleases. In addition to restriction analysis, integrity of the constructs for induced gene expression was verified by sequencing.

##### *Streptomyces* conjugation procedure

Conjugation procedure between *E. coli* and *Streptomyces* was performed following a procedure adapted from Mazodier et al. *(16)*. For conjugation SM-agar plates were used. The plates were prepared one week before in order to dry the plates. *E. coli* ET12567 harboring pUZ8002 was transformed with the respective plasmid/genetic construct for heterologous production of cittilin in *Streptomycs albus* del14 and *Streptomyces coelicolor* CH999 for recombinant protein production (cittilin cytochrome P450). O/n cultures of *E. coli* ET12567 + pUZ8002 harboring the genetic construct of interest were prepared: 5 mL of 2TY medium were inoculated with the respective *E. coli* strains from a cryogenic long term stock with 25 µg/mL chloramphenicol, 50 µg/mL kanamycin and 50 µg/mL apramycin. The next day, the o/n cultures were used with 0.75 mL, 1.00 mL and 1.25 mL as inoculation volume to inoculate 20 mL of 2TY medium with same antibiotics and concentrations for (o/n) cultures. When $OD_{600}$ reached 0.4, the cultures were centrifuged for 10 min at 4000 rpm. The supernatant (SN) was discarded and the cell pellet (CP) was re-suspended in 10 mL 2TY medium. Applying same centrifugation parameters, the cells were centrifuged, the SN was discarded and the CP was washed with 2TY medium again. After centrifugation according the same parameters as before, the SN was discarded and the CP was re-suspended in 1 mL 2TY medium and stored on ice.

Spores of *S. coelicolor* CH999 or *S. albus* del14, which have been aliquoted and frozen at -80 °C were thawed on ice. The spore aliquots of about 100 µL were washed with 1 mL 2TY medium by pipetting up and down and centrifuged for 10 min at 6000 rpm. A second washing procedure with 1 mL of 2TY medium followed by centrifugation for 2 min at 6000 rpm was performed. After discarding the SN, the spore pellet was re-suspended in 0.5 mL 2TY medium and heat shock procedure was conducted for 5 min at 50 °C in a HLC thermo-block heater and then stored on ice.

*Streptomyces* spores and *E. coli* were combined in a volume ratio of 2:1; 0.5 mL spores and 0.25 mL of *E. coli* cells were suspended in an Eppendorf tube and 350 µL of the mixture was plated on SM agar plates. On the next day, each plate was overlaid with 25 µg/mL nalidixic acid and 50 µg/mL apramycin (final concentration in the plate) in 1 mL MQ-$H_2O$. The plates were incubated at 30 °C. As soon as exconjugants *of Streptomyces* were detected, these single colonies were transferred to a new SM-agar plate with 25 µg/mL nalidixic acid and 50 µg/mL apramycin. After these isolated exconjugants showed vital growth, these cultures were transferred again to a SM-agar plate containing this time only 50 µg/mL apramycin. This agar culture was incubated at 30°C until sporulation occurred.

**Tab. S2** List of oligonucleotides used in this study

| No. | Primer name | Primer sequence 5'–3' |
|---|---|---|
| 1 | JHuFw_*citT$_a$*_DK1622_KO | ATATAAGCTTGACATGTTCCCCCAGCACCT |
| 2 | JHuRv_*citT$_a$*_DK1622_KO | ATATACTAGTTTCATTGTCCAGCACCTGCC |
| 3 | JHuFw_*citB*_DK1622_KO | ATATAAGCTTGACCTGATTGAACGCGTCCT |
| 4 | JHuRv_*citB*_DK1622_KO | ATATACTAGTCCGAACGGGAAGTAGACGTA |
| 5 | JHuFw_*citC*_DK1622_KO | ATATAAGCTTAGTTCTTCCGCTCGGCATACC |
| 6 | JHuRv_*citC*_DK1622_KO | ATATACTAGTGGGTCGGTCTGCTCGTAGTG |
| 7 | JHuFw_*mx_pep*_KO | ATATAAGCTTCCTGGGACGGCAAGAAGGTG |
| 8 | JHuRv_ *mx_pep* _KO | ATATACTAGTCGGTGCTGACGGACGTCTTGTAG |
| 9 | JHuFw_*citA*_activate | ATATCATATGAAGAAGGCCCTGTACTCTTTG |
| 10 | JHuRv_*citA*_activate | ATATGAATTCCGTGCCCGGACATGATCATC |
| 11 | JHuFw_ citT$_a$_activate | ATATCATATGAGTCCATCCAACAGGCGTT |
| 12 | JHuRv_ citT$_a$_activate | ATGAATTCATCTGCATGATGAGGCCCG |
| 13 | JHuFw_PermE* | ATATTTAATTAAAAGCGAGCGAAGCCACTGAG |
| 14 | JHuRv_PermE* | ATATTCTAGAATATCTTAAGGGCCATATGTGGGGT CCTCC |
| 15 | JHuFw_cittilin_operon_NdeI | ATATCATATGAAGAAGGCCCTGTACTCTTTGG |
| 16 | JHuRv_cittilin_operon_BspTI | ATATCTTAAGTCGAACTGCTGGCGGAGTGA |
| 17 | JHuRv_cittilin_operon_Bsp1407I | ATATTGTACATCGAACTGCTGGCGGAGTG |
| 18 | JHuFw_DK1622_*mx_pep* | ATATTGTACACCCGGTGTTGTCTGGTTGAC |
| 19 | JHuRv_DK1622_*mx_pep* | ATATCTTAAGCATCAAGGGAACACCCGAGG |
| 20 | JHuFw_DK1622_*citC* | ATATCATATGGAAAACCTGTATTTTCAGGGCGGCA TGCGGCGGGAGCATGAAGG |
| 21 | JHuRv_DK1622_ *citC* | ATATGAATTCCGCCTGAGTCAGTCCTCTGTCG |
| 22 | JHuFw_DK1622_*mx_pep* | ATATCATATGTCCTACCCGGCGACC |
| 23 | JHuRv_DK1622_*mx_pep* | ATATAAGCTTTCAGCGGCCCTGCGCCGCCAC |
| 24 | JHuFw_DK1622_*citB* | ATATCCATGGGGTTGGGTCTCAAGAGCTGGTCGA |
| 25 | JHuRv_DK1622_*citB* | ATATAAGCTTCATGCTCCCGCCGCATGC |
| 26 | JHuFw_DK1622_*citB*_Strepto | CTTCATATGGAAAACCTGTATTTTCAGGGCGGCAT GGAGCGCATGGTTCGCT |
| 27 | JHuRv_DK1622_*citB*_Strepto | ATGAATTCCTTCATGCTCCCGCCGC |
| 28 | JHuFw_Cbvi35_*citB*_Strepto | CTTCATATGGAAAACCTGTATTTTCAGGGCGGCAT GGCGGGGGAGGACTCCT |
| 29 | JHuRv_Cbvi35_*citB*_Strepto | ATAAGCTTTCATGCGCTCACCTCTTG |
| 30 | JHuFw_MCy9171_*citB*_Strepto | CTTCATATGGAAAACCTGTATTTTCAGGGCGGCAT GTCGACGCGGGAGGAAT |
| 31 | JHuRv_MCy9171_*citB*_Strepto | ATAAGCTTTCAGGCTCGTGCTGCATG |
| 32 | JHuFw_pCJW93_exchange_noHis | CGTCAGACCCCGTAGAAAAG |

| | | |
|---|---|---|
| 33 | JHuRv_pCJW93_exchange_noHis | ATATCATATGTGTCCGCTCCCTTCTCTG |
| 34 | JDa14Rv_pCJW93_test1 | CGCTGCTGTGATGATGATG |
| 35 | JDa14Rv_pCJW93_test2 | GATGATGATGATGGCTGCTG |
| 36 | JHuFw_DK1622_*citB*_Strepto_no_stop_ C-His6 | CTTCATATGGAGCGCATGGTTCGCT |
| 37 | JHuRv_DK1622_*citB*_Strepto_no_stop_C-His6 | ATGAATTCTCAATGGTGATGGTGATGGTGGCCGGC GCCTGCTCCCGCCGCATG |
| 38 | JHuFw_Cbvi35_*citB*_Strepto_no_stop_ C-His6 | CTTCATATGGCGGGGGAGGACTCCT |
| 39 | JHuRv_Cbvi35_*citB*_Strepto_no_stop_C-His6 | ATGAATTCTCAATGGTGATGGTGATGGTGGCCGGC GCCTGCGCTCACCTCTTGAGGAAG |
| 40 | JHuFw_MCy9171_*citB*_Strepto_no_stop_C-His6 | CTTCATATGTCGACGCGGGAGGAAT |
| 41 | JHuRv_MCy9171_*citB*_Strepto_no_stop_C-His6 | ATGAATTCTCAATGGTGATGGTGATGGTGGCCGGC GCCGGCTCGTGCTGCATGCC |

**Tab. S3** List of oligonucleotides used as sequencing primers

| No. | Primer name | Primer sequence 5’–3’ |
|---|---|---|
| 1 | Test_Fw1_DK1622_citT$_a$_KO | CGACTACCGTGAGTCCATCCAACA |
| 2 | Test_Rv2_DK1622_citT$_a$_KO | TCCACGCGTTGACCGTCTGAA |
| 3 | Test_Rv3_DK1622_citT$_a$_KO | TCTACGTGTTCCGCTTCCTTTAGCAG |
| 4 | Test_Fw4_DK1622_citT$_a$_KO | CCTTTGAGTGAGCTGATACCGCTCG |
| 5 | Test_Fw1_DK1622_citB_KO | TGGGTCTCAAGAGCTGGTCGAC |
| 6 | Test_Rv2_DK1622_citB_KO | CAGGTATGCCGAGCGGAAGAAC |
| 7 | Test_Rv3_DK1622_citB_KO | ATTCAGGCTGCGCAACTGTTGG |
| 8 | Test_Fw4_DK1622_citB_KO | GCCACCTCTGACTTGAGCGTC |
| 9 | Test_Fw1_DK1622_*citC*_KO | CTCTCACACAACACAGGAAGGCG |
| 10 | Test_Rv2_DK1622_*citC*_KO | CTTCACCTGCTGTCACCCCG |
| 11 | Test_Rv3_DK1622_*citC*_KO | CGCCCAGTCTAGCTATCGCCA |
| 12 | Test_Fw4_DK1622_*citC*_KO | CTTCCGGCTCGTATGTTGTGTGG |
| 13 | Test_Fw1_DK1622_*mx_pep*_KO | CCACAAGGACAAGGAGAAGGCC |
| 14 | Test_Rv2_DK1622_*mx_pep*_KO | CGTTGCTGCCGCCGTAGAT |
| 15 | Test_Rv3_DK1622_*mx_pep*_KO | TCTTCGCTATTACGCCAGCTGG |
| 16 | Test_Fw4_DK1622_*mx_pep*_KO | CGTATTACCGCCTTTGAGTGAGCTG |
| 17 | Test_Fw1_DK1622_*citA*_activate | CTGTGCCACATTTCACCGCG |
| 18 | Test_Rv2_DK1622_*citA*_activate | CTCACCGCCAGCATCGTTCC |
| 19 | Test_Rv3_DK1622_*citA*_activate | CTGCGTTATCCCCTGATTCTGTGG |
| 20 | Test_Fw4_DK1622_*citA*_activate | TGTCAAGCTGCTGTTTTCGCCG |
| 21 | Test_Fw1_DK1622_*citT$_a$*_activate | AGAAGAAACACCCAGGCTTTGAC |
| 22 | Test_Rv2_DK1622_*citT$_a$*_activate | CACCATCATCTTCCCGAGCA |

| | | |
|---|---|---|
| 23 | Test_Rv3_DK1622_ citT$_a$ _activate | TTAGCTCACTCATTAGGCACCC |
| 24 | Test_Fw4_DK1622_ citT$_a$ _activate | GGATCCAATAGGTCGCCGAA |
| 25 | Test_Fw1_Seq_pCJW93_*citB* | GACAAAACTTTAGATCTGGG |
| 26 | Test_Rv2_Seq_pCJW93_*citB* | GAACGTCCGGGCTTGCAC |

**Tab. S4 List of plasmids used in this study**

| No. | Plasmid name/ characteristic | Size [kb] | Function | Reference |
|---|---|---|---|---|
| 1 | pCR2.1-TOPO (EcoRI, religated) | 3.931 | Used as template to PCR-amplify *tn5_kanR* gene | TOPO®TA Cloning® Kit Thermo Fisher Scientific™ |
| 2 | pFP$_{Van}$_*pcyA* | 6.181 | pCR2.1-TOPO derivative used as vector to ligate PCR products for subsequent vanillate-induced gene expression in *M. xanthus* DK1622 | *(17)* |
| 3 | pSET152 | 5.549 | Shuttle vector for expression of secondary metabolites in *Streptomyces*, harboring *apraR* | *(18)* |
| 4 | pAB03 | 5.429 | Used as template to PCR-amplify P$_{ERME*}$ | *(19)* |
| 5 | pET-28b | 5.368 | Used as expression vector for recombinant protein production | (Novagen) |
| 6 | pHisTEV | 5.365 | Used as expression vector for recombinant protein production | *(20,21)* |
| 7 | pCJW93 | 8.672 | Used as expression vector for recombinant protein production in *Streptomyces coelicolor* CH999 | *(22)*, Kindly provided by Peter F. Leadlay |

**Tab. S5 List of PCR-amplified constructs**

| No. | PCR product name/ characteristic | Size [bp] | Template | Primer used |
|---|---|---|---|---|
| 1 | *citT<sub>a</sub>*_homology | 593 | gDNA from *M. xanthus* DK1622 | primer No.1 primer No. 2 |
| 2 | *citB*_homology | 994 | gDNA from *M. xanthus* DK1622 | primer No. 3 primer No. 4 |
| 3 | *citC*_homology | 531 | gDNA from *M. xanthus* DK1622 | primer No. 5 primer No. 6 |
| 4 | *mx_pep*_homology | 1021 | gDNA from *M. xanthus* DK1622 | primer No. 7 primer No. 8 |
| 5 | *citA*_activate | 970 | gDNA from *M. xanthus* DK1622 | primer No. 9 primer No. 10 |
| 6 | *citT<sub>a</sub>*_activate | 1226 | gDNA from *M. xanthus* DK1622 | primer No. 11 primer No. 12 |
| 7 | $P_{ermE*}$ | 470 | pAB03erm* | primer No.13 primer No.14 |
| 8 | DK1622_ *citA–citC*_operon_NdeI_BspTI | 2233 | gDNA from *M. xanthus* DK1622 | primer No.15 primer No.16 |
| 9 | DK1622_ *citA–citC*_operon_NdeI_Bsp1407I | 2233 | gDNA from *M. xanthus* DK1622 | primer No.15 primer No.17 |
| 10 | DK1622_*mx_pep*_Bsp1407I_ BspTI | 2407 | gDNA from *M. xanthus* DK1622 | primer No.18 primer No.19 |
| 11 | Cittilin_DK1622_CitC_recombinant_protein | 667 | gDNA from *M. xanthus* DK1622 | primer No. 20 primer No. 21 |
| 12 | Cittilin_DK1622_MX_PEP_recombinant_protein | 2087 | gDNA from *M. xanthus* DK1622 | primer No. 22 primer No. 23 |
| 13 | Cittilin_DK1622_CitB_recombinant_protein | 1461 | gDNA from *M. xanthus* DK1622 | primer No. 24 primer No. 25 |
| 14 | Cittilin_DK1622_CitB_recombinant_protein_Strepto | 1456 | gDNA from *M. xanthus* DK1622 | primer No. 26 primer No. 27 |
| 15 | Cittilin_Cbvi35_CitB_recombinant_protein_Strepto | 1475 | gDNA from *Cystobacter* Cb vi35 | primer No. 28 primer No. 29 |
| 16 | Cittilin_Mcy9171_CitB_recombinant_protein_Strepto | 1475 | gDNA from MCy9171 | primer No. 30 primer No. 31 |
| 17 | pCJW93_noHistag_exchange_construct | 1388 | pCJW93 | primer No. 32 primer No. 33 |
| 18 | Cittilin_DK1622_CitB_recombinant_protein_Strepto_no_stop_C-His | 1472 | gDNA from *M. xanthus* DK1622 | primer No. 36 primer No. 37 |

| | | | | |
|---|---|---|---|---|
| 19 | Cittilin_Cbvi35_CitB_recombinant_protein_Strepto_no_stop_C-His | 1393 | gDNA from *Cystobacter* Cb vi35 | primer No. 38 primer No. 39 |
| 20 | Cittilin_Mcy9171_CitB_recombinant_protein_Strepto_no_stop_C-His | 1471 | gDNA from MCy9171 | primer No. 40 primer No. 41 |

**Tab. S6 List of genetic constructs generated in this study**

| No. | Plasmid name | Construction details/ characteristic |
|---|---|---|
| 1 | pCR2.1-TOPO_*citT$_a$*_KO | Construct obtained by conventional restriction ligation of plasmid pCR2.1-TOPO and PCR product No. 1 |
| 2 | pCR2.1-TOPO_*citB*_KO | Construct obtained by conventional restriction ligation of plasmid pCR2.1-TOPO and PCR product No. 2 |
| 3 | pCR2.1-TOPO_*citC*_KO | Construct obtained by conventional restriction ligation of plasmid pCR2.1-TOPO and PCR product No. 3 |
| 4 | pCR2.1-TOPO_*mx_pep*_KO | Construct obtained by conventional restriction ligation of plasmid pCR2.1-TOPO and PCR product No. 4 |
| 5 | $pFP_{Van}$_*citA*_activate | Construct obtained by conventional restriction ligation of plasmid pFPVan_pcyA and PCR product No. 5 |
| 6 | $pFP_{Van}$_*citT$_a$*_activate | Construct obtained by conventional restriction ligation of plasmid pFPVan_pcyA and PCR product No. 6 |
| 7 | pSET152_$P_{ermE*}$ | Construct obtained by conventional restriction ligation of plasmid pSET152 and PCR product No. 7 |
| 8 | pSET152_$P_{ermE*}$_*citA–citC*_operon | Construct obtained by conventional restriction ligation of construct No. 7 and PCR product No. 8 |
| 9 | pSET152_$P_{ermE*}$_*citA–citC*_operon_*mx_pep* | Construct obtained by conventional restriction ligation of construct No. 7 and PCR products No. 8 and No. 9 |

| | | |
|---|---|---|
| 10 | pET28b_DK1622_CitC | Construct obtained by conventional restriction ligation of plasmid pET28b and PCR product No. 11 |
| 11 | pET28b_DK1622_MX_PEP | Construct obtained by conventional restriction ligation of plasmid pET28b and PCR product No. 12 |
| 12 | pHisTEV_DK1622_CitB | Construct obtained by conventional restriction ligation of plasmid pHisTEV and PCR product No. 13 |
| 13 | pCJW93_DK1622_CitB | Construct obtained by conventional restriction ligation of plasmid pCJW93 and PCR product No. 14 |
| 14 | pCJW93_Cbvi35_CitB | Construct obtained by conventional restriction ligation of plasmid pCJW93 and PCR product No. 15 |
| 15 | pCJW93_MCy9171_CitB | Construct obtained by conventional restriction ligation of plasmid pCJW93 and PCR product No. 16 |
| 16 | pCJW93noHis | Construct obtained by conventional restriction ligation of plasmid pCJW93 and PCR product No. 17 |
| 17 | pCJW93noHis_DK1622_CitB | Construct obtained by conventional restriction ligation of genetic construct No. 16 and PCR product No. 18 |
| 18 | pCJW93noHis_Cbvi35_CitB | Construct obtained by conventional restriction ligation of genetic construct No. 16 and PCR product No. 19 |
| 19 | pCJW93noHis_MCy9171_CitB | Construct obtained by conventional restriction ligation of genetic construct No. 16 and PCR product No. 20 |

#### Bacterial cultures and preparation of cryogenic long-term stocks

Myxobacteria were preserved at -80 °C. Ten mL of liquid culture was transferred to a 2 mL Eppendorf tube and centrifuged for 2 min, 7000 rpm. 1.5 mL of SN was discarded, leaving 0.5 mL SN in the Eppendorf tube. The CP was re-suspended by pipetting up and down and then mixed with 0.5 mL glycerol (50%). The mixture was transferred into a cryogenic vial.

For preservation of spores of *Streptomyces* spp. at -80 °C, 2 mL of glycerol (20%) was added on a sporulating agar culture. The glycerol was spread with a cell spreader on the plate to collect the spores. The glycerol-spore mixture was afterwards transferred from plate via pipetting into a cryogenic vial.

Bacterial strains for the experiments of this work, their relevant characteristics and sources are listed in the **Tab. S7**

**Tab. S7** Bacterial strains used in this study.

| **Strain** | **Function** | **Origin** |
|---|---|---|
| *E. coli* HS996 | Standard cloning host | Invitrogen |
| *E. coli* ET12567 | Donor for *Streptomyces* conjugation | ATCC® BAA-525™ |
| *E. coli* Lemo21 | Recombinant protein production | Novagen |
| *E. coli* BL21 | Recombinant protein production | Novagen |
| *Myxococcus xanthus* DK1622 | Investigated strain in this study | Internal strain collection |
| *Streptomyces albus* del14 | Host for heterologous production of cittilin A | Kindly provided by Andriy Luzhetskyy *(23)* |
| *Streptomyces coelicolor* CH999 | Host for recombinant protein production | Kindly provided by Peter F. Leadlay *(24)* |
| *E. coli* (TolC-deficient mutant) | Gram-negative bacterium to test bacterial cell entry | Internal strain collection *(25)* |

#### Cultivation media and buffers

The pH of all cultivation media was adjusted before autoclaving. Autoclaving was conducted at 121°C for 20 min.

2TY-medium

| Ingredient | Concentration |
|---|---|
| Tryptone | 16 g/L |
| Yeast extract | 20 g/L |
| NaCl | 5 g/L |

pH 6.8 +/- 0.2 with $NaOH_{aq}$

COM-medium

| Ingredient | Concentration |
|---|---|
| Glucose | 25 g/L |
| Soybean flour | 25 g/L |
| Baker's yeast (fresh) | 3 g/L |
| NaCl | 2 g/L |
| $(NH_4)_2SO_4$ | 2 g/L |
| $CaCO_3$ | 2 g/L |
| $K_2HPO_4$ | 0.15 g/L |

pH 8.4 +/- 0.2 with $NaOH_{aq}$

CTT-medium

| Ingredient | Concentration |
|---|---|
| Casitone | 10 g/L |
| Tris | 10 mM |
| $KH_2PO_4$ | 1 mM |
| $MgSO_4$ | 8 mM |

pH 7.6 with $KOH_{aq}$

LB-medium

| Ingredient | Concentration |
|---|---|
| Yeast extract | 5 g/L |
| Tryptone | 10 g/L |
| NaCl | 5 g/L |

pH 7.2 with $KOH_{aq}$

SM-agar

| Ingredient | Concentration |
|---|---|
| D-mannitol | 20 g/L |
| Soybean flour | 20 g/L |
| $MgCl_2$*$6H_2O$ | 10 mM |
| Agar | 20 g/L |

Super YEME-medium

| Ingredient | Concentration |
|---|---|
| Yeast extract | 3 g/L |
| Peptone | 5 g/L |
| Glucose | 10 g/L |
| Malt extract | 3 g/L |
| Sucrose | 340 g/L |
| Glycine | 5 g/L |
| $MgCl_2$ x $6H_2O$ | 2.35 g/L |
| L-proline | 75 mg/L |
| L-arginine | 75 mg/L |
| L-cysteine | 75 mg/L |
| L-histidine | 100 mg/L |
| Uracil | 15 mg/L |

TSB-medium

| Ingredient | Concentration |
|---|---|
| Tryptic soy broth | 30 g/L |

Lysis buffer

| Ingredient | Concentration |
|---|---|
| NaCl | 500 mM |
| BIS-TRIS | 20 mM |
| Imidazole | 20 mM |
| Glycerol | 10% (m/m) |

pH 6.8 with $NaOH_{aq}$

Elution buffer

| Ingredient | Concentration |
|---|---|
| NaCl | 500 mM |
| BIS-TRIS | 20 mM |
| Imidazole | 250 mM |
| Glycerol | 10% (m/m) |

pH 6.8 with $NaOH_{aq}$

Protein buffer

| Ingredient | Concentration |
|---|---|
| NaCl | 500 mM |
| BIS-TRIS | 20 mM |
| Glycerol | 5% (m/m) |

pH 6.8 with $NaOH_{aq}$

Lysis buffer No. 3

| Ingredient | Concentration |
|---|---|
| NaCl | 500 mM |
| BIS-TRIS | 20 mM |
| Imidazole | 20 mM |

pH 6.8 with $NaOH_{aq}$

Elution buffer No. 3

| Ingredient | Concentration |
|---|---|
| NaCl | 500 mM |
| BIS-TRIS | 20 mM |
| Imidazole | 250 mM |

pH 6.8 with $NaOH_{aq}$

Protein buffer No. 3

| Ingredient | Concentration |
|---|---|
| NaCl | 500 mM |
| BIS-TRIS | 20 mM |

pH 6.8 with $NaOH_{aq}$

#### Crude extracts preparation for analysis of secondary metabolism

*Myxococcus xanthus* DK 1622 and mutant strains were grown on CTT agar plates for 3–5 days at 30 °C. For the preparation of liquid pre-cultures, a suitable portion of overgrown agar was transferred to inoculate 20 mL of CTT medium in an Erlenmeyer flask (300 mL) and incubated at 30 °C and 200 rpm for 2–4 days. 1 mL of the pre-culture was used to inoculate 50 mL of CTT medium containing appropriate antibiotic selection and 2% amberlite resin XAD-16 (Sigma Aldrich).

Spores of *Streptomyces albus* del14 and mutant strains were used to inoculate 20 mL of TSB medium in an Erlenmeyer flask (300 mL) and incubated at 30 °C and 200 rpm for 2–4 days. 1–2 mL of the pre-culture was used to inoculate 50 mL of COM medium containing appropriate antibiotic selection and 2% amberlite resin XAD-16 (Sigma Aldrich).

In order to obtain statistically significant results, three independent transformants of each recombinant strain were selected and fermentations were performed at least in triplicates. The resulting fermentation broth were harvested by centrifugation at 8000 rpm for 10 min (Eppendorf centrifuge 5810 R). The SN was discarded, whereas the residual material was extracted first with 25 mL MeOH, stirred for 1 h, filtered through filter paper (folded filters grade: 3hw from Sartorius) into a round bottom flask and afterwards this procedure was repeated with 25 mL acetone. The solvent of the filtered extracts was removed under vacuum (BÜCHI Rotavapor R–210) and the extracts re-dissolved in 1.5 mL MeOH and stored at -20 °C. The re-dissolved extract were diluted with MeOH (1:3 (extract/MeOH (v:v)) centrifuged at 13000 g for 10 min (VWR centrifuge ECN521-3601, Hitachi Koki Co., Ltd) and 1 µL of the SN was subjected to HPLC-MS analysis as described further below.

#### Analysis of secondary metabolites in broth extracts, and enzymatic reactions

The secondary metabolism of broth extracts was analyzed by HPLC-HRESI-DAD-MS on a Bruker maXis 4G mass spectrometer coupled with a Dionex Ultimate 3000 RSLC system using a BEH C18 column (100 × 2.1 mm, 1.7 µm, Waters, Germany) with a gradient of 5–95% acetonitrile (ACN) + 0.1% formic acid (FA) in $H_2O$ + 0.1% FA at 0.6 mL/ min and 45 °C over 9 or 18 min with UV detection by a diode array detector at 200–600 nm. Mass spectra were acquired from 150 to 2000 m/z at 2 Hz. The detection was performed in the positive MS mode. The plugin for Chromeleon Xpress (Dionex) was used for operation of UltiMate 3000 LC System. HyStar (Bruker Daltonic) was used to operate on maXis 4G speed MS system. HPLC-MS mass spectra were analyzed with DataAnalysis 4.2 (Bruker Daltonic).

##### Analysis of recombinant proteins via LC-MS

Recombinant proteins were analyzed on a Dionex Ultimate 3000 UPLC system (Thermo Scientific™) coupled with an maXis4G Q-TOF MS (Bruker) using an ESI in positive mode. The samples were run on a Aeris Widepore XB-C8, 150 x 2.1 mm, 3.6 µm dp column (Phenomenex, USA). LC conditions: A: $H_2O$ dd + 0.1% FA; B: Acetonitrile + 0.1% FA at a flow rate of 300 µL/min and 45 °C. 0 min: 98% A / 2% B, 0.5 min: 98% A / 2% B, 10.5 min: 25% A / 75 % B, 13.5 min: 25% A / 75% B, 14 min: 98% A / 2% B. The LC flow was split to 75 µL/min before entering the maXis4G hr-ToF mass spectrometer (Bruker Daltonics, Bremen, Germany) using the standard Bruker ESI source. In the source region, the temperature was set to 180 °C, the capillary voltage was 4000 V, the dry-gas flow was 6.0 L/min and the nebulizer was set to 1.1 bar. Mass spectra were acquired in positive ionization mode ranging from 150–2500 m/z at 2.5 Hz scan rate. Protein masses were deconvoluted by using the Maximum Entropy algorithm (Copyright 1991-2004 Spectrum Square Associates, Inc.).

#### Compound isolation

##### Analysis during purification

All measurements to analyze the mass of cittilin A and B during purification were performed on a Dionex Ultimate 3000 RSLC system coupled to the amaZon iontrap MS using a BEH C18, 100 x 2.1mm, 1.7 µm dp column equipped with a C18 precolumn (Waters). Samples of 1 µL were separated by a gradient from (**A**) $H_2O$ + 0.1% formic acid (FA) to (**B**) ACN + 0.1% FA at a flow rate of 0.6 mL/min and 45 °C. The gradient was initiated by a 0.5 min isocratic step at 5% **B**, followed by linear increase to 95% **B** in 18 min. After a 2 min step at 95% **B** the system was re-equilibrated to the initial conditions (5% **B**). UV-spectra were recorded by a DAD in the range from 200–600 nm. The detection was performed in the positive ESI MS/MS mode.

##### Size exclusion chromatography via gel filtration

The CP of a 10 L fermentation culture with 2% XAD-16 was used to prepare a crude extract by mixing the CP with 400 mL (MeOH), stirring for 1 h and filtering through glass wool into a round bottom flask. This extraction was repeated once; afterwards the extraction fraction was dried using the rotary evaporator (BÜCHI Rotavapor R–210). The dried residue was re-dissolved MeOH, in order to run the obtained extract on a methanolic Sephadex® LH20 column. The cittilin-containing fraction was placed on a silica gel column, which was run with a $CHCl_3$ : MeOH gradient (starting 9:1 (v/v), ending with pure MeOH). For further purification another Sephadex® LH20 column was performed with $H_2O$/ MeOH (3:2).

#### Semi-preparative HPLC chromatography

Semi-preparative HPLC purification was done using a Dionex Ultimate 3000 SDLC low pressure gradient system on a XBridge® peptide BEH C18 OBD™ prep column (138 Å, 5 µm, 10 mm × 250 mm, 1/pkg). Column temperature was stabilized at 45 °C with the eluents $H_2O$ + 0.1% FA as A and ACN + 0.1% FA as B, at a flow rate of 1.5 mL/min. Detection of cittilin A was facilitated via mass spectrometry on the Agilent 1100 series coupled to the HCT 3D ion trap or with a UV detector on the Dionex 3000 SL systems by UV absorption at 256 nm and 320 nm. The gradient starts with a plateau at 95% A for 2 min followed by a ramp to 73% A during 5.3 min. Then, A content was kept to 73% during 11 min and finally ramped to 5% A during 2 min. A content is kept at 5% for 2 min and then ramped back to 95% during 30 s. The column was re-equilibrated at 95% A for 3 min. The white compound was subsequently dried by lyophilization yielding cittilin A and B.

### 2 Results

### NMR spectroscopic data

**Tab. S8** $^{1}H$ and $^{13}C$ NMR spectral data of cittilin A and B; recorded at 500 and 125 MHz in $CD_3OD$; d: doublet; dd: doublet of doublets; s: singlet; m: multiplet.

| Amino acid | Position | Cittilin A $^{1}H$ (*J*) ppm (Hz) | $^{13}C$ ppm | Cittilin B $^{1}H$ (*J*) ppm (Hz) | $^{13}C$ ppm | Multiplicity |
|---|---|---|---|---|---|---|
| **L-tyrosine 1** | 1-CO | --- | 169.9 | --- | 170.1 | --- |
| | 2-CH | 4.21 | 53.8 | 4.08 | 54.1 | M |
| | 3a-CH**H** | 3.46 | 36.7 | 3.41 | 37.6 | M |
| | 3b-CH**H** | 3.05 (6/15) | 36.7 | 2.99 (6/15) | 37.6 | dd |
| | 4-C | --- | 129.7 | --- | 128.0 | --- |
| | 5-CH | 6.92 | 137.0 | 6.90 | 137.0 | m |
| | 6-C | --- | 125.9 | --- | 125.6 | --- |
| | 7-C | --- | 157.9 | --- | 155.9 | --- |
| | 8-CH | 7.21 (2/8) | 131.7 | 7.19 (2/8) | 131.7 | dd |
| | 9-CH | 7.01 (9) | 112.0 | 6.98 (8) | 111.8 | d |
| | O-$CH_3$ | 3.88 | 56.3 | ---- | --- | s |
| **L-isoleucine** | 1-CO | --- | 169.9 | --- | 171.7 | --- |
| | 2-CH | 4.20 | 60.1 | 4.16 | 60.0 | m |
| | 3-CH | 1.77 | 37.1 | 1.77 | 37.1 | m |
| | 4a-CH**H** | 1.61 | 26.3 | 1.62 | 26.4 | m |
| | 4b-CH**H** | 1.27 | 26.3 | 1.26 | 26.4 | m |
| | 5-$CH_3$ | 0.95 (7) | 15.7 | 0.96 (6) | 15.9 | m |
| | 6-$CH_3$ | 0.92 (7) | 11.0 | 0.92 (7) | 10.9 | t |
| **L-tyrosine 2** | 1-CO | --- | 173.0 | --- | 171.5 | --- |
| | 2-CH | 3.68 | 59.8 | 3.68 | 59.8 | m |
| | 3a-CH**H** | 3.10 (2/14) | 40.7 | 3.16 (2/13) | 40.7 | dd |
| | 3b-CH**H** | 2.47 (4/13) | 41.0 | 2.47 (4/14) | 40.9 | dd |
| | 4-C | --- | 136.6 | --- | 135.4 | --- |
| | 5-CH | 5.78 (2) | 120.7 | 5.80 (2) | 120.8 | d |
| | 6-C | --- | 154.2 | --- | 152.6 | --- |
| | 7-C | --- | 145.2 | --- | 143.3 | --- |
| | 8-CH | --- | 129.7 | --- | 129.6 | |
| | 9-CH | 6.62 (2) | 126.3 | 6.63 (2) | 126.4 | d |
| **L-tyrosine 3** | 1-CO | --- | 176.8 | --- | 175.2 | --- |
| | 2-CH | 4.30 | 54.2 | 4.26 | 54.7 | m |
| | 3a-CH**H** | 3.61 | 37.3 | 3.61 | 37.4 | m |

| | | | | | | |
|---|---|---|---|---|---|---|
| | 3b-CH**H** | 2.98 (7/14) | 37.3 | 2.94 (m) | 37.4 | dd |
| | 4-C | --- | 135.5 | --- | 133.9 | --- |
| | 5-CH | 7.22 | 135.4 | 7.19 | 135.4 | m |
| | 6-C | 6.83 (3/8) | 124.8 | 6.79 (3/9) | 124.6 | dd |
| | 7-C | --- | 164.5 | --- | 162.5 | --- |
| | 8-CH | 7.47 (2/8) | 127.0 | 7.45 /3/9) | 127.0 | dd |
| | 9-CH | 7.35 (2/8) | 130.7 | 7.36 (2/8) | 131.0 | dd |

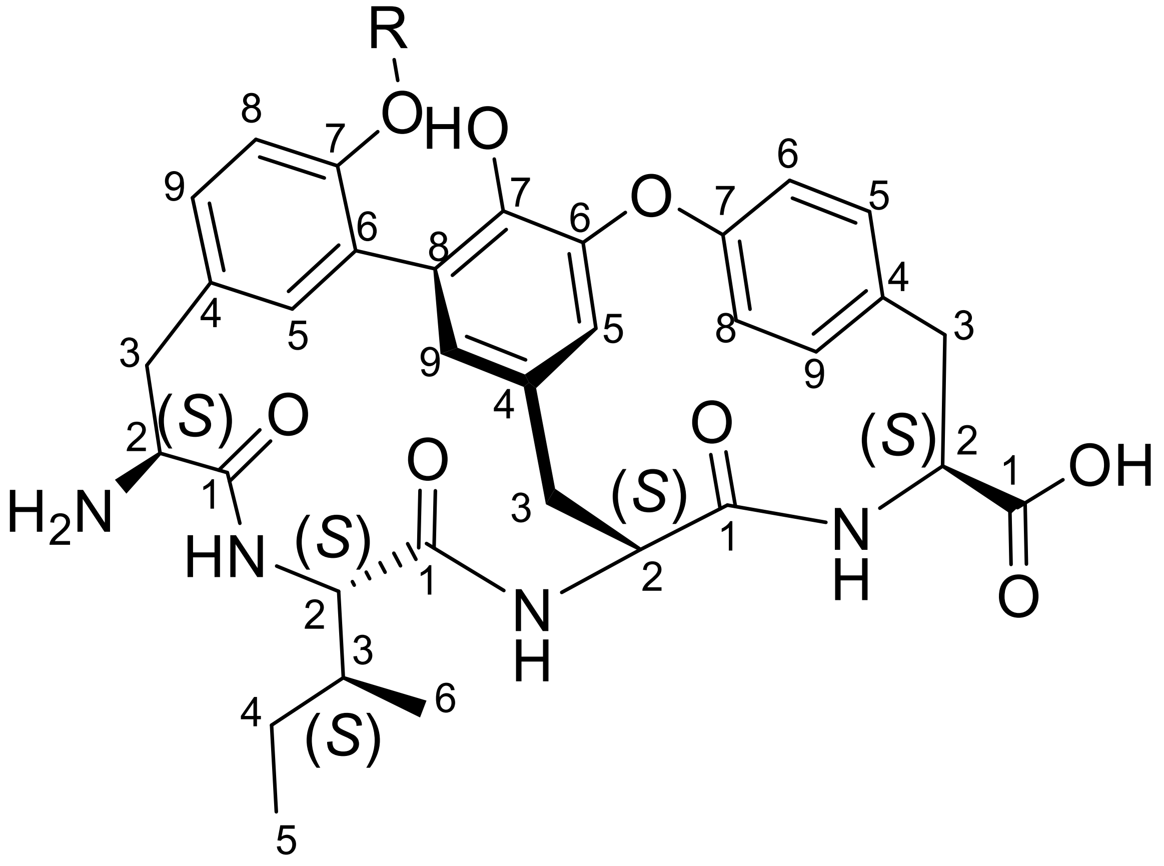

cittilin A (R=$CH_3$)

cittilin B/RP-66453 (R=H)

**Fig. S1** Structure and carbon numbering of cittilin A and B.

### Bioinformatic investigation of different cittilin A and B producing myxobacterial strains

The biosynthetic pathway of cittilin and their genetic organization was investigated by *in silico* analysis. As previously described by Krug et al. *(26)* several producers of cittilins are known, in particular the well described *M. xanthus* DK1622 (MCy9151), but also the closely related strains DK897 (MCy8986) and ST200611 (MCy11474). All those strains are LC-MS-confirmed producers with available whole genome sequences. In order to find more producer strains, the translated nucleotide sequence of the biosynthetically involved cytochrome P450 (CitB) from the characterized producer MCy9151 was used as query input to find via discontiguous MEGABLAST in the private MINS genome database and the public available database further candidate BGCs for cittilin A.

The list of similar citB homologs in different strains showed several potential results in the private MINS genome database and three hits in the public database, however only hits with identical sites up to 78% and with query coverage higher than 90% were considered as relevant hits. The following strains from the private MINS genome database were further investigated: MCy8278, MCy8337, MCy8375, MCy10608, MCy9171. From the publically available genome sequences eight hits were found. Six of these hits are strains, which are phylogenetically closely related to *M. xanthus* DK1622 (*Myxococcus xanthus* strain KF3.2.8c11, *Myxococcus xanthus* strain GH5.1.9c20, *Myxococcus xanthus* strain MC3.5.9c15, *Myxococcus xanthus* strain MC3.3.5c16, *Myxococcus xanthus* strain GH3.5.6c2, *Myxococcus xanthus* strain KF4.3.9c1) *(27)*, whereas *Myxococcus fulvus* HW-1 (MCy11108) and *Myxococcus hansupus mixupus* are phylogenetically less related. Roughly around 100 bp upstream of *citB*, a sequence encoding the core peptide YIYY followed by a stop codon could be found associated with all those hits, except for the strain *Myxococcus hansupus mixupus (28)*. Remarkably, it has a nucleotide sequence encoding the tetrapeptide YHYY. The precursor peptide (CitA), the cytochrome P450 (CitB) and the methyltransferase (CitC) amino acid sequence are highlighting high similarity to the characterized sequence from the strain MCy9151 that implies the BGC might be actively expressing a natural cittilin derivative. Since no metabolomic data or the strain itself is available, it was unfortunately not possible to search for this putative derivative. Interestingly the hit showing the highest percentage of identical sites to the query sequence derived from the strain A47 with 97% by showing at the same time very low query coverage of 12.96%. This strain has already been described as a non-producer of cittilins; however the genome harbors the same precursor peptide architecture like the confirmed cittilin producer strain MCy9151 *(29)*. The cytochrome P450 gene is truncated which is plausibly responsible for the lack of any cittilin in the metabolome of A47 and responsible for the low query coverage in the BLAST result. For this reason for further investigations only strains characterized by genomic data and being confirmed producers of cittilins by previously performed LC-MS measurements (**Fig. S2**), were considered for further *in silico* investigations. The following strains were used for in-depth *in silico* characterization: 1) MCy8278, 2) MCy8337, 3) MCy8375, 4) MCy10608, 5) MCy8986, 6) MCy9171, 8) MCy11108, 9) MCy11474. One benefit of having several BGCs is to use the information to determine the cluster borders. The final confirmation of the assumed borders is heterologous expression of the specific BGC in a heterologous host (assuming that this host is not capable of complementing any involved biosynthetic enzyme).

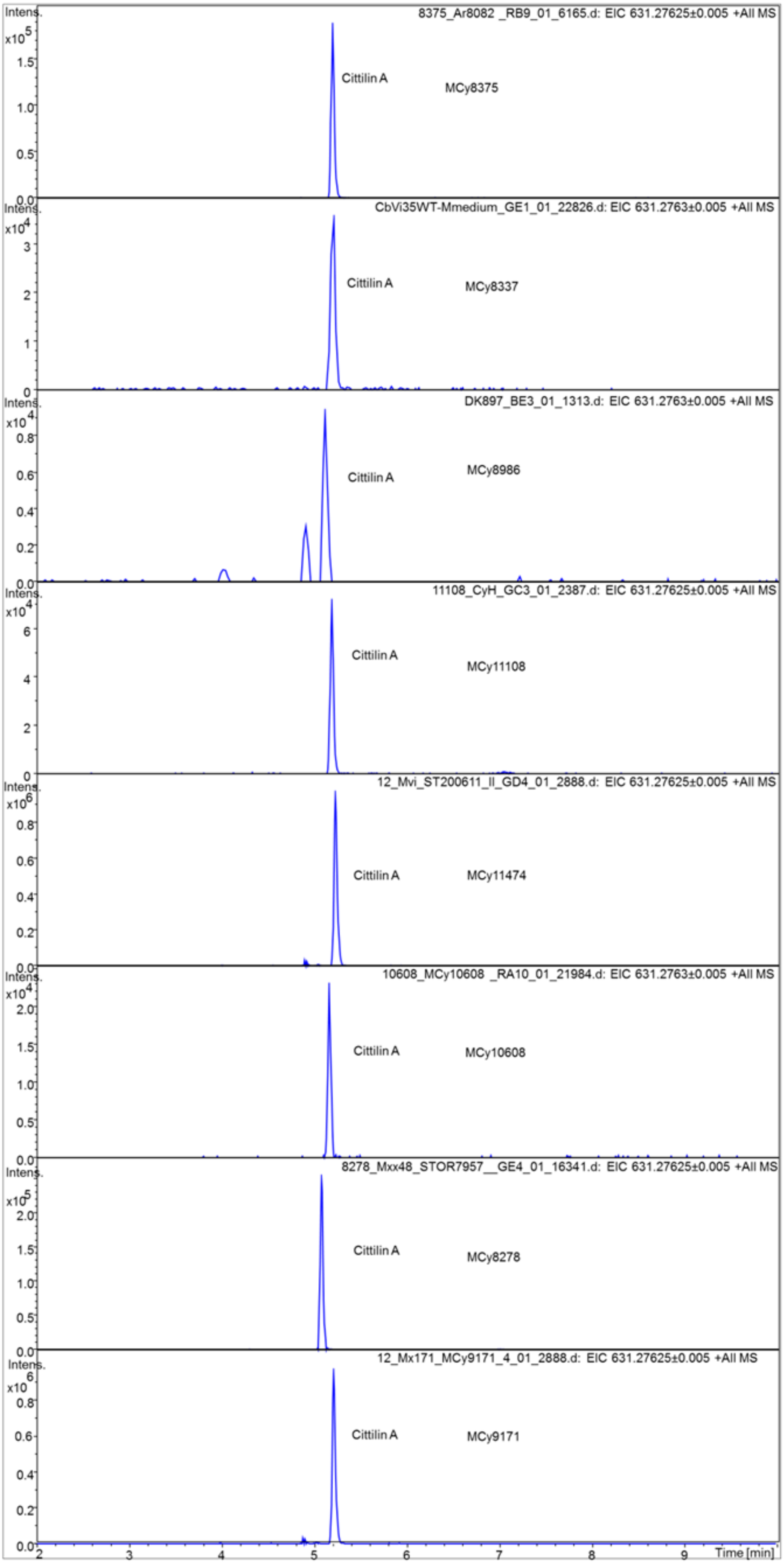

**Fig. S2** HPLC-MS extracted ion chromatograms of cittilin producers. The retention time and mass of cittilin A of 631.2768 *m/z* with a width of 7.9 ppm *m/z* in different strains with available genome data is shown. Cittilin B was detectable as well in lower concentration (not displayed).

To validate *a priori* that the precursor peptide gene (*citA*) consists of the 23 amino acid (aa) leader peptide gene and the four aa core peptide gene, the precursor peptide sequences of those nine different confirmed cittilin producer strains were aligned. The alignment is highlighting the identity among all producers within the leader and core peptide genes. Interestingly the putative ribosome binding site (RBS) sequence A(G)/GGAG is conserved in all cittilin producers. Directly upstream of the RBS and downstream of the core peptide the identity is significantly decreasing. Another hint for the start of peptide translation is the increasing 3rd-position GC content of all analyzed sequences *(30)*. Therefore, it can be assumed that for heterologous expression experiments the precursor peptide and the core peptide genes are sufficiently well defined for this work. Interestingly, the comparison of different leader peptide sequences showed two different residues upstream the core peptide. Unlike the other leader peptide sequences, the sequence in the strain MCy9171 is encoding the amino acids TT instead of AP. This position is likely critical for recognition of the putatively involved prolyl endopeptidase (MX PEP) to process the precursor peptide CitA. Therefore, for future *in vitro* studies with recombinant enzymes involved in the formation of cittilins, one has to consider the possibility that substrate specificity of the peptidase might deviate among different strains.

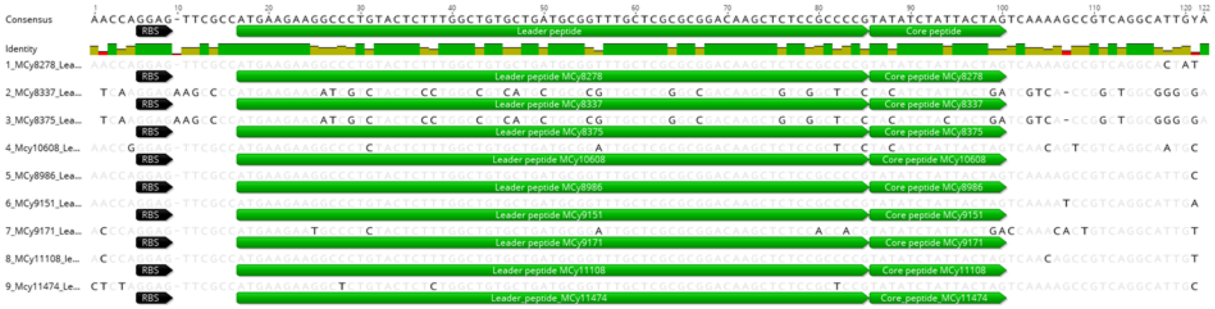

**Fig. S3** Nucleotide alignment of precursor peptide sequence of nine confirmed cittilin producers. The nucleotide sequence is highly conserved in particular the putative ribosome binding site (RBS) sequence A(G)/GGAG is conserved in all cittilin producers. Directly upstream of the RBS and downstream of the core peptide the similarity is significantly decreasing.

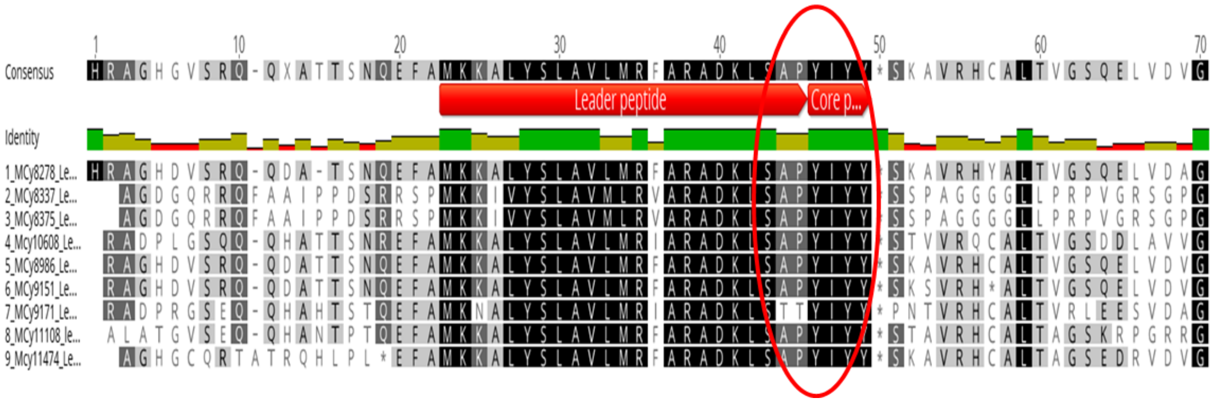

**Fig. S4** Protein alignment of precursor peptide sequences of nine confirmed cittilin producers. The amino acid sequence is highly conserved in particular the core peptide residues in the middle. However, the two different residues downstream the core peptide can vary. Unlike the other leader peptide sequences, the sequence in the strain MCy9171 is encoding the amino acids TT instead of AP (region indicated by red ring). Directly upstream of the core peptide the amino acid identity is significantly decreasing.

To narrow down essential genes for the production of cittilin, nine cittilin gene loci from confirmed producers were analyzed in detail. The strains MCy8278, MCy9151, MCy8986, MCy11474 and the non-producing strain A47 harboring the truncated cytochrome P450 gene *citB*, possess downstream of the precursor peptide gene *citA* four genes encoding an ABC transporter ($citT_a$–$citT_d$). The ABC transporter sequences are sharing high similarity in those strains and might play a plausible role for export of produced cittilin. Interestingly there exists another type of genetic organization, which has no ABC transporter upstream of the precursor peptide gene but an operon with an adenylate cyclase, a hypothetical protein and a peptidase. The BGCs which do not have an ABC transporter upstream of the precursor gene might have the gene somewhere else located in the genome, like it has already been demonstrated for the prolyl endopeptidase *mx pep*. MCy8837 and MCy8375 are sharing this gene cluster organization. The strain MCy10608 features upstream of the precursor peptide a tetracycline regulator gene and downstream of the methyltransferase gene *citC* a peptidoglycan-binding *lysM* gene. The surrounding genomic area in the strain of MCy10608 is unique in comparison to other cittilin producers. The producer strain MCy9171 and non-producing strain MCy8286 are not only very similar in the remarkable variation of their leader peptide but also the surrounding area is nearly identical.

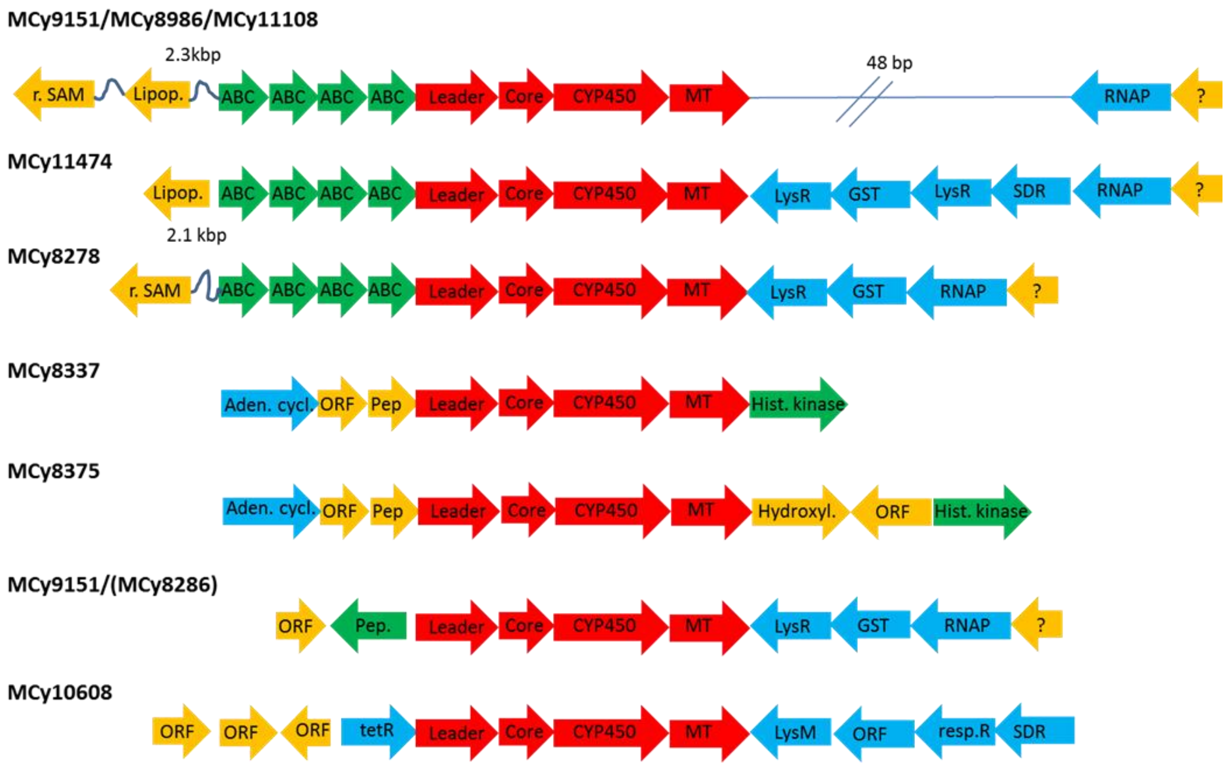

**Fig. S5 Schematic overview of cittilin BGCs in different strains.** Fundamentally the structural genes consisting of the precursor peptide gene (displayed separately), the cytochrome P450 and the methyltransferase, are present in the BGC. It seems to be that the ABC transporter upstream of the precursor peptide gene is conserved for the strains MCy9151, MCy8986, MCy11108, MCy11474 and MCy8278. The strains MCy9151, MCy8986 and MCy11108 are sharing a highly similar architecture not only the structural genes but also the surrounding genes 10 kb up- and downstream of the structural genes are almost identical according to the nucleotide alignments. Red: structural crucial genes, Green: Putative resistance conferring genes, Blue: Putative regulatory genes, Orange: Unknown function. ABC: ABC transporter, MT: Methyltransferase, rSAM: radical *S*-adenosyl-L-methionine (SAM) containing protein, Lipop: Lipoprotein, RNAP: RNA polymerase, GST: Glutathione S transferase, SDR: Short-chain dehydrogenase/reductase, Hist. kinase: Histidine kinase, ORF: Open reading frame or hypothetical protein, *tetR*: Tetracycline regulator, Aden. cycl: Adenylate cyclase.

Amino acid comparison of the cytochrome P450 (CitB) homologs, emphasized highly conserved residues, especially the heme-binding domain motif FxxGxRxCxG(S) *(31)* and the ExxR motif in the k-helix, which is important for the stabilization of the meander loop and the tertiary structure of the cytochrome *(32)*. Additionally it became clear that the translation most probably starts with the amino acid sequence motif MP/S(QVR/TAL)LP, whereas the *C*-terminal end of the sequence shows decreasing similarity with higher variability. Comparison of MEGABLAST search results showed that the sequence is conserved. The primary amino acid sequence of the cytochrome P450s of any of those nine strains was used to search in the RCSB PDB database for structurally elucidated homologous proteins. The hit with the highest similarity (27 % identity for MCy9151-CitB) was found for the albaflavenone monooxygenase deriving from *Streptomyces coelicolor* A3(2) *(33)*. The albaflavenone monooxygenase is catalyzing two steps in the biosynthesis, first the hydroxylation of epi-isozizaene to the individual epimers of albaflavenol and second the oxidation of both epimers to the same final antibiotic albaflavenone. Interestingly within the typical cytochrome P450 structure the active site of the functional farnesene synthase is located, catalyzing the conversion from farnesyl diphosphate to *E*)-β-farnesene (61%), (3*E*,6*E*)-α-farnesene (26%), (3*Z*,6*E*)-α-farnesene (6.8%), as well as nerolidol (4.9%) and farnesol (1.8%). However the two putative $Mg^{2+}$ binding motifs the aspartate-rich sequence DD*XX*(D/E) and the residues (N/D)D*XX*(S/T)*XXX*E, (NSE or DTE triad), which are conserved in all terpene synthases of microbial and plant origin could not be found in any of the cytochrome P450 homologs *(34)*. For the recombinant overexpression of the cytochrome P450 gene from the strains MCy9151, MCy9171 and MCy8337 the translation start was set earlier than the sequence alignment suggested. One reason for this approach is the low increase of the 3$^{rd}$-letter GC content across the conserved residues.

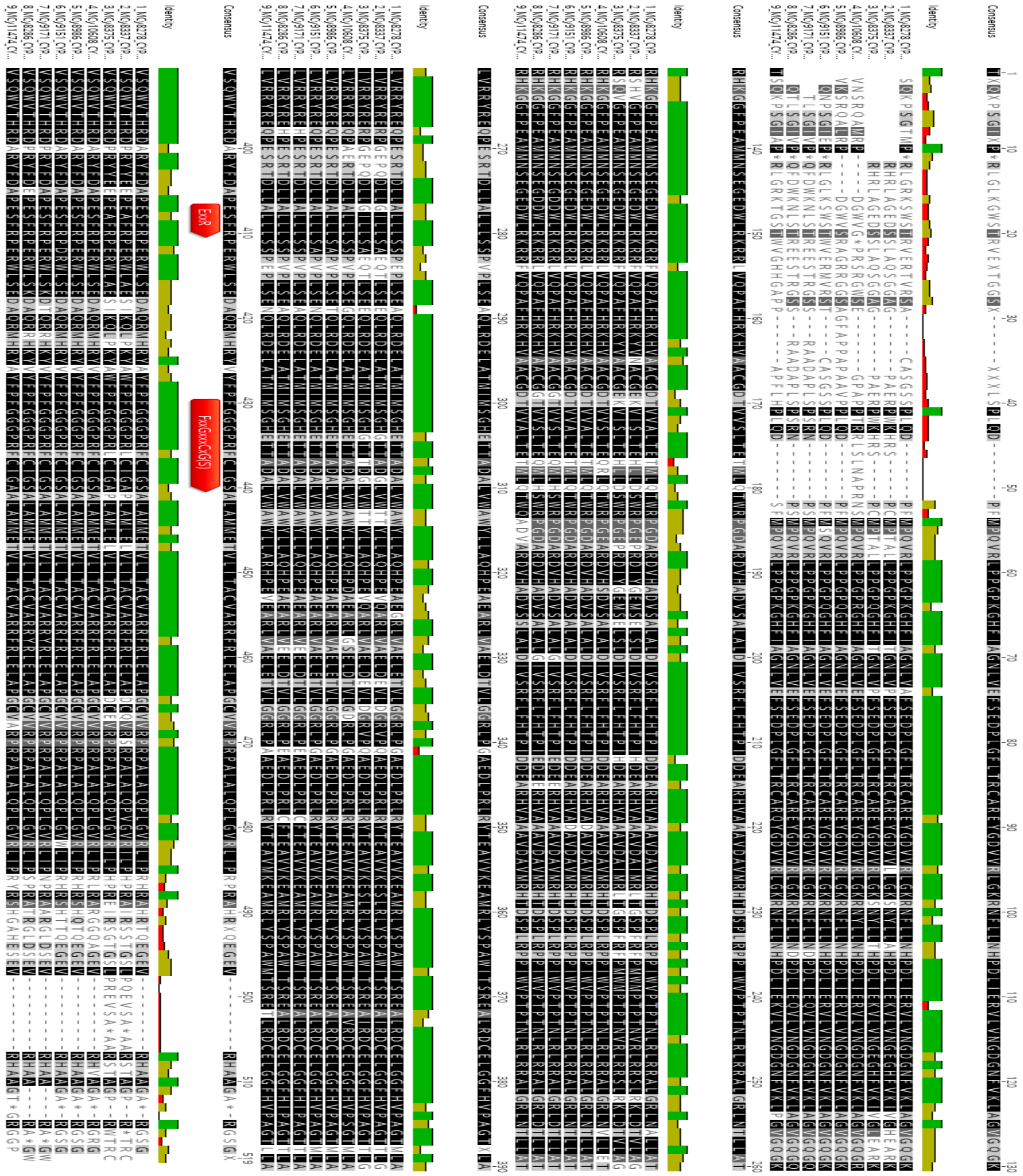

**Fig. S6** Protein alignment of CitB homologs from nine confirmed cittilin producer strains. The comparison shows clearly that the translation most probably starts with the amino acid sequence motif MP/S(QVR/TAL)LP. At the *C*-terminal end of the sequence, the similarity is decreasing significantly and the variability is relatively high.

#### Putative cittilin derivatives

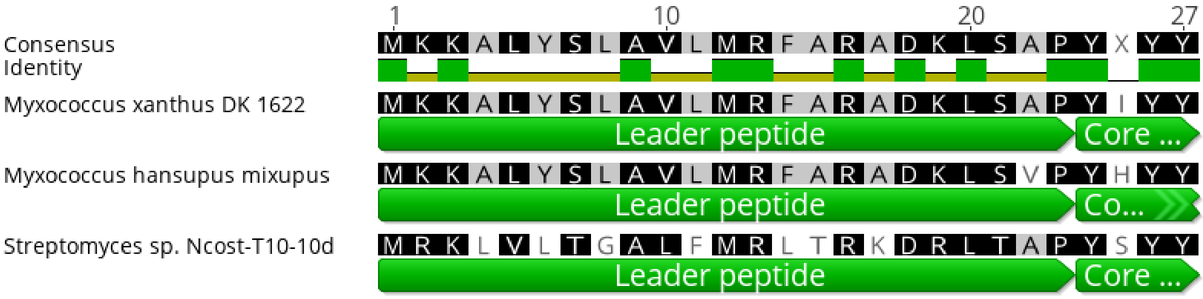

**Fig. S7** Protein alignment of precursor peptide sequences of putative cittilin derivative producers. The leader peptide amino acid sequence of *M. xanthus* DK1622 and *M. hansupus mixupus* are almost identical except for the motif DKLS**A**PY (*M. hansupus mixupus* DKLS**V**PY), whereas the leader peptide amino acid sequence of *Streptomyces* sp. Ncost-T10 shows more dissimilarities.

### Biosynthetic investigation of the cittilin biosynthesis in the native producer

#### Generation of disruption and induced-gene constructs to connect the identified genes to cittilin production

A 631–1021 bp homology sequence of the genes encoding the ABC transporter (only *citT<sub>a</sub>*, to abolish tetrameric ABC transporter expression), the cytochrome P450 (*citB*), the methyltransferase (*citC*) and the prolyl endopeptidase (*mx_pep*) has been PCR amplified by the primers as shown in **Tab. S2**. The specific homology sequence was subcloned via conventional restriction ligation into the pCR2.1 vector from the TOPO-TA cloning kit (Thermo Scientific™ TOPO-TA cloning Kit). Similarly to the creation of disruption constructs, a 1200 bp homology sequence starting from the translational start (not the RBS, but the coding sequence) of the genes encoding $CitT_a$ or the identified precursor peptide CitA has been PCR amplified by the primers as shown in in **Tab. S2**. The homology sequence was subcloned via conventional restriction ligation into the $pFP_{Van}$ vector, which has been constructed and utilized previously to express several genes in myxobacteria *(35,17,36)* The $pFP_{Van}$ vector is a derivative of the pCR2.1 vector featuring a vanillate-inducible promoter which is fused to the vanillate-responsive repressor as described in literature *(37)*.

#### Transfer and chromosomal integration of the constructs into the host *Myxococcus xanthus* DK1622

According to a previously established electroporation procedure for *Myxococcus xanthus* DK1622 *(38)* the strain *M. xanthus* DK1622 was transformed with the generated disruption and induced-gene constructs (**Tab. S6**, genetic constructs 1–6). *M. xanthus* DK1622 transformants were routinely cultivated at 30 °C in CTT medium or CTT agar. Liquid cultures were grown in Erlenmeyer flasks on an orbital shaker at 180 rpm for 3–6 days. *M. xanthus* DK1622 transformants were selected by adding 50 µg/µL kanamycin to the fermentation culture. Correct chromosomal integration of the expression constructs via homologous recombination into the site-specific locus was confirmed by PCR (**Fig. S8**). PCRs were performed according to the settings described above. Genomic DNA of the transformants were isolated using the Gentra® Puregene® Yeast/Bact. Genomic DNA Purification Kit (Qiagen) according to manufacturer's instructions. For each expression construct, correct chromosomal integration was confirmed using two different primer combinations revealing PCR products of the expected sizes:

- Construct No. 1, Seq. primer No.1/3 (1622 bp), and primer No.4/2 (1548 bp)
- Construct No. 2, Seq. primer No.5/7 (1499 bp), and primer No.8/5 (1719 bp)
- Construct No. 3, Seq. primer No.9/11 (1482 bp), and primer No.12/10 (1612 bp)

- Construct No. 4, Seq. primer No.13/15 (1547 bp), and primer No.16/14 (1542 bp)
- Construct No. 5, Seq. primer No.17/19 (1431 bp), and primer No.20/18 (1499 bp)
- Construct No. 6, Seq. primer No.21/23 (1477 bp), and primer No.24/22 (1373 bp)

Genomic DNA of *M. xanthus* DK1622 was used as negative control. A complementary experiment using the following primer combinations revealed a specific PCR product for *M. xanthus* DK1622, but not for any of the transformants of *M. xanthus* DK1622 harboring one of the generated constructs.

- Construct No.1, Seq. primer No. 1/2 (1515 bp PCR product for *M. xanthus* DK1622 wild type)
- Construct No.2, Seq. primer No. 5/6 (1482 bp PCR product for *M. xanthus* DK1622 wild type)
- Construct No.3, Seq. primer No. 9/10 (1597 bp PCR product for *M. xanthus* DK1622 wild type)
- Construct No.4, Seq. primer No. 13/14 (1343 bp PCR product for *M. xanthus* DK1622 wild type)
- Construct No.5, Seq. primer No. 17/18 (1294 bp PCR product for *M. xanthus* DK1622 wild type)
- Construct No.6, Seq. primer No. 21/22 (1425 bp PCR product for *M. xanthus* DK1622 wild type)

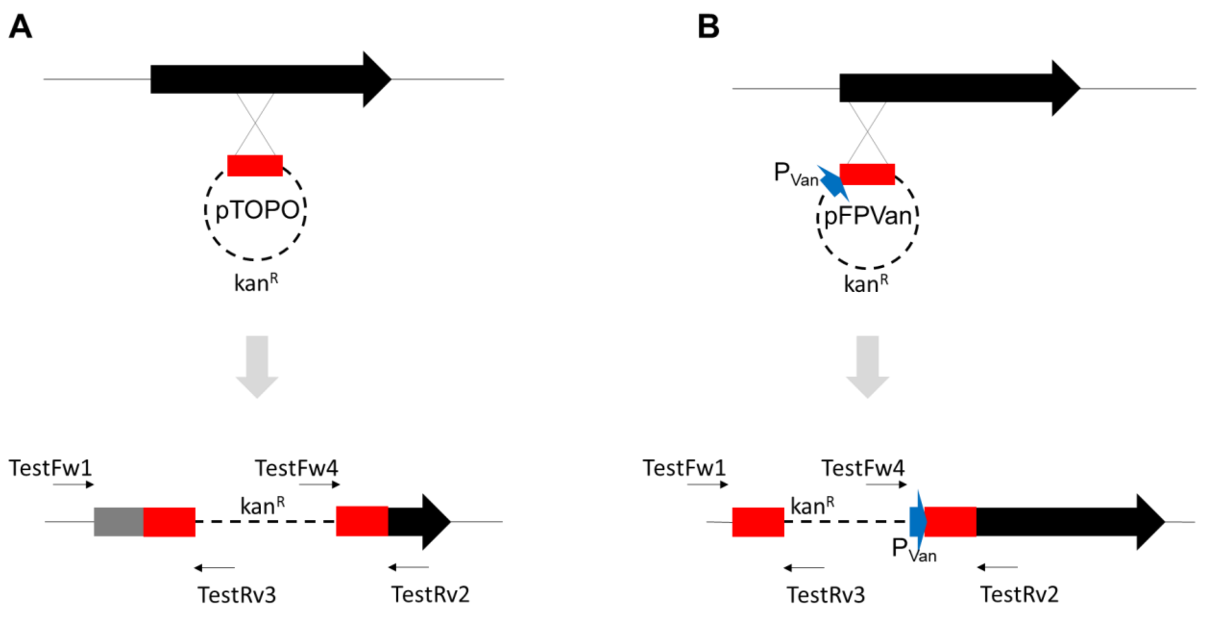

**Fig. S8** Genotypic verification procedure of correct chromosomal integration via PCR. Genetic disruption (**A**) and induced-gene expression constructs (**B**) were verified by multiplex PCR using four primers: two integrations site-specific primers (TestFw1 and TestRv2) and two vector specific primers (Test Rv3 and Test Fw4).

### Heterologous production of cittilin

#### Generation of the expression vector pSET152_$P_{ermE*}$

To express the cittilin BGC in *Streptomyces spp.*, an expression vector was constructed based on the plasmid pSET152. The native promoter sequence of pSET152 was exchanged by the ermE* promoter derived from *Saccharopolyspora erythraea* with nested deletions, finally to achieve higher promoter activity in *Streptomyces lividans* TK24 *(39)*. Therefore the backbone of the generated expression construct derives from the vector pSET152, whereas the ermE* promoter sequence was amplified via PCR from the plasmid pAB03 (Tab. S5, PCR construct 7), with additional restriction enzyme specific sequences for PvuI and XbaI. Afterwards the PCR-amplified ermE* promoter sequence was subcloned into the PacI (the compatible cohesive end of PvuI generated at this position the new restriction site for MseI) and XbaI site of pSET152 to generate the expression vector pSET152_$P_{ermE*}$ (Tab. S6, genetic construct 7).

#### PCR-based cloning of cittilin operon

The cittilin operon consisting of the genes *citA–citC* has been PCR-amplified (Tab. S5, PCR construct 8) and subcloned into the constructed expression vector pSET152_$P_{ermE*}$ via the restriction site NdeI and BspTI yielding the expression vector pSET152_$P_{ermE*}$_cittilin_operon (Tab. S6, genetic construct 8). For the construct including additionally *mx pep*, the cloning strategy was slightly changed. The cittilin operon *citA–citC* was PCR-amplified with a different reverse (Rv) primer harboring the restriction site Bsp1407I (Tab. S5, PCR construct 9), while *mx pep* was PCR-amplified with the restriction sites Bsp1407I and BspTI (Tab. S5, PCR construct 10). Subsequently both PCR amplified inserts were subcloned simultaneously into the constructed expression vector pSET152_$P_{ermE*}$, via the restriction site NdeI and BspTI yielding the expression vector pSET152_$P_{ermE*}$_*citA–citC*_operon_*mx_pep* (Tab. S6, genetic construct 9).

#### Intergeneric conjugation of generated genetic constructs

The generated genetic constructs were transformed for intergeneric conjugation with *Streptomyces albus* del14 into the methylation deficient *E. coli* ET12567 strain harboring the RK2 derivative pUZ8002, since many streptomycetes possess a potent methylation-specific restriction system, which effectively prevents the introduction of heterologous DNA *(40)* (see above). The plasmid pUZ8002 supplies transfer functions to the oriT carrying plasmid backbone of pSET152 without transferring itself due to a mutation in its own oriT *(41)*. After conjugation, the generated mutants *Streptomyces albus* del14_pSET152_$P_{ermE*}$, *Streptomyces albus* del14_pSET152_$P_{ermE*}$_*citA–citC*_operon and *Streptomyces albus* del14_pSET152_$P_{ermE*}$_*citA–citC*_operon_*mx_pep* were used from a sporulating agar plate to inoculated 20 mL of TSB medium to obtain seed cultures. After three days, the seed cultures were well grown and used to inoculate with 1–2 mL COM medium containing XAD-16. Four days later, the cultures were centrifuged, CPs and XAD-16 extracted (50% methanol/ 50% acetone (v:v)) and measured via HPLC-MS. As reference, a crude extract from *M. xanthus* DK1622 and a pure reference of cittilin A was used. Cittilin A production was observed in the mutants *Streptomyces albus* del14_pSET152_P$_{ermE*}$_*citA–citC*_operon and *Streptomyces albus* del14_pSET152_P$_{ermE*}$_*citA–citC*_operon_*mx_pep*. Furthermore, MS²-spectra and fragmentation pattern confirmed the identity of heterologously produced cittilin A (Fig. S9, Fig. S10).

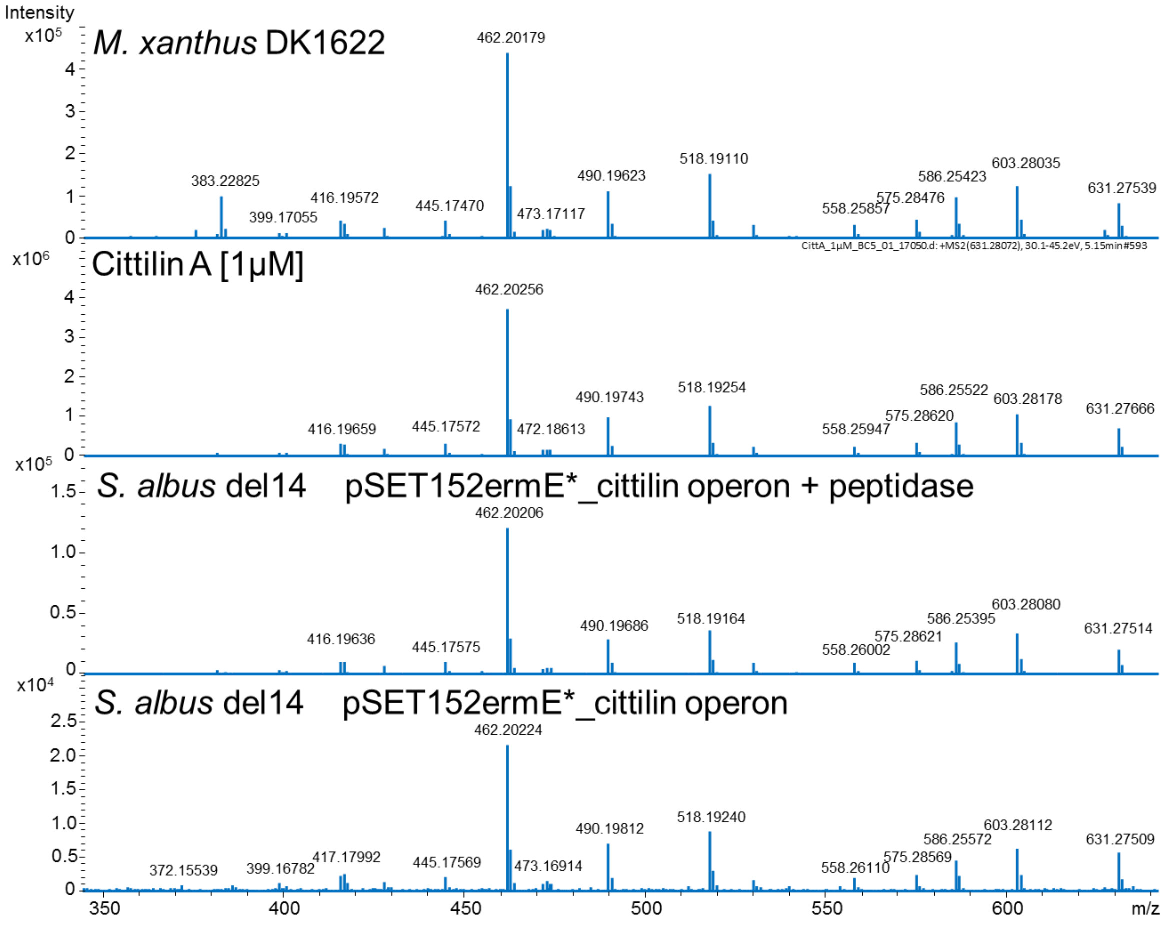

**Fig. S9** MS²-spectra of cittilin A. Cittilin A produced by *M. xanthus* DK1622, cittilin A pure compound as reference and the heterologous produced cittilin A in *Streptomyces albus* del14 pSET152ermE*_cittilin operon with and without coproduction of MX PEP. All four MS²-spectra exhibit the same fragmentation pattern, which confirms the successful heterologous production of cittilin A. Due to the selective MS/MS fragmentation it was not possible to show a time interval of the MS/MS fragmentation pattern.

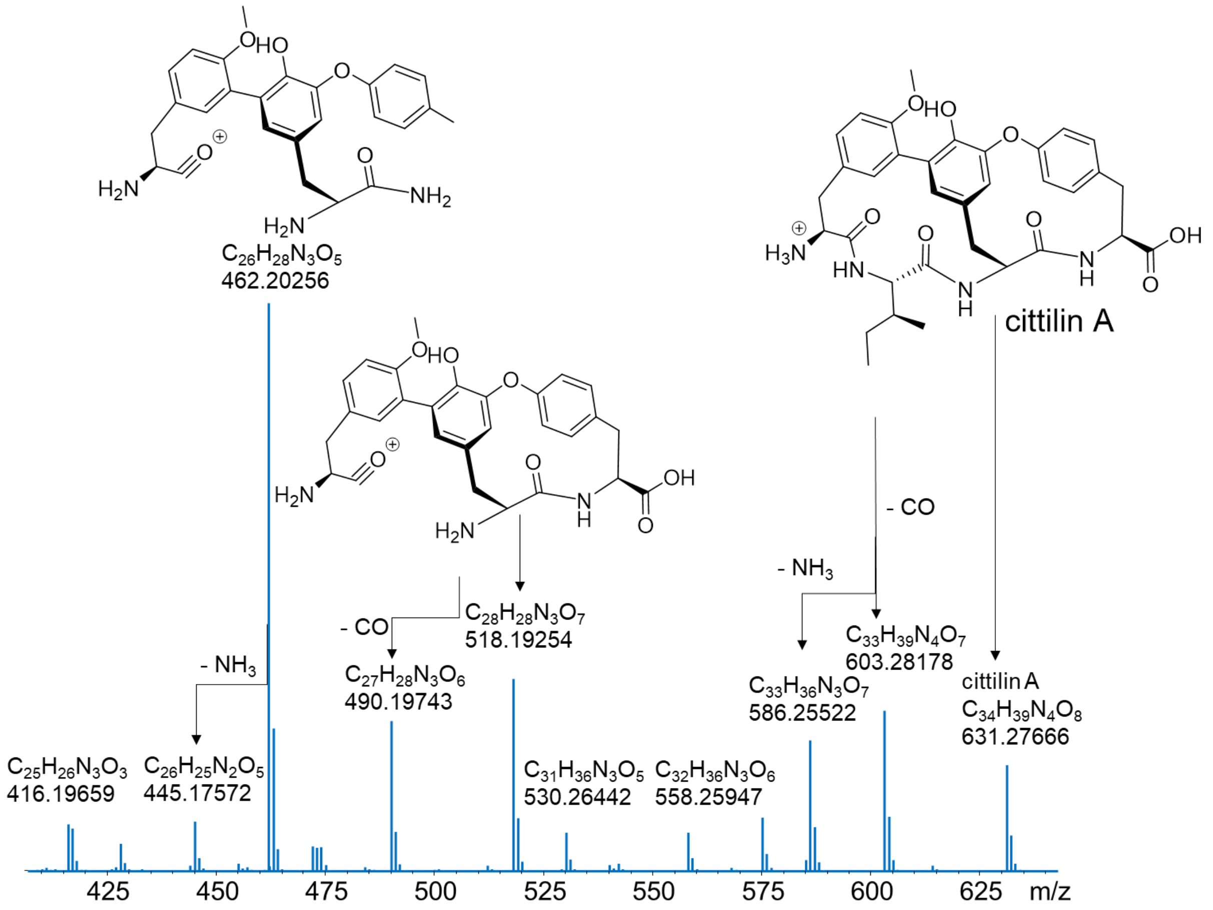

**Fig. S10** $MS^2$-spectrum and fragmentation pattern of cittilin A.

### *In vitro* and cell free lysate investigation of cittilin biosynthesis

#### *In vitro* investigation of CitC

##### Cloning of *citC* into overexpression vector

The gene encoding the cittilin-specific methyltransferase CitC in *M. xanthus* DK1622 has been PCR-amplified by the primers as shown in **Tab. S2**. The amplified DNA fragment encoding CitC was subcloned into the expression vector pET28b via the restriction site NdeI and EcoRI yielding the expression vector pET28b_DK1622_CitC (**Tab. S6**, genetic construct 10)

##### *E. coli*-based recombinant CitC production

The generated expression vector pET28b_DK1622_CitC was transformed into *E. coli* Lemo21 for the recombinant production of CitC containing the *N*-terminal $His_6$-tag and Tobacco etch virus (TEV) cleavage site and an IPTG-inducible *lac*-operon. The generated *E. coli* strain was grown o/n in LB medium containing 50 µg/mL kanamycin at 37 °C. Antibiotic supplemented LB medium was inoculated with the o/n culture and incubated at 37 °C until an $OD_{600}$ of 0.6 was reached. The culture was equilibrated at 16 °C before expression of $His_6$-tagged *citC* was induced with 1 mM IPTG. The culture was incubated at 16 °C for 20 h. Subsequently the cells were harvested at 3400 g for 10 min at 4 °C. The CP of *E. coli* Lemo21_ pET28b_DK1622_CitC was re-suspended in lysis buffer and cells were lysed using a CD-017a constant cell disruption system. Cell debris was removed by centrifugation (15 min at 50000 g) and the SN was loaded to a 5 mL HisTrap FF column (GE Healthcare) with 5 mL/min on an ÄKTA™ pure system (GE Healthcare) after the column was equilibrated with five column volumes (CVs) lysis buffer. The column loaded with recombinant protein was washed with 10 CV lysis buffer. Elution was performed isocratically with 5 CV elution buffer at a flow rate of 5 mL/min. Protein-containing fractions, protein identity and purity were assessed by SDS-PAGE analysis (12% acrylamide) and LC-MS. Combined fractions containing CitC were concentrated via centrifugal filtration using Amicon Ultra-30 columns (MW 30000 Da, Merck). Size exclusion chromatography was performed using a Superdex 200 Increase prepacked column. After equilibration with 1.2 CV protein buffer, the recombinant protein solution was loaded via a 2 mL loading loop. The size excluded protein fractions were identified via SDS-PAGE (Fig. S11). The combined protein fraction was digested with TEV protease (1.5 mg/10 mg recombinant protein) (o/n) at 4 °C. The next day, a second Ni-affinity purification was performed using a 5 mL HisTrap FF column (GE Healthcare) with 5 mL/min on an ÄKTA™ pure system (GE Healthcare). The column was equilibrated with 5 CV lysis buffer and the TEV digested protein solution was loaded on the HisTrap FF column with an flow rate of 5 mL/min and 10 CV column wash followed. The column with bound $His_6$-TEV protease and $His_6$-TEV site was isocratically eluted with 5 CV elution buffer. The column wash fractions with CitC without $His_6$-tag were combined and protein identity was assessed by SDS-PAGE. The protein solution was concentrated via centrifugal filtration using Amicon Ultra-30 columns (MW 30000 Da, Merck). Size exclusion chromatography followed using a Superdex 200 Increase prepacked columns. After equilibration with 1.2 CV protein buffer with a flow rate of 1 mL/min the recombinant protein solution was loaded via a 2 mL loading loop. Protein-containing fractions, protein identity and purity were assessed by SDS-PAGE and LC-MS. Combined CitC fractions were concentrated via centrifugal filtration using Amicon Ultra-30 columns (MW 30000 Da, Merck). The concentrated CitC solution was adjusted at 10% glycerol and 100 µM and aliquoted into PCR tubes. Protein concentrations were determined by UV spectroscopy (with ε280 nm values) using Thermo Scientific™ NanoDrop™ 2000/2000c. The aliquots were frozen immediately in liquid nitrogen and stored at -80 °C.

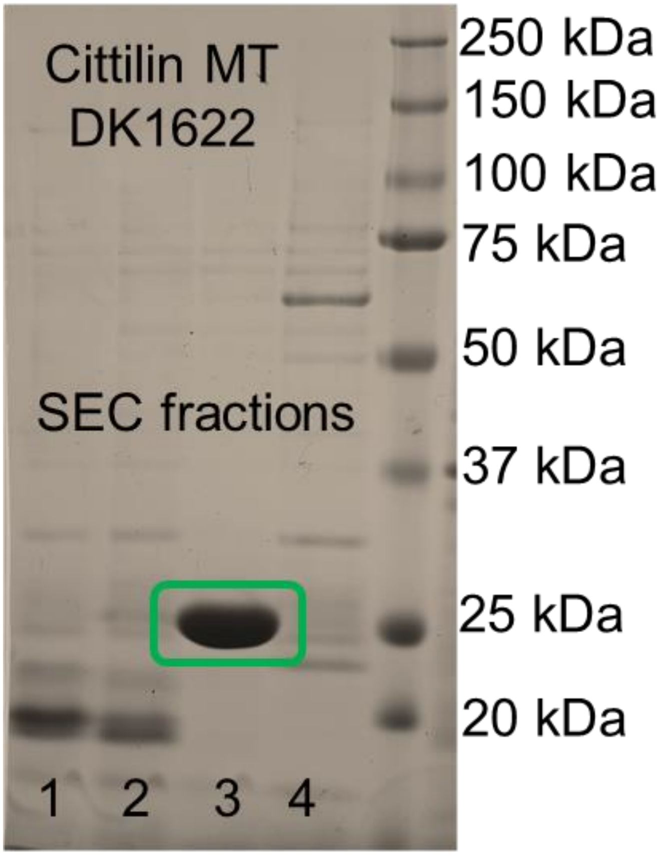

**Fig. S11** SDS-PAGE gel of CitC purification. Green circle indicate CitC after size exclusion chromatography.

#### Catalytic activity testing of recombinant CitC

Catalytic activity of the purified recombinant CitC was tested in a reaction mixture (50 µL end volume) containing 2 µM CitC, 2 mM *S*-adenosyl-L-methionine (SAM), 5 µL of concentrated crude extract (see below) in MQ water. Negative control testing were performed by omitting SAM or CitC (data not shown). Replacing concentrated crude extract with 100 µM precursor ((synthesized and commercially purchased peptide with the sequence KKALYSLAVLMRFARADKLSAPYIYY (DK1622 motif)) peptide or core peptide (synthesized and commercially purchased peptide with the sequence YIYY) showed no methylation of these substrates. The reaction was carried out o/n at 30°C. The reaction was terminated by adding MeOH (50 µL, final concentration 50% v/v). The mixture was transferred to -80 °C for at least 1 h, centrifuged at 13000 g for 15 min at 4°C (VWR centrifuge ECN521-3601, Hitachi Koki Co., Ltd) and 1 µL of the SN was subjected to HPLC-MS analysis as described previously.

#### Preparation of cittilin B as substrate from crude extract

One mL of crude extract from the mutant *M. xanthus* DK1622_*citC*_disruption (obtained previously see above) was dried under $N_2$. Residue was re-dissolved with 100 µL MeOH. Centrifugation for 15 min at 15000 rpm, 4 °C. The SN was used subsequently as substrate (cittilin B).

### *In vitro* investigation of the prolyl endopeptidase MX PEP

#### Cloning of the *mx pep* into overexpression vector

The gene encoding the prolyl endopeptidase MX PEP in *M. xanthus* DK 1622 has been PCR-amplified by the primers as shown in **Tab. S2**. The amplified DNA fragment encoding MX PEP was subcloned into the expression vector pET28b via the restriction site NdeI and HindIII, yielding the expression vector pET28b_DK1622_MX_PEP (**Tab. S6**, genetic construct 11).

#### *E. coli*-based recombinant MX PEP production

The generated expression vector pET28b_DK1622_MX_PEP was transformed into *E. coli* BL21 (λDE3) for the recombinant production of MX PEP containing the *N*-terminal $His_6$-tag and TEV cleavage site and an IPTG-inducible *lac*-operon. The *E. coli* host harboring the expression vector was grown o/n in LB medium containing 50 µg/mL kanamycin at 37 °C. Antibiotic supplemented LB medium was inoculated with the o/n culture and incubated at 37 °C until an $OD_{600}$ of 0.6 was reached. The culture was equilibrated at 16 °C before the expression of $His_6$-tagged *mx pep* was induced with 1 mM IPTG. The culture was incubated at 16 °C for 20 h. Subsequently the cells were harvested at 3400 g for 10 min at 4 °C. The CP of *E. coli* BL21 (λDE3) with pET28b_DK1622_MX_PEP was re-suspended in lysis buffer and cells were lysed using a CD-017a constant cell disruption system. Cell debris was removed by centrifugation (15 min at 50000 g) and the SN was loaded to a 5 mL HisTrap FF column (GE Healthcare) with 5 mL/min on an ÄKTA™ pure system (GE Healthcare) after the column was equilibrated with 5 CV lysis buffer. The column loaded with recombinant protein was washed with 10 CV lysis buffer. Elution was performed via a linear gradient up to 100%, with 5 CV elution buffer at a flow rate of 5 mL/min. Protein-containing fractions, protein identity and purity were assessed by SDS-PAGE analysis (12% acrylamide) and LC-MS. Combined fractions containing MX PEP were concentrated via centrifugal filtration using Amicon Ultra-30 columns (MW 50000 Da, Merck). Size exclusion chromatography was performed using a Superdex 200 Increase prepacked column. After equilibration with 1.2 CV protein buffer, the recombinant protein solution was loaded via a 2 mL loading loop. The size excluded protein fractions were identified via SDS-PAGE (Fig. S12). Combined MX PEP fractions were concentrated via centrifugal filtration using Amicon Ultra-30 columns (MW 30000 Da, Merck). The concentrated protein solution was adjusted to 10% glycerol and 100 µM and aliquoted into PCR tubes. Protein concentrations were determined by UV spectroscopy (with ε280 nm values) using Thermo Scientific™ NanoDrop™ 2000/2000c. The aliquots were frozen immediately in liquid nitrogen and stored at -80 °C.

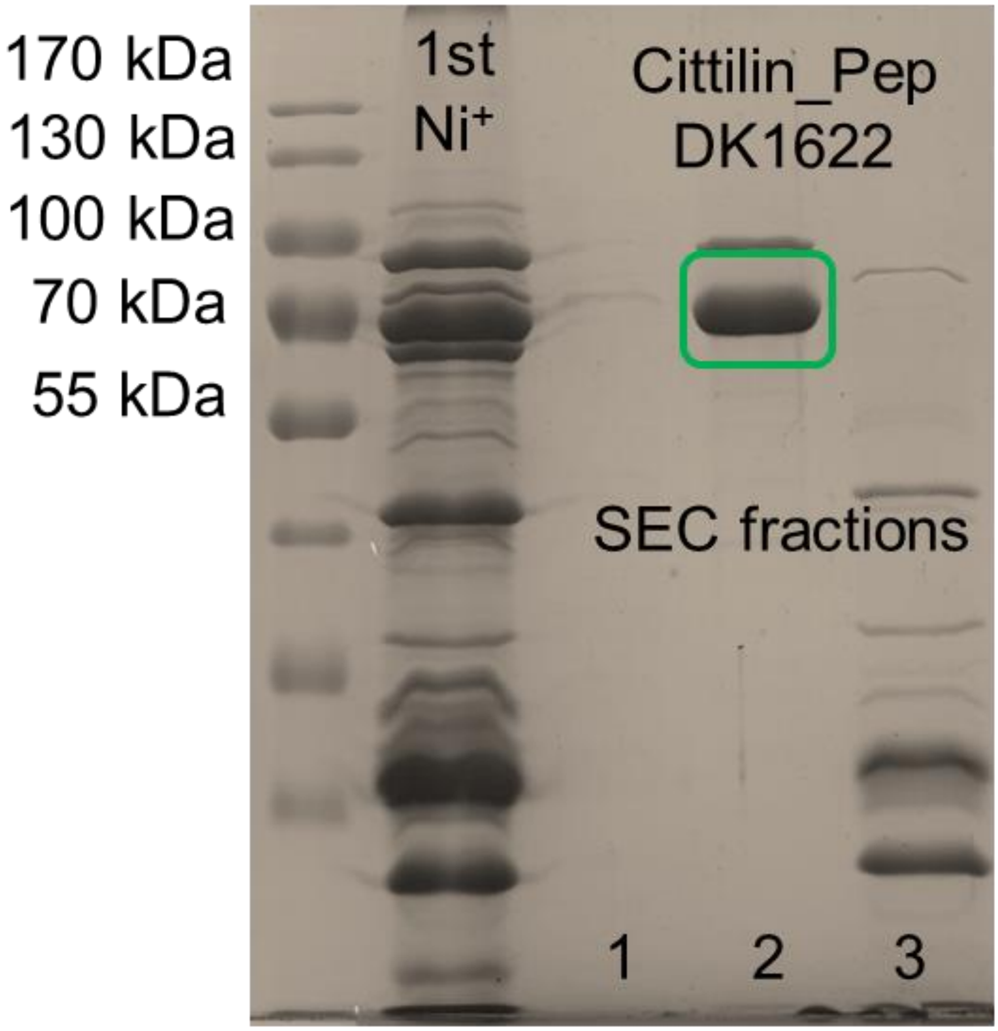

**Fig. S12:** SDS-PAGE gel of MX PEP purification. Green box indicates recombinant MX PEP after size exclusion chromatography (SEC) fractionation.

#### Catalytic activity testing recombinant MX PEP

Since the modified precursor peptide was not available as a substrate for MX PEP, we decided to first investigate conversion of the unmodified precursor peptide. Catalytic activity of the purified recombinant MX PEP was tested in a reaction mixture containing 5 µM MX PEP and 100 µM precursor peptide (synthesized and commercially purchased peptide with the sequence KKALYSLAVLMRFARADKLSAPYIYY (DK1622 motif) in 50 mM NaCl, 20 mM Bis-TRIS buffer (pH 6.8). Negative control testing were performed by omitting recombinant MX PEP. The reaction was carried out (o/n) at room temperature. The reaction was terminated by adding MeOH (final concentration 50% v/v). The mixture was transferred to -80 °C for at least 1 h, centrifuged at 13000 g for 15 min at 4 °C (VWR centrifuge ECN521-3601, Hitachi Koki Co., Ltd) and 1 µL of the SN was subjected to HPLC-MS analysis as described previously.

The recombinantly produced prolyl endopeptidase originating from *M. xanthus* DK1622 did catalyze the cleavage of the leader peptide yielding the linear core peptide, which was confirmed by mass spectrometric detection of a 621.28 Da fragment (**Fig. S13**). In conclusion, the identified prolyl endopeptidase catalyzes the cleavage of different peptides as reported previously *(42,43)*, and is not exclusively optimized for the cleavage of the modified or native cittilin precursor peptide.

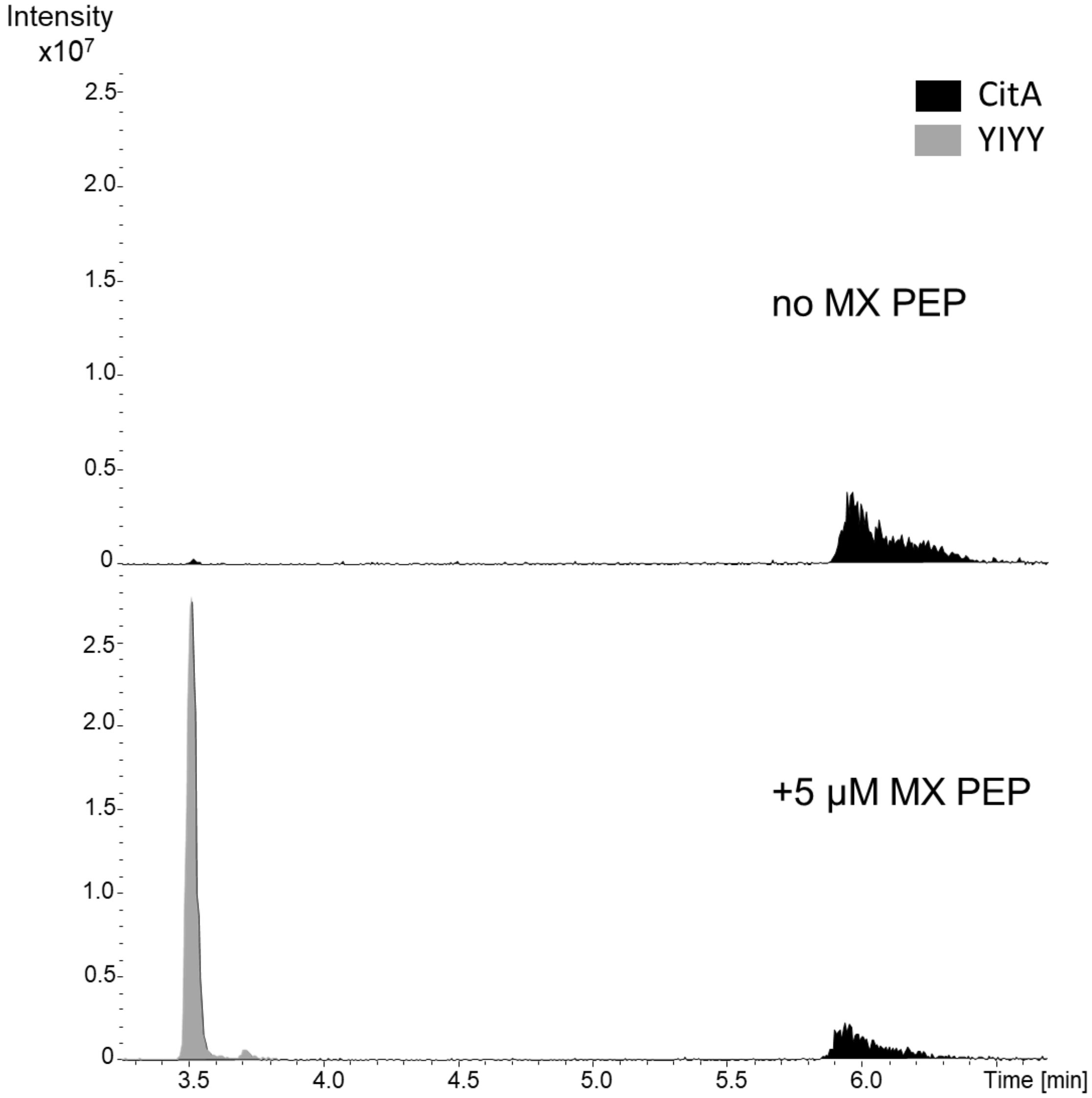

**Fig. S13:** HPLC-MS EIC chromatograms (3051.75 m/z (grey) and 621.28 m/z (black), precursor peptide [M+H] and core peptide [M+H]) shows the catalytic cleavage of the precursor peptide into the core peptide via recombinantly produced prolyl endopeptidase**.**

### *In vitro* investigation of the cittilin cytochrome P450 enzyme CitB

#### Cloning of *citB* into overexpression vector for *E. coli*-based expression

The gene encoding the cittilin cytochrome P450 CitB in *M. xanthus* DK 1622 has been PCR-amplified by the primers as shown in Tab. S2. The amplified DNA fragment encoding $CitB_{DK1622}$ was subcloned into the expression vector pHisTEV via the restriction site NcoI and HindIII, yielding the expression vector pHisTEV_DK1622_CitB (**Tab. S6**, genetic construct 12).

#### *E. coli*-based recombinant $CitB_{DK1622}$ production

Recombinant production of CitB was coupled to *in vivo* co-production of chaperons, using recombinant vectors from a commercially available chaperone plasmid set (Takara Bio Inc.). The generated recombinant expression vector pHisTEV_DK1622_CitB and pGro7 (Takara Chaperone plasmid set) for co-production of GroEL/GroES were co-transformed into *E. coli* C43. *E. coli* C43 with pHisTEV_DK1622_CitB and pGro7 was grown o/n in LB medium containing 50 µg/mL kanamycin and 25 µg/mL chloramphenicol at 37 °C. Antibiotic supplemented TB medium was inoculated with the (o/n) culture and incubated at 37 °C until an $OD_{600}$ of 0.6 was reached. The culture was equilibrated at 16 °C before the expression of $His_6$-tagged *citB*$_{DK1622}$ was induced with 0.5 mM IPTG and co-expression of *groEL/groES* with 3 mg/mL L-arabinose. The culture was incubated at 16 °C for 18 h. Subsequently, cells were harvested at 3400 g for 10 min at 4 °C. The CP of *E. coli* C43 with pHisTEV_DK1622_CitB and pGro7 was re-suspended in lysis buffer and cells were lysed using a CD-017a constant cell disruption system. Cell debris was removed by centrifugation (15 min at 50000 g) and the SN was loaded to a 5 mL HisTrap HP column (GE Healthcare) with 5 mL/min on an ÄKTA™ pure system (GE Healthcare) after the column was equilibrated with 5 CV lysis buffer. Subsequently the column with loaded recombinant protein was washed with 10 CV lysis buffer. Elution was performed isocratically with 5 CV elution buffer and a flow rate with 5 mL/min. Protein-containing fractions, protein identity and purity were assessed by SDS-PAGE. Combined $CitB_{DK1622}$ were concentrated via centrifugal filtration using Amicon Ultra-30 columns (MW 30000 Da, Merck). Size exclusion chromatography was performed using a Superdex 200 Increase prepacked columns. After equilibration with 1.2 CV protein buffer, the recombinant protein solution was loaded via a 5 mL loading loop. The size excluded protein fractions were identified via SDS-PAGE. The combined protein fractions were digested with TEV protease (1.5 mg/ 10 mg recombinant protein) o/n at 4 °C. The next day, a second Ni-affinity purification was performed using a 5 mL HisTrap FF column (GE Healthcare) with 5 mL/min on an ÄKTA™ pure system (GE Healthcare). The column was equilibrated with 5 CV lysis and the TEV digested protein solution was loaded on the HisTrap FF column with a flow rate of 5 mL/min and 10 CV column wash followed. The column with bound $His_6$-TEV protease and $His_6$-TEV-site was isocratically eluted with 5 CV elution buffer. The column wash fractions with $CitB_{DK1622}$ without $His_6$-tag were combined and protein identity was assessed by SDS-PAGE. On the gel, the chaperone GroEL is present, referred with the binding of the chaperone to the substrate protein. LC-MS measurements failed due to overlapping peaks in the chromatogram, further it was not possible to deconvolute an exact mass off the protein. The protein solution was concentrated via centrifugal filtration using Amicon Ultra- 30 columns (MW 30,000 Da, Merck). Size exclusion chromatography followed using a Superdex 200 Increase prepacked columns. After equilibration with 1.2 CV protein buffer, the recombinant protein solution was loaded via a 2 mL loading loop. Protein-containing fractions, protein identity and purity were assessed by SDS-PAGE (**Fig. S14**). Combined $CitB_{DK1622}$ fractions were concentrated via centrifugal filtration using Amicon Ultra-30 columns (MW 30000 Da, Merck). The concentrated protein solution was adjusted at 10% glycerol and 100 µM and aliquoted into PCR tubes. The aliquots were frozen immediately in liquid nitrogen and stored at -80 °C. Protein concentrations were determined by UV spectroscopy (with ε280 nm values) using Thermo Scientific™ NanoDrop™ 2000/2000c. The amount of protein was high, related to the high amount of recombinant chaperon bound to the substrate protein.

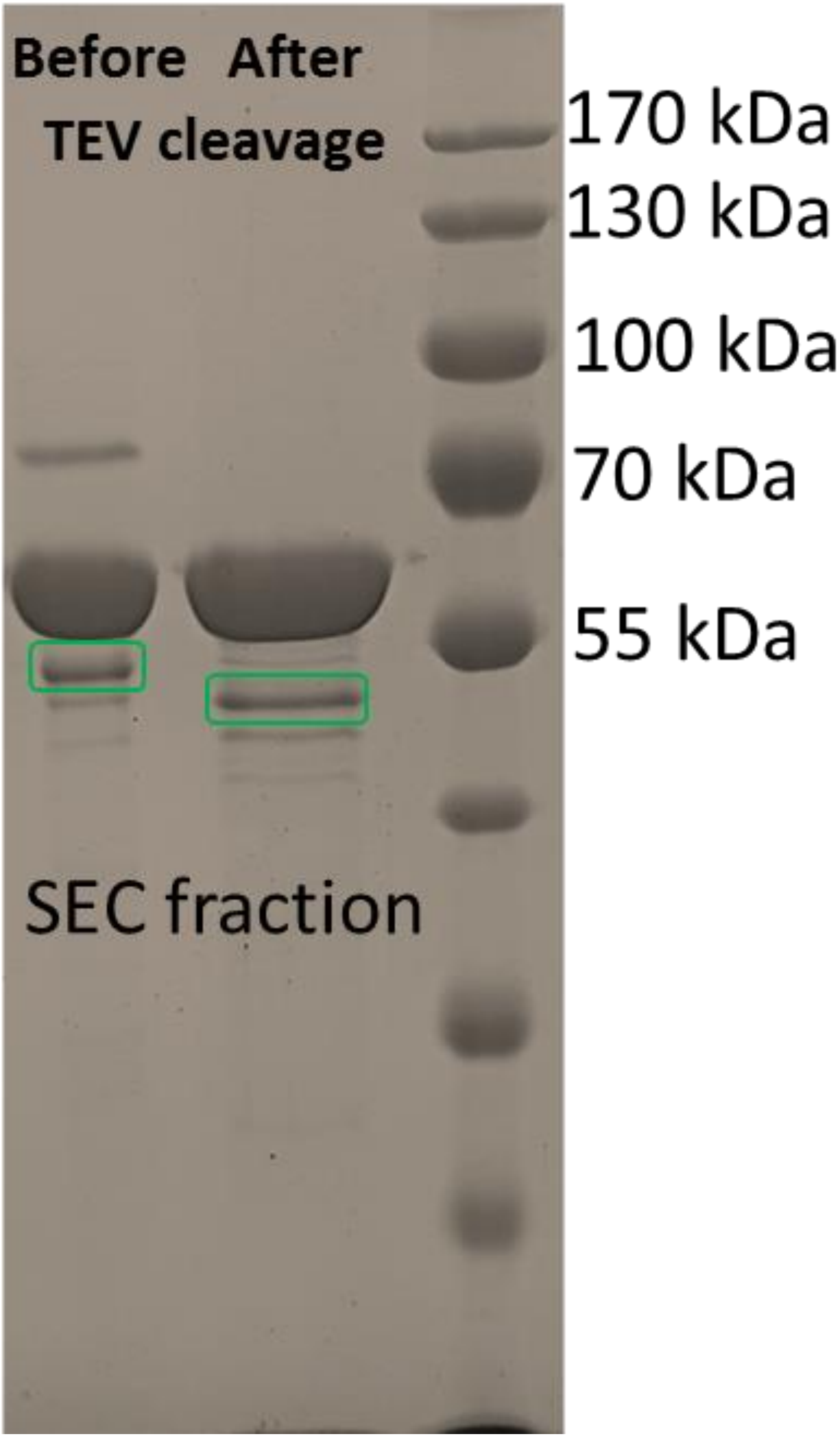

**Fig. S14:** SDS-PAGE gel of recombinant $CitB_{DK1622}$ purification. Green rectangles indicate pure $CitB_{DK1622}$ (left; with $His_6$-TEV-site, right; without $His_6$-TEV-site), whereas the upper bands indicate co-produced GroEL.

#### Catalytic activity testing of recombinant $CitB_{DK1622}$ from *E. coli* based production

Catalytic activity of the recombinant $CitB_{DK1622}$ was tested in a reaction mixture containing approx. 5 µM $CitB_{DK1622}$ and 100 µM precursor peptide (synthesized and commercially purchased peptide with the sequence KKALYSLAVLMRFARADKLSAPYIYY (DK1622 motif)) or core peptide (synthesized and commercially purchased peptide with the sequence YIYY). In addition, 500 µM $MgCl_2$, 1 mM ATP and 2 mM of co-factor (one of the following; FMN, $FAD^+$, $NAD^+$,$NADP^+$, NADPH, NADH) in 500 mM NaCl, 20 mM Bis-TRIS buffer (pH 6.8) was added. As alternative approach to achieve co-factor regeneration, the commercially available Fdx/FdR reductase pair system from *Spinacia oleracea* was performed with 2.5 µM Fdx, 2.5 µM FdR and 2 mM NADPH. Control testing were performed by omitting $CitB_{DK1622}$. The reaction was carried out for 10 min, 1 h and (o/n) at room temperature and 30 °C. The reaction was terminated by adding MeOH (final concentration 50% v/v). The mixture was transferred to -80 °C for at least 1 h, centrifuged at 13000 g for 15 min at 4 °C (VWR centrifuge ECN521-3601, Hitachi Koki Co., Ltd) and 1 µL of the SN was subjected to HPLC-MS analysis as described previously. No activity of recombinant $CitB_{DK1622}$ was observed.

#### Cloning of the *citB* into overexpression vector for *Streptomyces*-based expression

Since the $CitB_{DK1622}$ seemed to be recalcitrant for efficient recombinant production in *E. coli*, an alternative strategy was conducted utilizing *Streptomyces* as heterologous host for recombinant production of CitB. The successful heterologous expression of the cittilin BGC in *S. albus* del14 and previous successful recombinant cytochrome P450 enzyme production in *Streptomycetes (44)* underline the potential of this approach.

In order to increase the prospects for successful recombinant protein production, not only *citB* from *M. xanthus* DK1622 was cloned and tested for heterologous expression but also homologs from *Cystobacter* spp. (MCy9171) and Cb vi35 (MCy8337) were cloned. The expression vector pCJW93, which has an *N*-terminal $His_6$-tag and thrombin cleavage site (LVPRGS) was used for recombinant CitB production. Since the available thrombin cleavage site has several disadvantages for later performed purification steps *(45)*, the TEV cleavage site was additionally amplified for *N*-terminal $His_6$-tag constructs to facilitate the cleavage of the *N*-terminal $His_6$-tag during protein purification. In order to test the influence of either an *N*-terminal or *C*-terminal $His_6$-tag for the functional production of cytochrome P450 enzyme, the $His_6$-tag was modified by cloning procedures to obtain the shuttle vector pCJW93_noHis without any $His_6$-tag and thrombin cleavage site, in which *citB* with amplified *C*-terminal $His_6$-tag can be cloned. This modification was conducted through restriction digestion with Alw44I and NdeI of the shuttle vector pCJW93, to yield two different DNA fragments. The smaller DNA fragment was exchanged through a PCR amplified fragment (pCJW93_noHistag_exchange_construct, **Tab. S5**, No. 17), which was digested with the restriction enzymes Alw44I and NdeI. The native larger DNA fragment pCJW93_backbone and the smaller PCR amplified pCJW93no $His_6$-tag fragment were ligated via Alw44I and NdeI to obtain the pCJW93noHis plasmid after transformation of the ligation product into *E. coli* HS996. The shuttle vectors pCJW93 and pCJW93noHis were used for further cloning of different *citB* homologs. In general, all CitB constructs designed for production of recombinant protein with *N*-terminal $His_6$-tag, the PCR products contained the additional TEV cleavage site for further purification steps, whereas constructs designed for production of recombinant protein with *C*-terminal $His_6$-tag, the PCR products had to be amplified with a *C*-terminal $His_6$-tag.

- $citB_{MCy8337}$ from strain Cb vi35 (MCy8337) with *C*-terminal $His_6$-tag and $citB_{MCy9171}$ from MCy9171 with *N*-terminal $His_6$-tag were digested with EcoRI and HindIII and cloned into the EcoRI and HindIII site of pCJW93 and respectively of pCJW93noHis.
- $citB_{DK1622}$ from DK1622 gene with *N*-terminal $His_6$-tag, $citB_{DK1622}$ from DK1622 gene with *C*-terminal $His_6$-tag, $citB_{MCy9171}$ from MCy9171 with *C*-terminal $His_6$-tag and $citB_{vi35}$ from Cb vi35 with *N*-terminal $His_6$-tag was cloned into the *Nde*I and *EcoR*I site of pCJW93 and respectively of pCJW93noHis.

The gene encoding CitB in *M. xanthus DK 1622* and MCy9171 has been PCR-amplified by the primers as shown in Tab. S2. The amplified DNA fragment encoding the CitB was subcloned into the expression vector pCJW93 via the restriction site NdeI and HindIII, yielding the expression vector pCJW93_DK1622_CitB (Tab. S6, genetic construct 13).

##### *Streptomyces*-based recombinant production of CitB; test productions

The generated plasmids were conjugated into *Streptomyces coelicolor* CH999 with *E. coli* ET12567 harboring the plasmid pUZ8002 as donor strain as described previously (see above). After conjugation, the generated *S. coelicolor* CH999 mutants were used to inoculate TSB media to obtain seed cultures. After three days, the seed cultures were used to inoculate duplicates of 100 mL of Super YEME media with 5 mL of the seed culture and were incubated at 30 °C and 180 rpm for at least two days. The cultures were prepared in 250 mL Erlenmeyer flasks with metal spirals to ensure a suspension cell culture. After two days, the cultures were well grown and gene expression was induced with thiostrepton (working concentration 10 µg/mL). The duplicates of cultures were separated into two parts; one was incubated at 25 °C, 180 rpm, the other at 30 °C, 180 rpm. 24 h later, the incubation was stopped and 10 mL of each culture was transferred into a Falcon tube and centrifuged for at 4000 rpm for 10 min at 4 °C. After discarding the SN, the CP was re-suspended in 500 µL lysis buffer No. 3. The cell suspension was sonicated, centrifuged and nickel pulldown was performed (see below). SDS-PAGE revealed that the $citB_{MCy9171}$ from *Cystobacter* MCy9171 could be recombinantly overexpressed according to the described parameters. Fig. S15 and Fig. S16 display a specific band with the size of 50 kDa, which can be connected to soluble $CitB_{MCy9171}$ (Fig. S15) and $CitB_{MCy9171}$ in the inclusion bodies (Fig. S16).

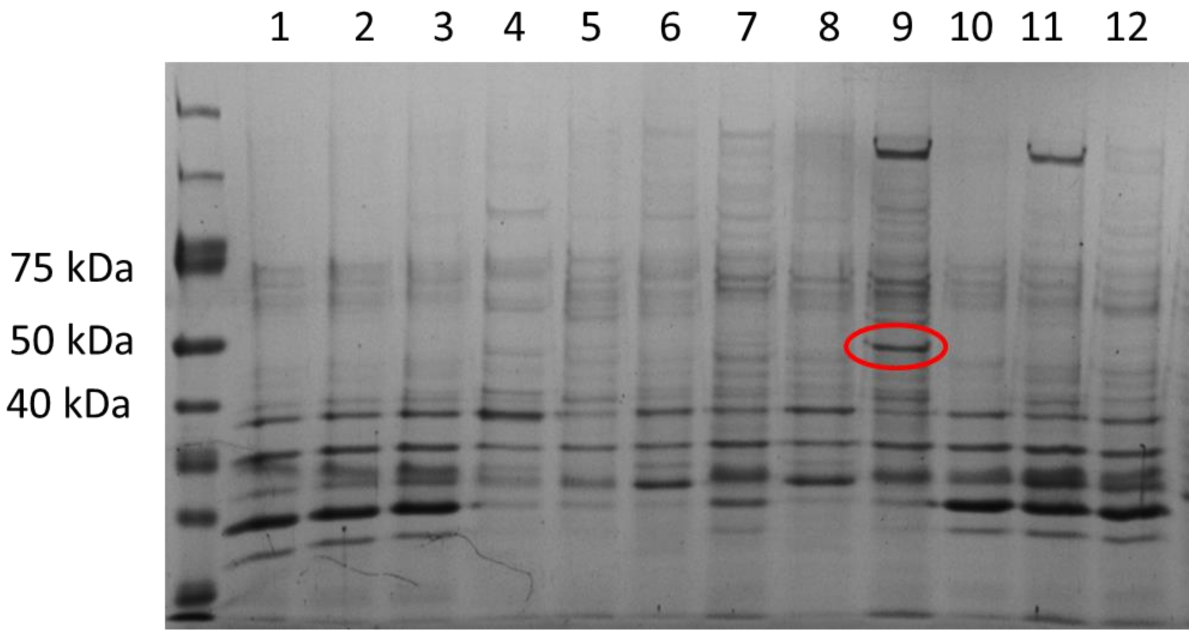

**Fig. S15:** SDS-PAGE of *citB* expression tests in *S. coelicolor* CH999 after nickel pulldown. The temperature of incubation after induction was 25 °C and 30 °C respectively. 1) *S. coelicolor* CH999_pCJW93_DK1622_CitB at 25 °C; 2) *S. coelicolor* CH999_pCJW93noHis_DK1622_CitB at 25 °C; 3) *S. coelicolor* CH999_pCJW93_DK1622_CitB at 30 °C; 4) *S. coelicolor* CH999_pCJW93noHis_DK1622_CitB at 30 °C; 5) *S. coelicolor* CH999 pCJW93_Cb_vi35_CitB at 25 °C; 6) *S. coelicolor* CH999_pCJW93noHis_Cb vi35_CitB at 25 °C; 7) *S. coelicolor* CH999_pCJW93_Cb_vi35_CitB at 30 °C; 8) *S. coelicolor* CH999_pCJW93noHis_Cb_vi35_CitB at 30 °C; 9) *S. coelicolor* CH999_pCJW93_MCy9171_CitB at 25 °C; 10) *S. coelicolor* CH999_pCJW93noHis_MCy9171_CitB at 25 °C; 11) *S. coelicolor* CH999_pCJW93_MCy9171 CitB at 30 °C; 12) *S. coelicolor* CH999_pCJW93noHis_MCy9171_CitB at 30 °C.

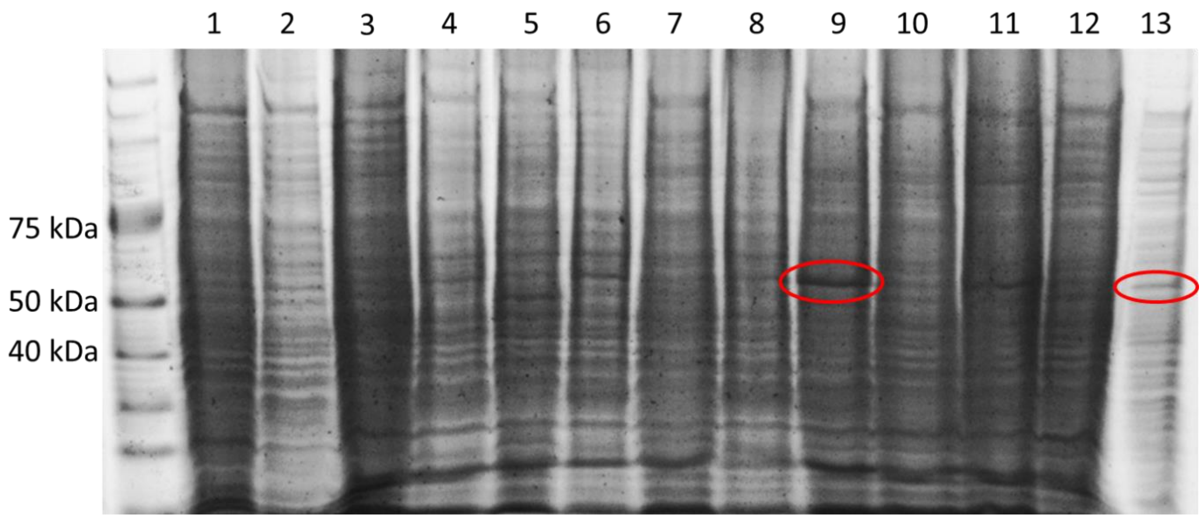

**Fig. S16:** SDS-PAGE (cell pellets (CPs) *citB* expression tests in *S. coelicolor*. The temperature of incubation after induction was 25 °C and 30 °C respectively. 20 µL of CP after sonication and centrifugation of 1) *S. coelicolor* CH999_pCJW93_DK1622_CitB at 25 °C; 2) *S. coelicolor* CH999_pCJW93noHis_DK1622_CitB at 25 °C; 3) *S. coelicolor* CH999_pCJW93_DK1622_CitB at 30 °C; 4) *S. coelicolor* CH999_pCJW93noHis_DK1622_CitB at 30 °C; 5) *S. coelicolor* CH999_pCJW93_Cb_vi35_CitB at 25 °C; 6) *S. coelicolor* CH999_pCJW93noHis_Cb_vi35_CitB at 25 °C; 7) *S. coelicolor* CH999_pCJW93_Cb vi35_CitB at 30 °C; 8) *S. coelicolor* CH999_pCJW93noHis_Cb_vi35_citB at 30 °C; 9) *S. coelicolor* CH999_pCJW93_MCy9171_CitB at 25 °C; 10) *S. coelicolor* CH999_pCJW93noHis_MCy9171_CitB at 25 °C; 11) *S. coelicolor* CH999_pCJW93-MCy9171_CitB at 30 °C; 12) *S. coelicolor* CH999_pCJW93noHis_MCy9171 at 30 °C 13) Supernatant (SN) of *S. coelicolor* CH999_pCJW93-MCy9171_CitB at 25 °C.

In Fig. S15 and Fig. S16 the band of the expected protein is visible in the CP and in the SN, revealing an overexpression of $citB_{MCy9171}$. The amount of protein depends on the composition of lysis buffer, used to extract the protein for further purification steps. For that reason, different lysis buffers were used to solute the protein after mechanic cell lysis. To test the correlation of protein purification and different lysis buffers, 20 mL of TSB medium was inoculated with spores of *S. coelicolor* CH999 harboring the constructed shuttle vector pCJW93_MCy9171_CitB from a glycerol (20%) cryogenic long-term stock to cultivate a seed culture. After three days, the culture was well-grown and suitable to inoculate two times 100 mL of Super YEME media containing 50 µg/mL apramycin in a 250 mL shake flask with metal spiral. After four days, the cultures were induced with either 10 or 20 µg/mL thiostrepton. The cultures were incubated for 24 h at 25 °C with 180 rpm and after incubation 10 mL was transferred in a Falcon tube, centrifuged at 4000 rpm, for 10 min at 4 °C, whereas the SN was discarded. The CPs were re-suspended in lysis buffer No. 3, No. 4 (NaCl 500 mM, Tris pH 8.0 20 mM, imidazole pH 8.0 20 mM) and 12 (NaCl 500 mM, Bis-Tris pH 6.8 20 mM, imidazole pH 8.0 20 mM, glycerol 10% m/m). After sonication and centrifugation at 15000 rpm for 10 min at 4 °C, nickel pulldown was performed. The SDS-PAGE revealed that lysis buffer No. 3 and 12 with a thiostrepton concentration of 10 µg/mL to induce gene expression in the fermentation culture is optimal in order to obtain the highest amount of recombinantly expressed protein (Fig. S17).

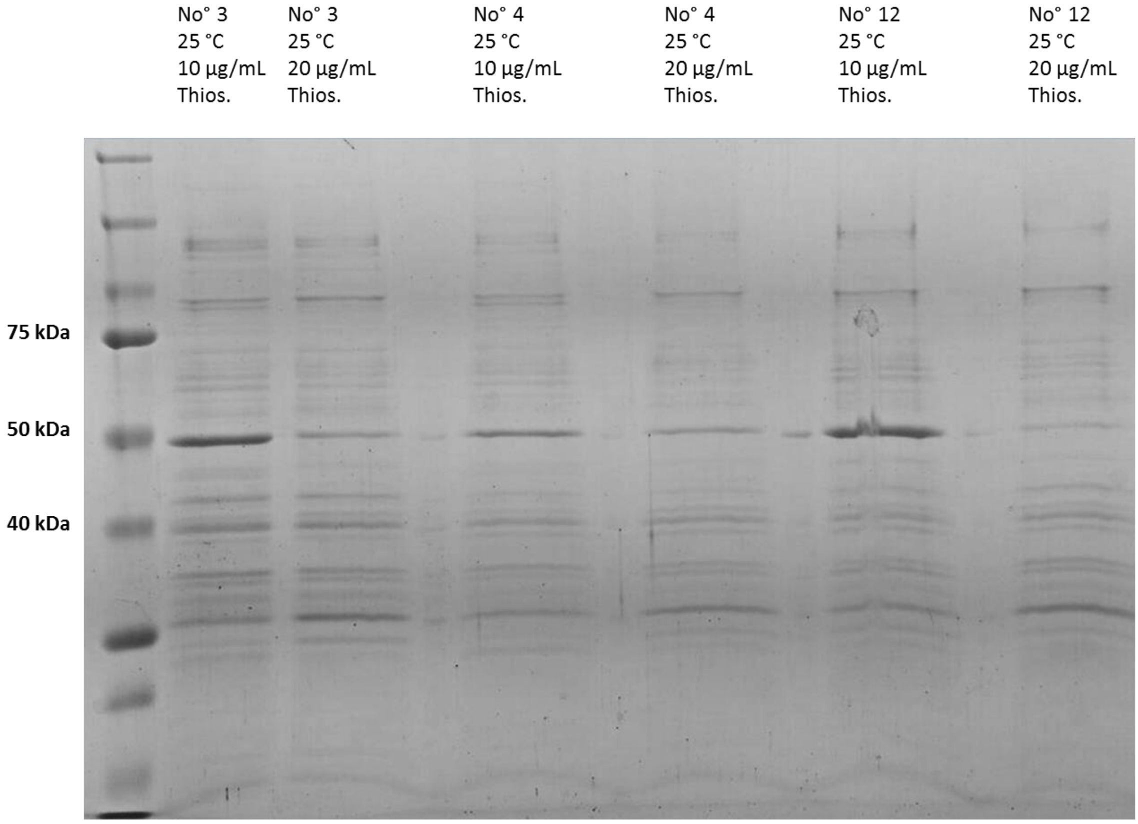

**Fig. S17:** SDS-PAGE of *citB*$_{MCy9171}$ homolog expression tests from *Cystobacter* MCy9171. Each band illustrates the use of different lysis buffers and thiostrepton concentration for induced gene expression. Lysis buffers No. 3 and 12, with a thiostrepton concentration of 10 µg/mL showed the highest yield of recombinant protein.

##### ***Streptomyces*-based recombinant production of $CitB_{MCy9171}$ from *Cystobacter* MCy9171**

The conclusions of the conducted test-expression experiments of the *$citB_{MCy9171}$* homolog from *Cystobacter* MCy9171 were directly implemented for targeted protein purification in larger scale. 20 mL of TSB medium was inoculated with spores of *S. coelicolor* CH999 harboring the constructed shuttle vector pCJW93_MCy9171_CitB from a glycerol (20%) cryogenic long-term stock to cultivate a seed culture. After three days, the seed culture was used to inoculate with 20 mL, 100 mL TSB medium as pre-culture. After two days, this pre-culture was densely grown and used to inoculate with 5 mL, 18 x 100 mL of Super YEME media incubated at 30 °C and 180 rpm for at least two days. The cultures were prepared in 250 mL Erlenmeyer flasks with metal spirals to ensure cell growth in suspension. After two days, the cultures were well grown and gene expression was induced with thiostrepton (working concentration 10 µg/mL). The cultures were incubated at 25 °C, 180 rpm after thiostrepton induction. Twenty-four hours later, the incubation was stopped and the cell broth was centrifuged at 4000 rpm for 10 min at 4 °C. After discarding the SN, the CP (27 g) was re-suspended in 100 mL ice-cold lysis buffer No. 3. Two protease inhibitor cocktail tablets (Roche diagnostics) and 10.8 mg of bovine pancreas were added to the cell suspension. Subsequently the cells were lyzed using a CD-017a constant cell disruption system. Cell debris was removed by centrifugation (15 min at 50000 g) and the SN was loaded to a 5 mL HisTrap FF column (GE Healthcare) with 5 mL/min on an ÄKTA™ pure system (GE Healthcare) after the column was equilibrated with 5 CV lysis buffer No. 3. The column loaded with recombinant protein was washed with 30 CV lysis buffer. Elution was performed a linear gradient up to 100%, with 5 CV elution buffer No. 3 at a flow rate of 5 mL/min. Protein-containing fractions, protein identity and purity were assessed by SDS-PAGE analysis (12% acrylamide) and LC-MS. Combined fractions containing $CitB_{MCy9171}$ were concentrated via centrifugal filtration using Amicon Ultra-30 columns (MW 30000 Da, Merck). Size exclusion chromatography was performed using a Superdex 200 Increase prepacked column. After equilibration with 1.2 CV protein buffer 3, the recombinant protein solution was loaded via a 2 mL loading loop. The size excluded protein fractions were identified via SDS-PAGE (Fig. S18). Combined $CitB_{MCy917}$ fractions were concentrated via centrifugal filtration using Amicon Ultra-30 columns (MW 30000 Da, Merck). The concentrated protein solution was adjusted at 100 µM and aliquoted into PCR tubes. Protein concentrations were determined by UV spectroscopy (with ε280 nm values) using Thermo Scientific™ NanoDrop™ 2000/2000c. The aliquots were frozen immediately in liquid nitrogen and stored at -80 °C.

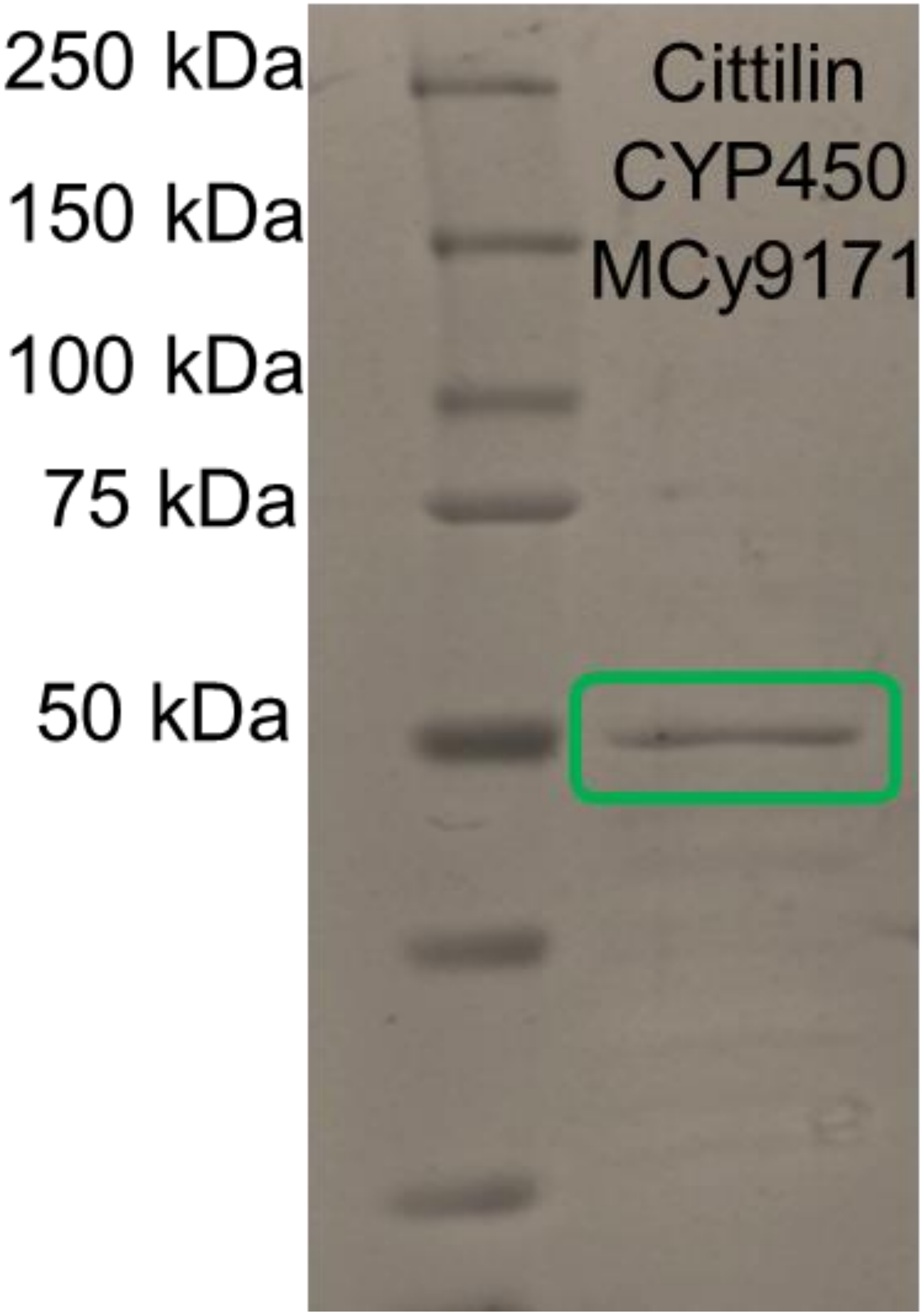

**Fig. S18:** SDS-PAGE gel of $CitB_{MCy9171}$ purification from *Streptomyces coelicolor* CH999. Green box indicate purified $CitB_{MCy9171}$.

#### Catalytic activity testing recombinant $CitB_{MCy917}$ from *Streptomyces* based production

Catalytic activity of the recombinant $CitB_{MCy9171}$ was tested in a reaction mixture containing 4.2 µM $CitB_{MCy9171}$, 125 µM precursor peptide (synthesized and commercially purchased peptide with the sequence KKALYSLAVLMRFARADKLSAPYIYY (DK1622 motif)) or core peptide (synthesized and commercially purchased peptide with the sequence YIYY), 10 µM iron(II) sulfate and iron(III) citrate, 500 µM of co-factor (one of the following: $FAD^+$, $NAD^+$,$NADP^+$, NADPH, NADH) in 500 mM NaCl, 20 mM Bis-TRIS buffer (pH 6.8). As alternative approach to achieve co-factor regeneration, the commercially available Fdx/FdR reductase pair system from *Spinacia oleracea* was performed with 2.5 µM Fdx, 2.5 µM FdR and 2 mM NADPH. Controls were performed by omitting recombinant cittilin cytochrome P450. The reaction was carried out for 10 min, 1 h and (o/n) at room temperature and 30°C. The reaction was terminated by adding MeOH (final concentration 50% v/v). The mixture was transferred to -80 °C for at least 1 h, centrifuged at 13000 g for 15 min at 4 °C (VWR centrifuge ECN521-3601, Hitachi Koki Co., Ltd) and 1 µL of the SN was subjected to HPLC-MS analysis as described previously. As described above catalytic activity of recombinant $CitB_{MCy9171}$ could not be observed.

#### Analysis of recombinant produced CitB by nickel pulldown and SDS-PAGE

10 mL of the liquid cultures of *S. coelicolor* CH999 harboring pCJW93 constructs, which are potential recombinant producers of recombinant CitB were centrifuged for 10 min at 4000 rpm and 4 °C. The CP was re-suspended with 500 µL lysis buffer (500 mM NaCl, 20 mM Bis-Tris pH 6.8, 20 mM imidazole pH 8.0, 10% glycerol, 3 mM β-mercaptoethanol). The mixture was transferred to an 1.5 mL Eppendorf tube and sonicated via "sonics Vibra-cell" (ZinsserAnalytic) with an amplitude of 80% for 15 s on-time and 15 s off-time for a total of 2 min of on-time. After sonication, the tubes were centrifuged for 10 min at 15000 rpm and 4 °C. The SN was transferred to a new 1.5 mL Eppendorf tube and stored on ice. The "KINGFISHER mL"(Thermo Scientific™) was used to perform a nickel pulldown. The principle is based on the use of magnetic nickel beads, which have an affinity to the $His_6$-tag of the recombinant protein. Five hundred µL of cell lysate was mixed with 50 µL of magnetic nickel beads to bind the $His_6$-tagged protein. Afterwards, the mixture was washed twice with 500 µL of lysis buffer, before the protein was eluted with 50 µL of elution buffer (250 mM imidazole pH 8.0). After nickel pulldown, 20 µL of the elution fraction was transferred to a 1.5 mL Eppendorf tube and heated up to 95 °C for 2 min with 4 µL of SDS loading dye (6x). The SDS-PAGE was loaded with 20 µL of the prepared samples; 2 µL of "PageRuler Unstained BroadRange Protein Ladder" (Thermo Fisher Scientific™) was loaded for monitoring the progress of SDS-PAGE and for estimating the approximate size of separated proteins after staining of the gel. The electrophoresis was performed in a "Mini-PROTEAN® Tetra System" (BIO RAD) with SDS (1x) Laemmli buffer with a voltage of 120 V for 90 min. After electrophoresis

was finished, the SDS-PAGE was heated in a microwave with staining solution (Coomassie-blue 0.5%, MeOH 50%, acetic acid 7%, $H_2O$ 43 %) and stored in water for 24 h.

##### *In vitro* enzymatic conversion of $CitB_{MCy9171}$ in cell-free lysate

10 mL of liquid cultures of thiostrepton-induced cultures of *S. coelicolor* CH999 + pCJW93, *S. coelicolor* CH999 + pCJW93_MCy9171_CitB and non-induced *S. coelicolor* CH999 wild type were centrifuged for 10 min at 4000 rpm and 4 °C. The CP was re-suspended with 500 µL lysis buffer (500 mM NaCl, 20 mM Bis-Tris pH 6.8, 10% glycerol (v/v)). The mixture was transferred to an 1.5 mL Eppendorf tube and sonicated via "sonics Vibra- cell" (ZinsserAnalytic) with an amplitude of 80% for 15 s on-time and 15 s off-time for a total of 2 min of on-time. After sonication, the tubes were centrifuged for 10 min at 15000 rpm and 4 °C. The SN was transferred to a new 1.5 mL Eppendorf tube and stored on ice as cell-free lysate.

Catalytic activity of the respective cell-free lysate was tested in a reaction mixture containing 97.5 µL of the freshly prepared cell-free lysate and 62.5 µM precursor peptide (KKALYSLAVLMRFARADKLSAPYIYY (DK1622 motif)) (total reaction volume: 100 µL). Controls were performed by replacing 62.5 µM precursor peptide through 100 µM core peptide (synthesized and commercially purchased peptide with the sequence YIYY). In order to detect the formation of cittilin B, similar reaction set-up as mentioned above was prepared, with additional supplementation of 5 µM recombinantly produced prolyl endopeptidase. The reaction was carried out for 5 h at 30°C. The reaction was terminated by adding MeOH (final concentration 50% v/v). The mixture was transferred to -80 °C for at least 1 h, centrifuged at 13000 g for 15 min at 4 °C (VWR centrifuge ECN521-3601, Hitachi Koki Co., Ltd) and 1 µL of the SN was subjected to HPLC-MS analysis as described previously.

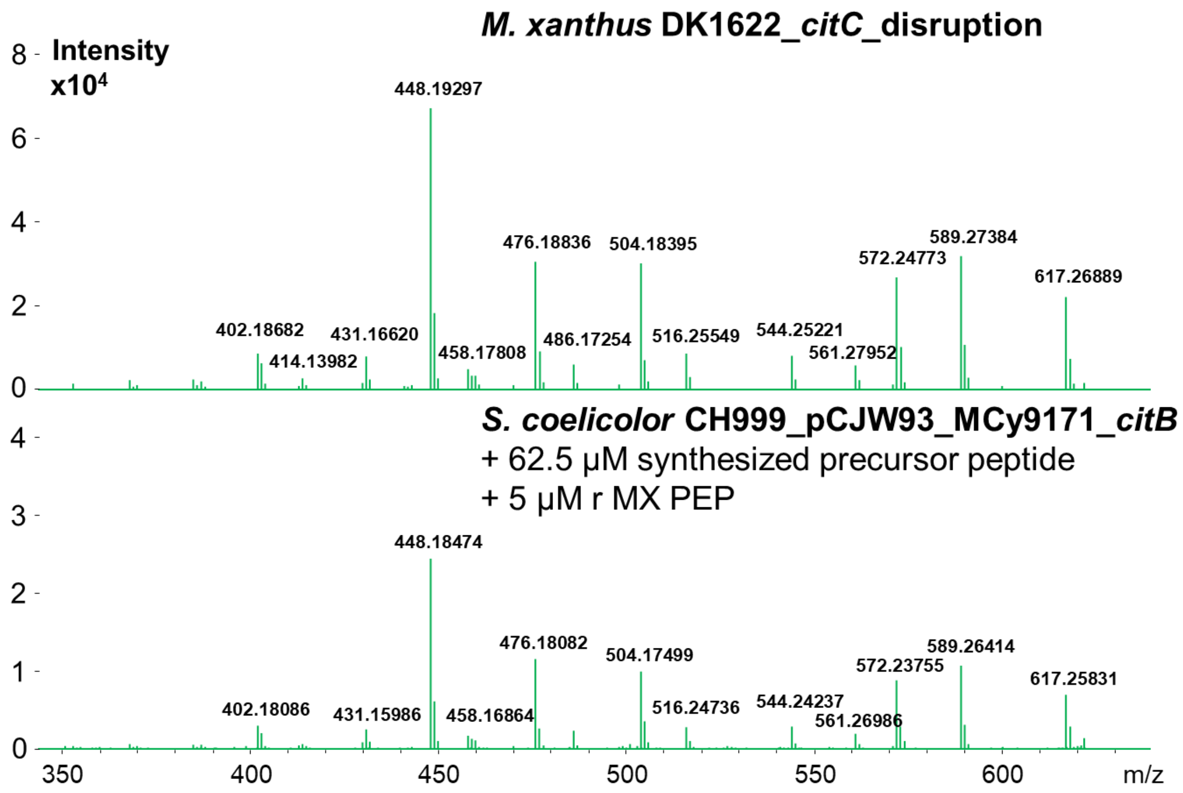

**Fig. S19** MS²-spectra of cittilin B. Cittilin B produced by *M. xanthus* DK1622_*citC*_disruption mutant (**top**) and cell-free lysate of *S. coelicolor* CH999_pCJW93_MCy9171_*citB* supplemented with chemically synthesized precursor peptide (62.5 µM) and recombinantly produced prolyl endopeptidase MX PEP (5 µM) (bottom). Due to the selective MS/MS fragmentation, it was not possible to show a time interval of the MS/MS fragmentation pattern.

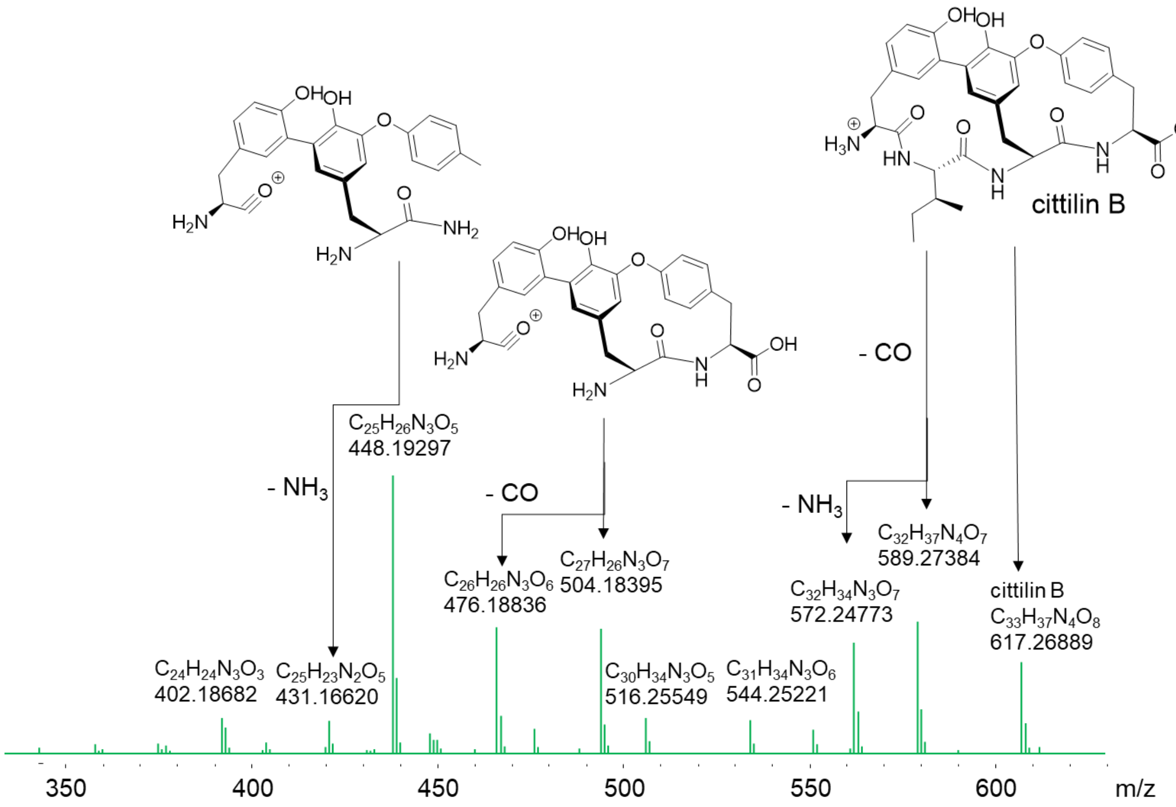

**Fig. S20** MS²-spectrum and fragmentation pattern of cittilin B.

#### CO difference spectra of CitB$_{MCy9171}$

Spectroscopic properties of purified recombinant CitB$_{MCy9171}$ were analyzed using a double-beam spectrophotometer (UV-2101PC, Shimadzu, Japan). Recombinant CitB$_{MCy9171}$ was diluted to 2.5 µM (water or buffer, see above), reduced by using 10 µL of a saturated solution of sodium dithionate and CO was generated by the addition of a small spatula of sodium boranocarbonate. All UV-visible absorbance spectra were recorded from 200 to 700 nm *(46,47)* (data not shown).

### Biological function of cittilin A

#### Cell based bioactivity profiling

##### Antimicrobial assay

Standard sterile microbiological techniques were maintained throughout. All microorganisms were handled according to standard procedures and were obtained from the German Collection of Microorganisms and Cell Cultures (Deutsche Sammlung für Mikroorganismen und Zellkulturen, DSMZ) or were part of our internal strain collection. Cittilin A was tested in microbroth dilution assays on the following panel of microorganisms: *E. coli* DSM-1116, *E. coli* JW0451-2 (*acrB*-efflux pump deletion mutant of *E. coli* BW25113), *Pseudomonas aeruginosa* PA14, *Bacillus subtilis* DSM-10, *Mycobacterium smegmatis* mc2-155, *Staphylococcus aureus* Newman, *Candida albicans* DSM-1665, *Citrobacter freundii* DSM 30039, *Pichia anomala* DSM-6766 and *Acinetobacter baumannii* DSM30007. Microbroth dilution assays were conducted with prepared o/n cultures from cryogenically preserved long-term cultures and were diluted to achieve a final inoculum of $10^4$–$10^5$ cfu/mL. Serial dilutions of compounds were prepared in sterile 96-well plates in the respective test medium. The cell suspension was added and microorganisms were grown for 18–48 h at 37, 30 °C, respectively. Growth inhibition was evaluated by visual inspection and given as minimum inhibitory concentration (MIC) values (the lowest concentration of antibiotic at which no visible growth was observed). No inhibition of one of the tested microorganisms was observed at concentration up to 64 µg/mL of cittilin A.

##### Cytotoxic activity

Cell lines were obtained from the German Collection of Microorganisms and Cell Cultures (Deutsche Sammlung für Mikroorganismen und Zellkulturen, DSMZ) or were part of our internal collection and were cultured under conditions recommended by the depositor. HCT-116 (human colon carcinoma cell line, DSMZ No. ACC 581) and KB-3-1 (cervix carcinoma cell line, DSMZ No. ACC 158) cells were seeded at 6 x $10^3$ cells per well of 96-well plates in 180 µL complete medium and treated with cittilin A in serial dilution after 2 h equilibration. After 5 days incubation, 20 µL of 5 mg/mL MTT (thiazolyl blue tetrazolium bromide) in phosphate buffered saline (PBS) was added per well and it was further incubated for 2 h at 37°C. The medium was discarded and cells were washed with 100 µL PBS before adding 100 µL isopropanol/10 N HCl (250:1) in order to dissolve formazan granules. The absorbance at 570 nm was measured using a microplate reader (Tecan Infinite M200Pro), and cell viability was expressed as percentage relative to the respective MeOH control. $IC_{50}$ values were determined by sigmoidal curve fitting. The $IC_{50}$ of cittilin A against HCT-116 cells was determined to 110.4 µg/mL and against KB-3-1 cells to 74.8 µg/mL.

#### Pyocyanin assay

For determination of extracellular levels of pyocyanin produced by *Pseudomonas aeruginosa* strain PA14, cultivation was performed in the following way: cultures (initial $OD_{600}$ = 0.02) were incubated with or without inhibitor (final DMSO concentration 1%, v/v) at 37 °C, 200 rpm and a humidity of 75% for 16 h in 24-well Greiner BioOne. Cellstar plates containing 1.5 mL of PPGAS medium per well. Pyocyanin produced by PA14 was quantified using the method of Essar et al. *(48)* with some modifications, as described in detail by Klein et al.*(49)*. Briefly, 900 µL of each culture were extracted with 900 µL of $CHCl_3$ and 800 µL of the organic phase re-extracted with 250 µL of 0.2 M HCl. $OD_{520}$ was measured in the aqueous phase using FLUOstar Omega. For each sample, cultivation and sample work-up were performed in triplicates. Inhibition values of pyocyanin formation were normalized to $OD_{600}$. Ten µM of cittilin A, inhibited 3.34% of pyocyanin production.

#### Synthesis and purification of rhodamine-coupled cittilin A

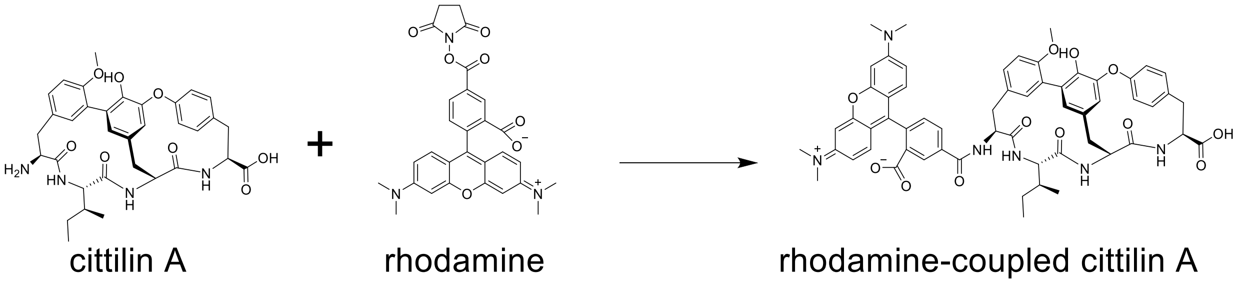

**Fig. S21** Synthesis and purification of rhodamine-coupled cittilin A

The following reaction was carried out under nitrogen atmosphere; To a solution of cittilin A (2.18 mg, 0.0035 mmol) in anhydrous DCM (3.0 mL) with molecular sieves 4 °A, a solution of 5-carboxy-tetramethylrhodamine *N*-succinimidyl ester (2.489 mg, 0.0047 mmol) was added in anhydrous DMF (0.5 mL) followed by *N*,*N*-Diisopropylethylamine (12 µL, 0.691 mmol). The mixture was stirred for six days at 28 °C and monitored by LC-MS. The mixture was dried (o/n) under vacuum to remove DCM and DMF. The residue was re-dissolved in 550 µL bidestilled MeOH and directly subjected to preparative RP-HPLC without further workup. Semi-preparative HPLC purification was done using a Dionex Ultimate 3000 SDLC low pressure gradient system on a XBridge peptide BEH C18 column 138 Å, 4.6 mm × 2500 mm column. Column temperature was stabilized at 45 °C with the eluents $H_2O$ + 0.1% FA as **A** and ACN + 0.1% FA as **B**, at a flow rate of 1.5 mL/min. Detection of rhodamine-coupled cittilin A was facilitated via mass spectrometry on the Agilent 1100 series coupled to the HCT 3D ion trap or with a UV detector on the Dionex 3000 SL systems by UV absorption at 256 nm and 320 nm. The gradient starts with a plateau at 95% **A** for 2 min followed by a ramp to 62% **A** during 6.3 min. Then, **A** content was kept to 62% during 6 min and finally ramped to 5% **A** during 1 min. **A** content is kept at 5% for 1 min and then ramped back to 95% during 30 s. The column was re-equilibrated at 95% **A** for 3 min. The pure bright pink compound was subsequently dried by lyophilization yielding 0.4 mg rhodamine-coupled cittilin A.

#### Bacterial cell entry test

Cell entry of rhodamine-tagged cittilin A, cittilin A and free rhodamine was tested in Gram-negative bacterial cells (TolC efflux deficient *E. coli* mutant, from internal strain collection) by fluorescence microscopy of treated cultures. The bacterial cells were prepared as follows before testing cell entry:

- 5 mL pre-cultures of *E. coli* TolC were used (o/n, incubation at 37 °C, LB medium, $OD_{600}$:~1)
- O/n cultures were centrifuged (5 min, 8000 rpm, 4 °C in 15 mL Falcon tubes)
- SN was discarded, re-suspended CP with 5 mL PBS buffer, repeated centrifugation
- Discarded SN, re-suspended CP with 5 mL PBS buffer, split up re-suspended *E. coli* cells:

**PFA fixation**

Transferred 4 x 500 µL of re-suspended *E. coli* cells (in PBS buffer) to 4 x 2 mL Eppendorf tubes

500 µL of re-suspended *E. coli* cells + 10 µL of rhodamine-coupled cittilin A [1 mg/mL]

500 µL of re-suspended *E. coli* cells + 21.7 µL of free rhodamine [1 mg/mL]

500 µL of re-suspended *E. coli* cells + 16.5 µL of cittilin A [1 mg/mL]

500 µL of re-suspended *E. coli* cells + 10 µL of MeOH [100%]

**LC-MS analytic**

Transferred 4 x 500 µL of re-suspended *E. coli* cells (in PBS buffer) to 4 x 2 mL Eppendorf tubes

500 µL of re-suspended *E. coli* cells + 20 µL of rhodamine-coupled cittilin A [1 mg/mL]

500 µL of re-suspended *E. coli* cells + 43.4 µL of free rhodamine [1 mg/mL]

500 µL of re-suspended *E. coli* cells + 33 µL of cittilin A [1 mg/mL]

500 µL of re-suspended *E. coli* cells + 10 µL of MeOH [100%]

After supplementation of compounds/MeOH, samples were incubated for 30 min, 37 °C, 400 rpm (Eppendorf incubator), light protected. (A: confocal microscopy/ B: LC-MS analytic)

- Cultures were centrifuged (5 min, 8000 rpm, 4 °C)
- A) discarded SN, re-suspended CP with 500 µL PBS buffer
- B) collected SN, re-suspended CP with 500 µL PBS buffer
- Repeated centrifugation (5 min, 8000 rpm, 4 °C)
- A) discarded SN, added 1 mL PBS buffer + 4% PFA, mixed gently, incubation for 20 min, RT
- B) collected SN, re-suspended CP with 500 µL PBS buffer, repeated centrifugation
- B) collected SN and CP. CP and SN were stored at -20 °C.
- A) After 20 min of PFA fixation, mixtures were centrifuged (5 min, 8000 rpm , 4°C)
- A) discarded SN, added 1 mL PBS buffer, re-suspended cells, repeated centrifugation
- A) discarded SN, CP contains mounted cells, which are now accessible for confocal microscopy

#### LC-MS analysis

- Added to SN samples (four samples) 1.7 mL of MeOH and re-dissolved samples via sonication bathing (30 °C) for 5 min
- Added to CP samples (four samples) 0.5 mL MeOH and 0.5 mL acetone, extracted samples via sonication bathing (30 °C) for 5 min
- Centrifuged all samples (eight samples) for at least 10 min, max. speed, 4 °C (all samples in 2 mL Eppendorf tubes)
- Afterwards transferred SN of samples into brown glass vials (1.5 mL), dried samples under $N_2$ flow. Re-dissolved samples in 200 µL MeOH.
- Transferred 60 µL of each sample to 1.5 mL Eppendorf tubes. Centrifuged all samples for at least 10 min, max. speed, 4 °C (all samples in 2 mL Eppendorf tubes). Submitted undiluted samples for maXis 4G measurement as described above (**Fig. S22**).

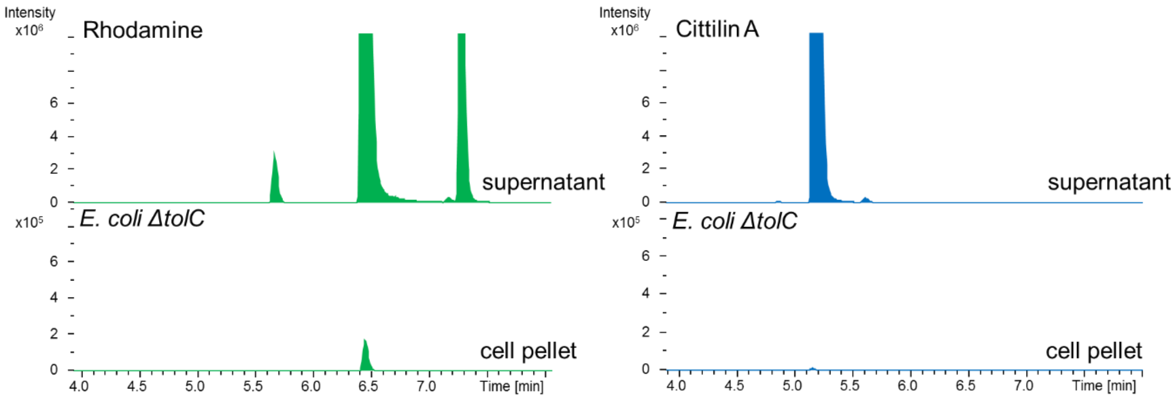

**Fig. S22** Test for bacterial uptake of cittilin A by TolC-deficient *E. coli*. Under the conditions tested, cittilin A is not detected in the bacterial extract. EIC: Extracted ion chromatogram, green: 431.1600 m/z, with a width of 7.9 ppm, free rhodamine [M+H]; blue: 631.2768 m/z, with a width of 7.9 ppm, cittilin A [M+H].

### $^{1}H$ and $^{13}C$ NMR spectra of cittilin A and rhodamine-coupled cittilin A

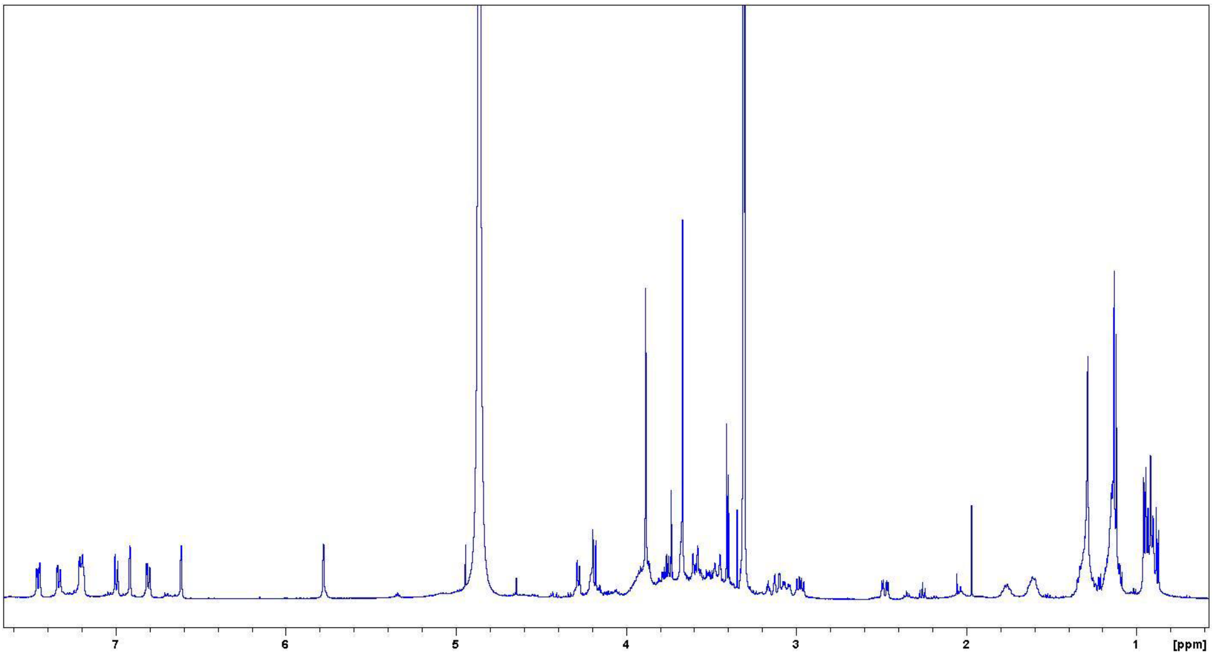

**Fig. S23** $^{1}$*H NMR* spectrum of cittilin A in MeOD.

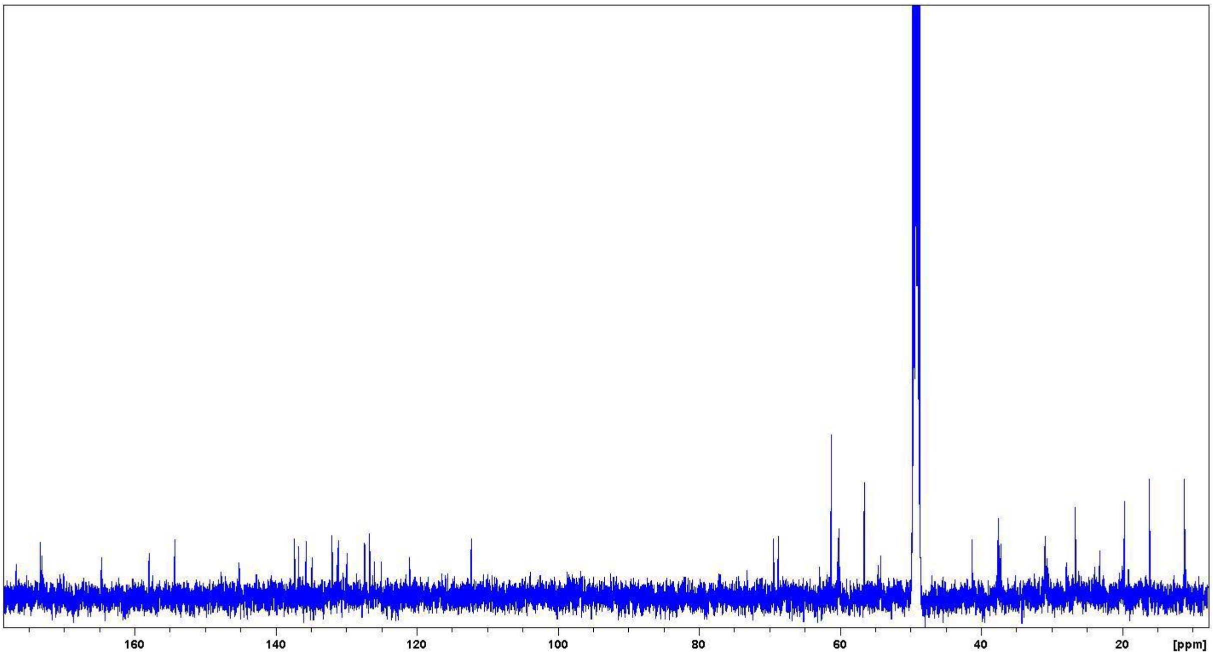

**Fig. S24** $^{13}$*C NMR* spectrum of cittilin A in MeOD.

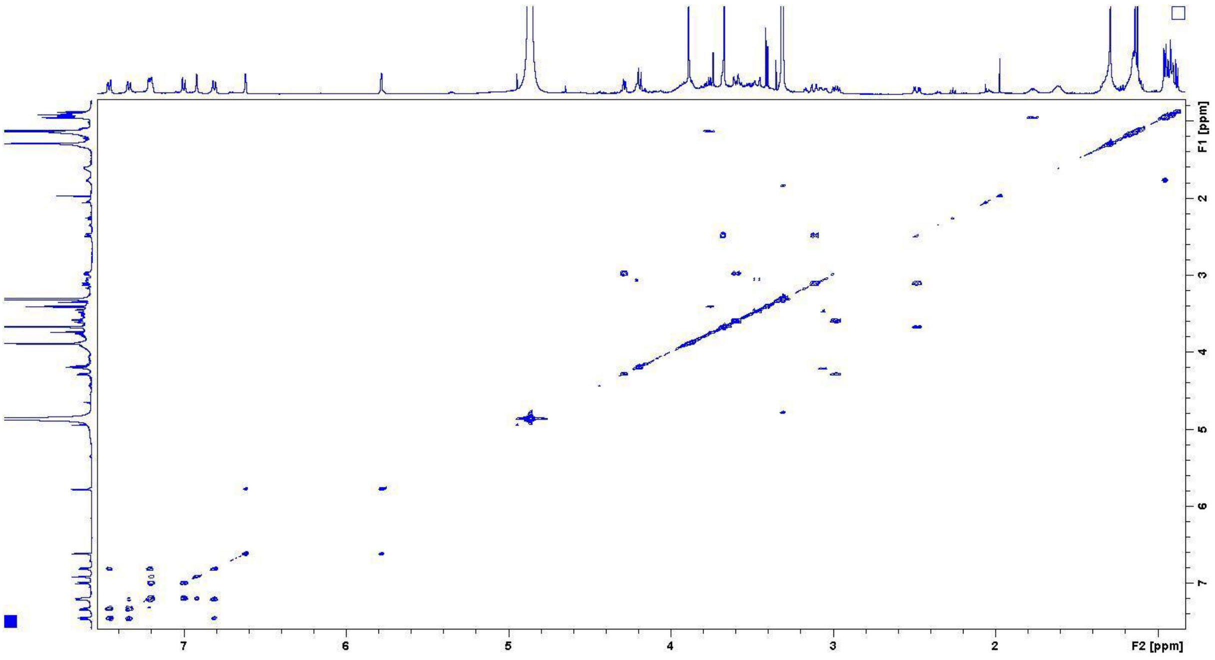

**Fig. S25** *COSY* spectrum of cittilin A in MeOD.

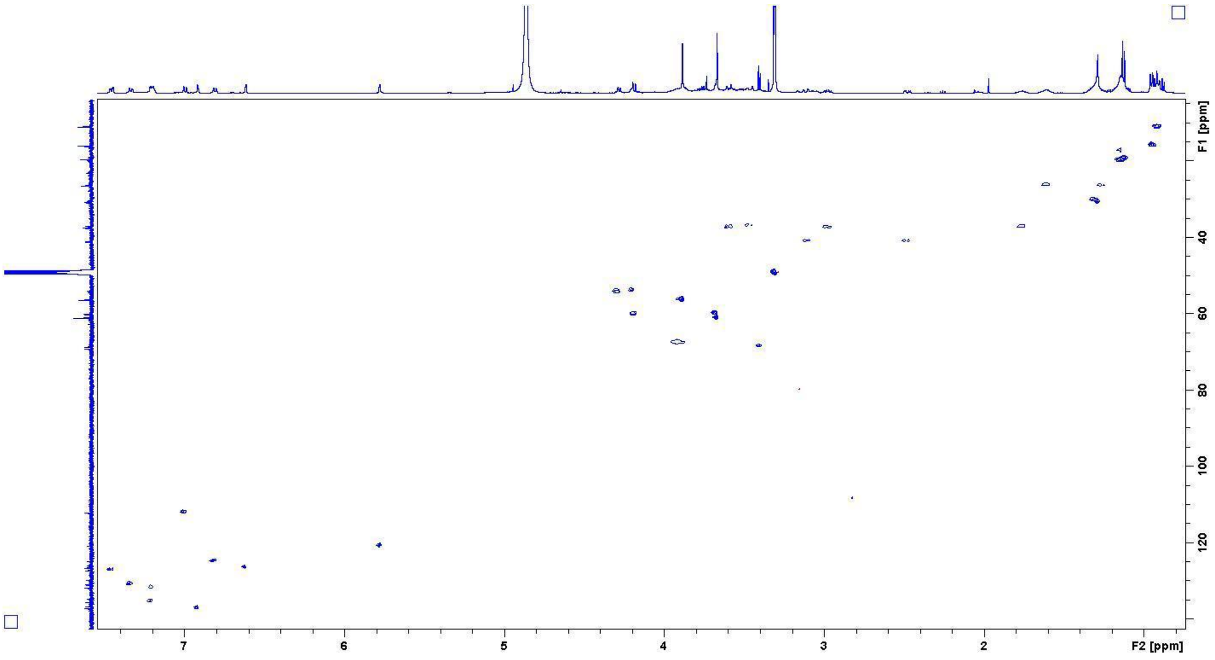

**Fig. S26** *HSQC* spectrum of cittilin A in MeOD.

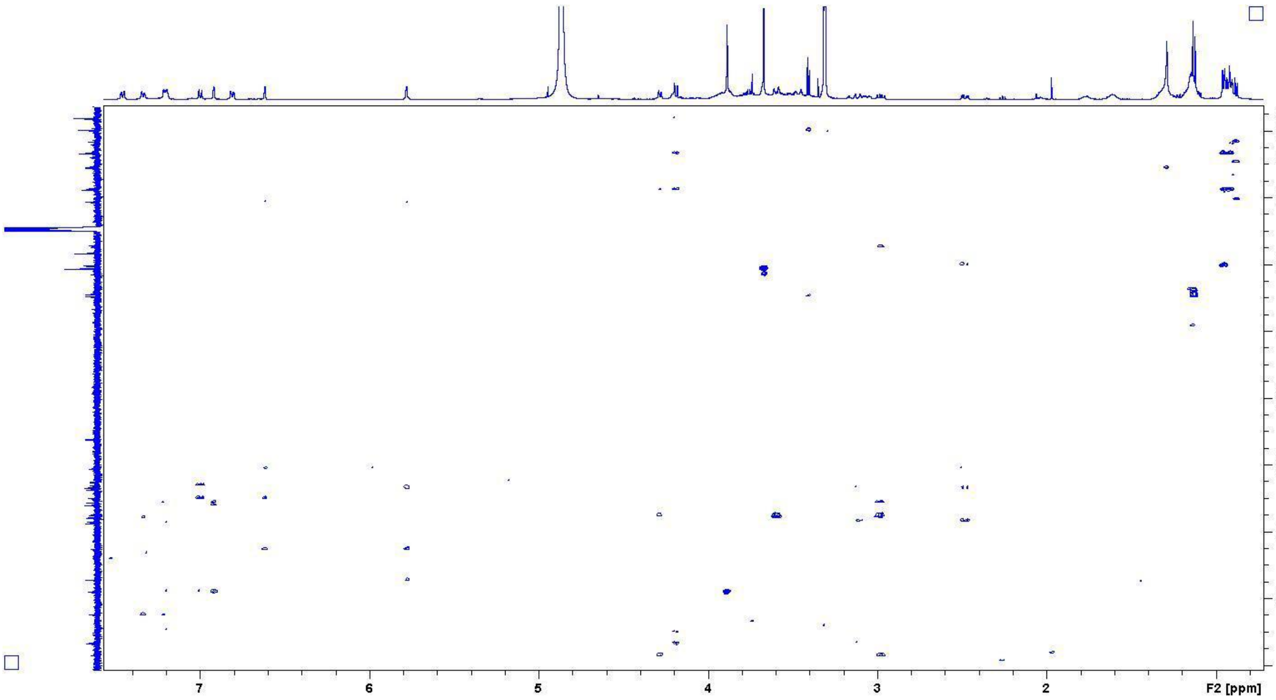

**Fig. S27** *HMBC* spectrum of cittilin A in MeOD.

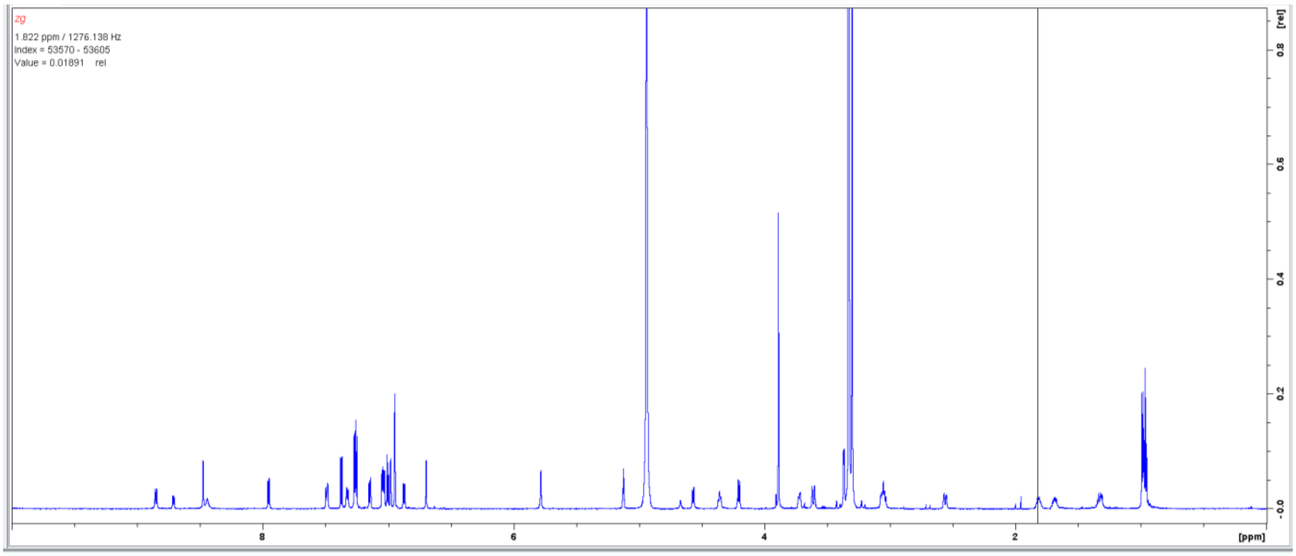

**Fig. S28** $^{1}H$ *NMR* spectrum of rhodamine-coupled cittilin A in MeOD.

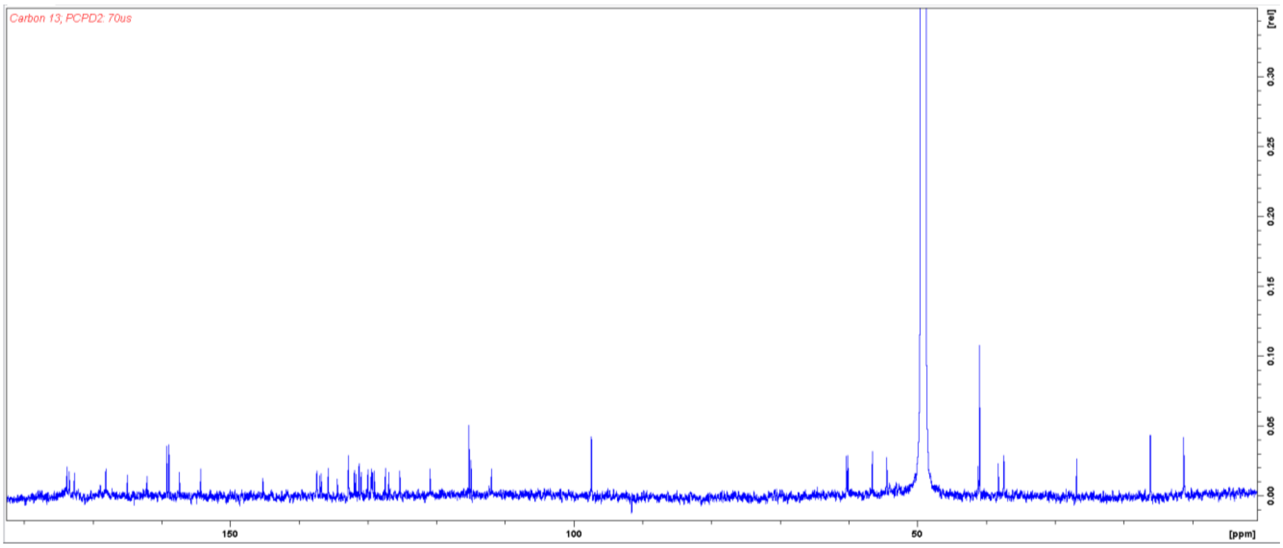

**Fig. S29** $^{13}$*C NMR* spectrum of rhodamine-coupled cittilin A in MeOD.

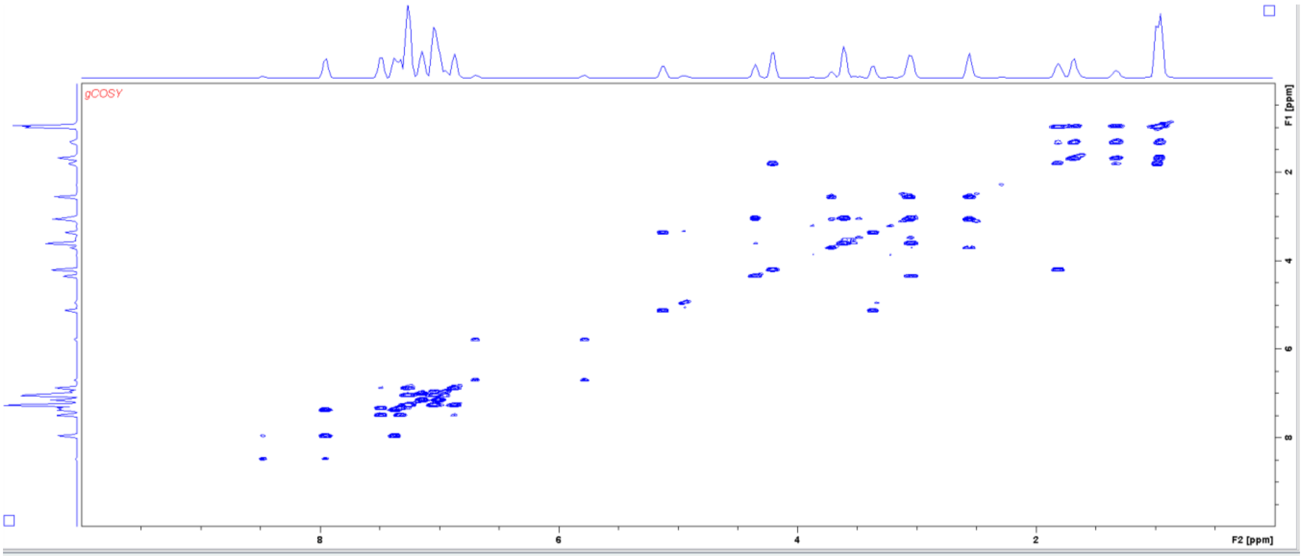

**Fig. S30** *COSY NMR* spectrum of rhodamine-coupled cittilin A in MeOD.

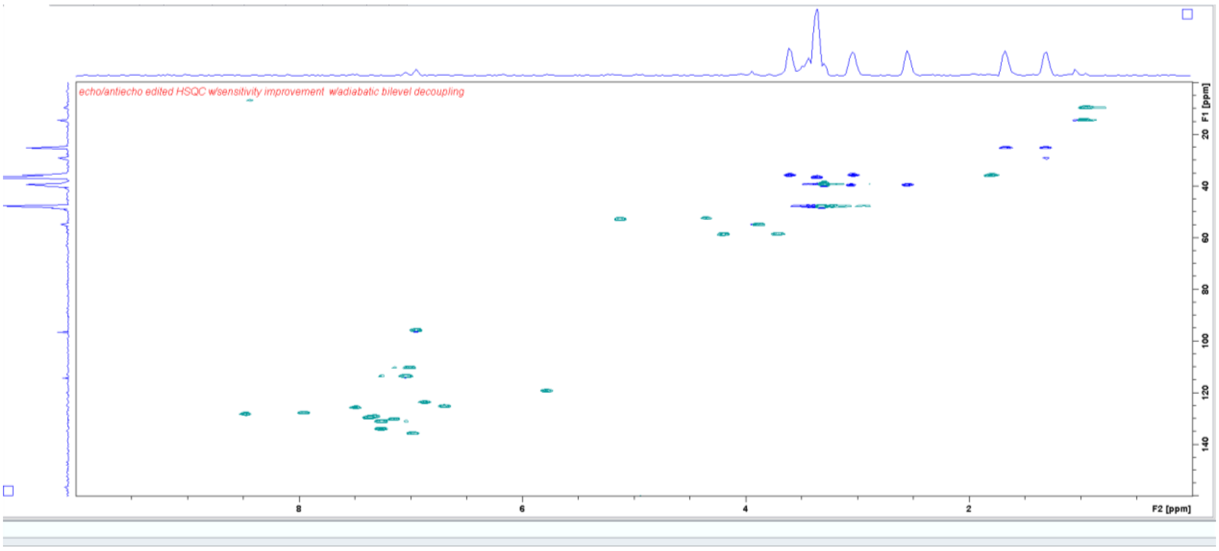

**Fig. S31** *Echo Antiecho* spectrum of rhodamine-coupled cittilin A in MeOD.

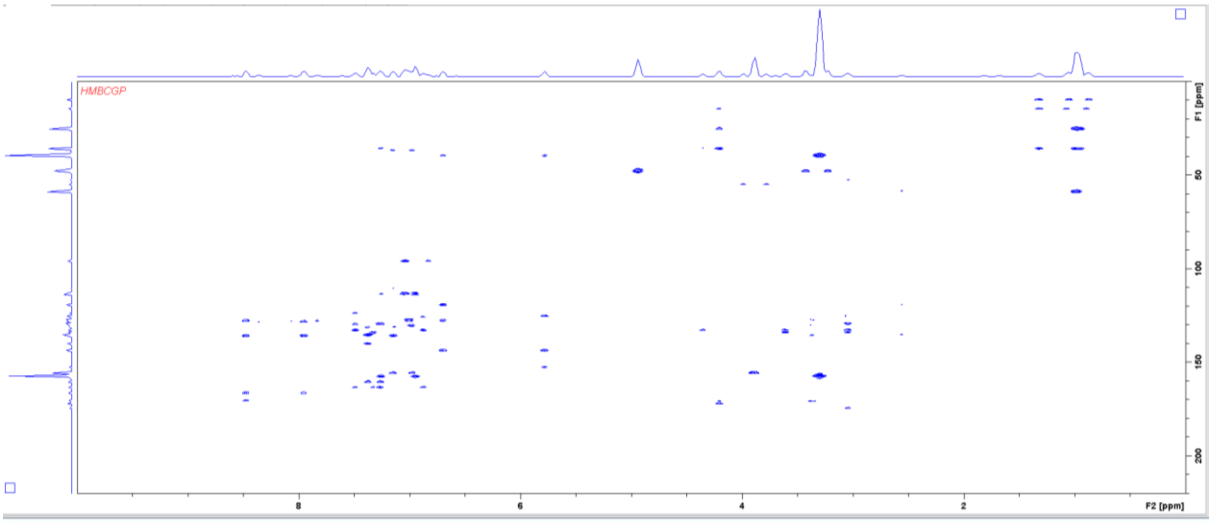

**Fig. S32** *HMBC* of rhodamine-coupled cittilin A in MeOD.
